# A NOD2-Encoded Toggle Switch Resolves the Host–Microbe Battle Over Cyclic AMP Control

**DOI:** 10.64898/2026.03.29.715116

**Authors:** Mahitha Shree Anandachar, David Chen, Kennith Carpio-Perkins, Gajanan D. Katkar, Suchismita Roy, Celia R. Espinoza, Madhubanti Mullick, Saptarshi Sinha, Michelle Nakayama, Jasmin Salem, Megan Estanol, Rishav Bhattacharjee, Kameron Zablan, Shu-Ting Hsu, Sinclair Williams, Annan Sun, Courtney Tindle, Jerry Yang, Irina Kufareva, Pradipta Ghosh

**Affiliations:** Department of Cellular and Molecular Medicine, University of California San Diego, La Jolla, CA 92093, USA; Biomedical Sciences Graduate Program, School of Medicine, University of California San Diego, La Jolla, CA 92093, USA; UC San Diego HUMANOID™ Center, University of California San Diego, La Jolla, CA 92093, USA; Department of Chemistry and Biochemistry, University of California San Diego, La Jolla, CA 92093, USA; Skaggs School of Pharmacy and Pharmaceutical Sciences, University of California at San Diego, La Jolla, CA 92093 USA; Department of Medicine, University of California San Diego, University of California San Diego, La Jolla, CA 92093 USA

## Abstract

Pathogens hijack macrophages by triggering pathological cAMP surges that block phagolysosomal killing—a defect mirrored in phagocytes from refractory colitis. We identify a host-encoded, pathogen-specific surge-protector comprised of a three-protein toggle: The innate sensor NOD2 binds and masks an evolutionarily conserved motif in GIV that activates trimeric-GTPase Gαi, enforcing a biphasic surge-to-plunge cAMP-program: early, NOD2•GIV assembly permits a brief, tolerogenic cAMP rise, whereas subsequent GIV•Gαi engagement collapses cAMP to drive phagolysosomal fusion and microbial clearance. Structural, biochemical, and ultrastructural analyses reveal how molecular toggling imposes precise spatial and temporal control. Pharmacogenomic perturbations pinpoint cAMP–PKA hyperactivation as the defining lesion in GIV-deficient macrophages. Functional studies in primary macrophages and human gut organoid co-cultures show that toggling the NOD2•GIV•Gαi-axis is necessary and sufficient to convert tolerant macrophages into microbicidal machines that preserve mucosal barrier integrity. These findings uncover a druggable cAMP-control pathway with therapeutic promise in colitis.

**GRAPHICAL ABSTRACT:** 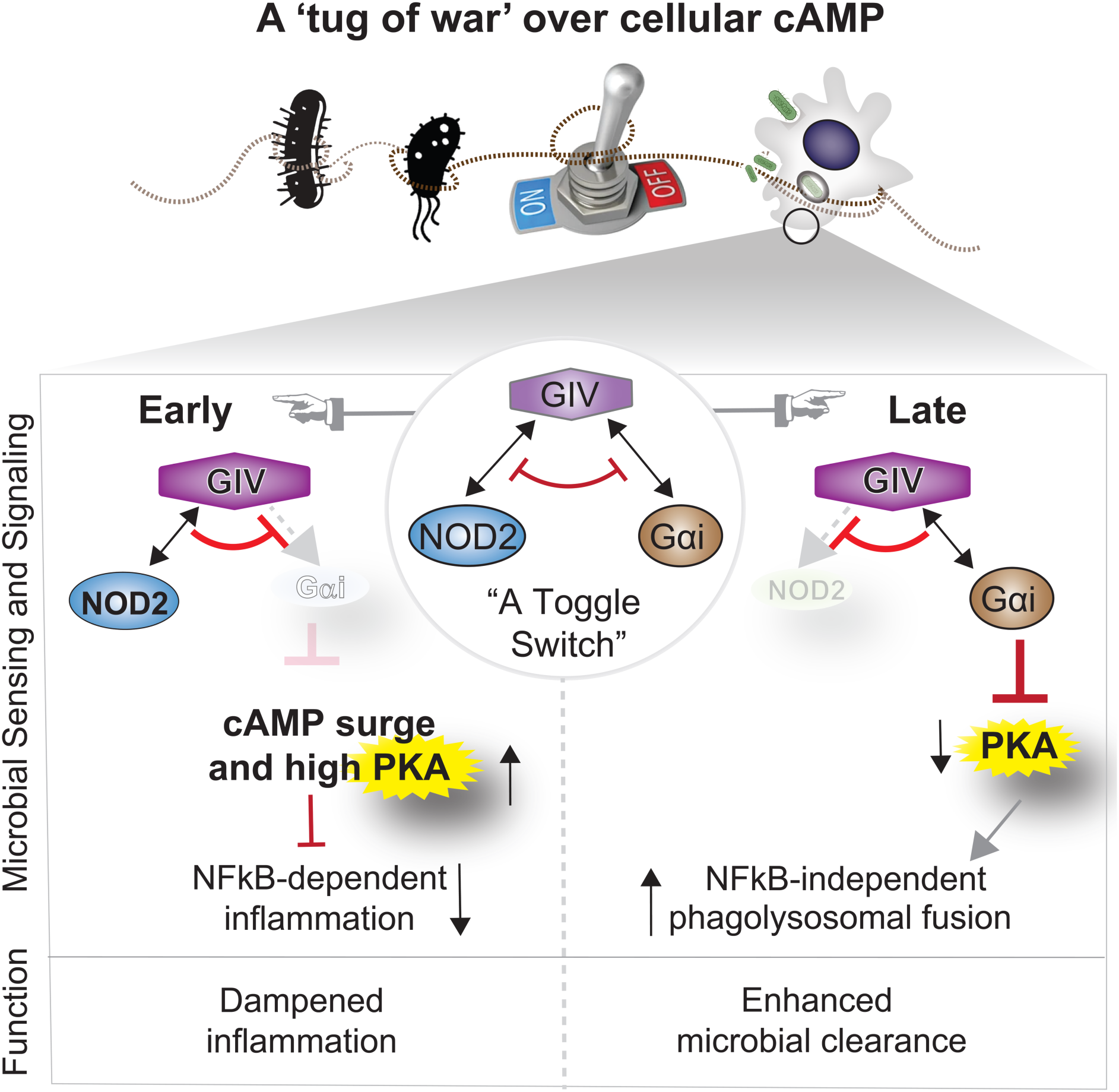

**HIGHLIGHTS:**

- Pathogens elevate macrophage cAMP to evade phagolysosomal killing
- GIV toggles between NOD2 and Gαi to link pathogen-sensing to cAMP control
- Timely cAMP–PKA suppression enables phagolysosomal fusion and pathogen clearance
- NOD2→GIV→Gαi signaling in macrophages protects the intestinal barrier in infection

**eTOC BLURB:** Anandachar et al. identify a host signaling toggle in which NOD2 and Gαi compete for GIV to couple microbial sensing to timed cAMP control. GIV-dependent termination of cAMP–PKA signaling promotes phagolysosomal maturation, pathogen clearance, and intestinal barrier protection

## INTRODUCTION

Half a century ago, the first radioimmunoassays in phagocytes revealed a defining biochemical signature of host defense: intracellular cAMP surges within minutes of microbial encounter, then rapidly returns toward baseline as pathogens are engulfed^1^. Decades later, live-cell FRET imaging confirmed this surge-and-plunge pattern at single-particle resolution^2^, establishing cAMP flux as one of the earliest and most tightly choreographed events in phagocytosis.

The pathophysiological significance of this transient cAMP flux is only partly understood. The early surge, driven by host adenylyl cyclases and amplified by diverse pathogens, tempers macrophage activation, restrains inflammation, and promotes tolerance^3^. Pathogens routinely exploit this window through toxins, efflux pumps, and mimic enzymes that elevate host cAMP and suppress microbicidal programs (summarized in^4^). This widespread hijacking underscores both the evolutionary conservation of an early host-protective signal and its vulnerability. By contrast, the mechanism and significance of the late-phase plunge remain poorly understood. Failure to collapse cAMP impairs phagosomal acidification, lysosomal fusion, and microbial clearance^5^. Accordingly, pathogens that sustain high-cAMP states—including *Mycobacterium*^6–10^*, Salmonella*^11,12^, pathogenic *E. coli*^11,12^ and *Vibrio cholerae*^13^---exploit ‘cAMP intoxication<u>^5^</u>’ to evade clearance, sustain intracellular survival, and promote virulence. Elevated cAMP is also associated with fungal and protozoan infections^4^ and chronic infection-inflammation associated diseases^14^, extending its relevance far beyond acute microbial encounters (**Figure S1A**).

Cellular cAMP s controlled by a multilayered network of adenylyl cyclases, phosphodiesterases (PDEs), and other cAMP-modulating proteins. PDEs are major determinants of the magnitude and duration of cAMP signals^15^ and, together with other regulators, help establish compartmentalized intracellular cAMP pools^16^. Yet how these mechanisms are coupled directly to microbial sensing to produce the rapid, pathogen-proximal collapse of cAMP accompanying phagosome maturation remains incompletely understood. This distinction is important because successful microbial clearance may require not simply restoration of basal cellular cAMP, but termination of cAMP signaling where the pathogen is being processed^17^. Whether phagocytes possess a molecular switch that directly couples pathogen sensing to local cAMP termination therefore remains an important unanswered question.

Clinical evidence further supports the essential role of this host-pathogen standoff in controlling cAMP flux. Patients with sepsis who receive β-blockers exhibit improved survival^18–31^, a benefit whose mechanistic basis remains unclear. By dampening adrenergic drive, β-blockers can reduce cAMP-generating inputs^32,33^, potentially mimicking one aspect of the host response. Thus, whereas microbial mechanisms that accelerate cAMP are well documented, the host-encoded brake that executes the surge-to-plunge transition has remained elusive.

Here, we identify a host-encoded molecular switch that directly couples pathogen sensing to local cAMP termination. The switch centers on GIV (a.k.a., Girdin; *CCDC88A*), a multimodular signal transducer that belongs to the guanine-nucleotide exchange modulator (**GEM**) family^34^ and an established activator of Gαi which suppresses cAMP ^34–36^ (**Figure S1A**). We show that GIV dynamically switches between Gαi and the innate immune sensor NOD2 through mutually exclusive, nucleotide-dependent partner engagement. This partner exchange couples pathogen sensing to Gαi activation and the late cAMP plunge, thereby linking cAMP–PKA control to phagolysosomal maturation and microbial clearance. Together, our findings define a spatially and temporally organized mechanism by which macrophages wrest control of cAMP from invading microbes and determine whether pathogens persist or are cleared.

## RESULTS

### Identification of GIV as a putative candidate that restrains maladaptive cAMP–CREB signals in colitis

Using our previously published, unbiased SMaRT framework [‘*<u>S</u>*ignature of *<u>Ma</u>*crophage *<u>R</u>*eactivity and *<u>T</u>*olerance’ (SMaRT^37^; see **Figure 1A**)], derived from >12,500 human transcriptomic datasets, we defined a conserved macrophage-state continuum^38,39^ and subsequently refined it in healthy and inflamed gut tissues from patients with inflammatory bowel diseases [IBD; representing the two major clinical subtypes, Crohn’s disease (CD) and ulcerative colitis (UC)^40^]. This analysis identified two dominant macrophage programs: **iColAMs** (“accelerators”), tuned for inflammatory signaling, and **niColAMs** (“brakes”), equipped as ‘brakes’ for microbial clearance and resolution^37^.

**Figure 1.**
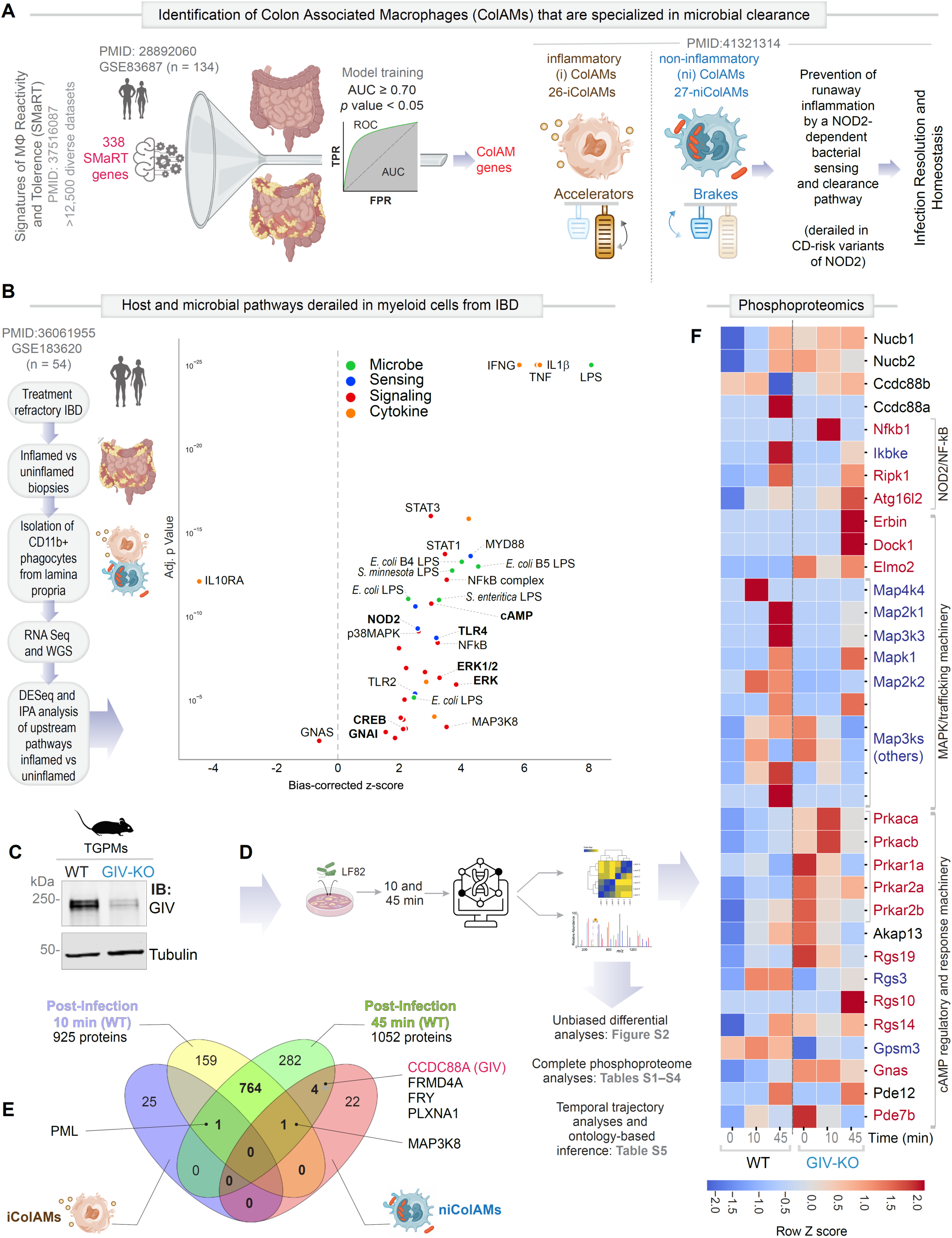
Macrophage *CCDC88A* (GIV) links NOD2 sensing to maladaptive cAMP–PKA signaling in IBD-associated dysbiosis. **A.** A schematic of the SMaRT framework^37^, which resolves colon-associated macrophages (ColAMs) along a continuum into inflammatory ColAMs (iColAMs, “accelerators”) and non-inflammatory ColAMs (niColAMs, “brakes”). niColAMs were previously identified as a specialized macrophage state that supports NOD2-dependent microbial sensing and clearance and is enriched for *CCDC88A* (GIV). **B.** Ingenuity Pathway Analysis (IPA) of lamina propria myeloid cells from patients with treatment-refractory inflammatory bowel diseases (IBD) compared with uninflamed controls in remission highlights host- and microbe-driven upstream pathways^41^, including microbial sensing, inflammatory signaling, cytokine pathways, and cAMP–CREB-associated regulators. Pathways most relevant to this study are labeled. See also **Figure S1A** for a summary of established cAMP-dependent immune sensing and signaling and **Figure S1B** expression of key niColAM genes linking cAMP–CREB signaling to NOD2. **C.** Immunoblot (IB) confirming loss of GIV in thioglycolate-elicited peritoneal macrophages (TGPMs) from *Ccdc88a* (GIV)-KO mice compared with wild-type (WT) littermate controls. Tubulin serves as loading control. **D.** Experimental workflow for time-resolved phosphoproteomic profiling. WT and GIV-KO TGPMs were challenged with *AIEC* LF82 and harvested at baseline and at 10 and 45 min after infection for phosphoproteomic analyses. **E. I**ntersection of the WT infection-induced phosphoproteome with iColAM and niColAM gene sets. Phosphoproteins detected at 10 and/or 45 min after *AIEC* LF82 infection are compared with the transcriptionally defined ColAM programs to identify macrophage-state-associated proteins undergoing dynamic phosphomodulation. Selected intersecting genes are indicated. See also **Figure S2A–B** and **Tables S1–S5** for genotype- and time-resolved analyses and ontology-based interpretation of the phosphoproteomic response. **F.** Heatmap of representative infection-responsive phosphoproteins in WT and GIV-KO macrophages at 0, 10, and 45 min, organized by functional modules. Values represent row-wise Z-scores of protein-level phosphorylation-associated abundances. Red and blue indicate values above and below, respectively, the mean for the same protein across displayed conditions.

GIV was prioritized from these prior unbiased analyses^37^. Briefly, multi-scale analyses—transcriptomic, proteomic, and perturbational—highlighted a specialized bacterial-sensing module within niColAMs^37^ (**Figure 1A**), centered on the innate immune receptor **NOD2** and its physical interactor **GIV** (a.k.a., Girdin, encoded by the *CCDC88A* gene. The latter is a multimodular signal transducer that belongs to the guanine-nucleotide exchange modulator (**GEM**) family and functions as a *bona fide* suppressor of cAMP ^34–36^(**Figure S1A**). Among niColAM-associated genes, GIV emerged as a key determinant of NOD2-dependent restraint of NF-κB signaling in niColAMs. This relationship was validated *in vivo* across three murine models, where myeloid GIV loss impaired inflammatory resolution and microbial clearance; similarly, the IBD-risk variant NOD2-1007fs, which fails to bind GIV, exhibits defective inflammatory control and clearance^40^. These findings motivated our investigation of how the NOD2•GIV interaction controls microbial sensing and clearance.

We next asked whether this previously identified NOD2–GIV axis was represented in human disease states associated with dysregulated microbial sensing and signaling. To provide clinical context rather than *de novo* target discovery, we interrogated transcriptomic profiles of lamina propria phagocytes from patients with IBD stratified by disease type, location, inflammation, and treatment response^41^. Ingenuity Pathway Analysis (IPA) of patients whose disease is refractory to treatments (vs. those in remission) revealed the enrichment of microbial products (e.g., LPS), host microbe sensors (PRRs, NOD2 and TLRs), and downstream cascades (cAMP–CREB, MAPK, NF-κB) and host cytokines (TNF, IL-1β, IFN-γ) (**Figure 1B**). Phagocytes from patients with treatment-refractory IBD showed increased hyperactive CREB signaling, along with a complex network of modulators to finetune it: cAMP (activator), MAPK (activator),^42^ GNAI (inhibitor), STAT1 (antagonist via transcriptional cofactor C/EBPβ competition^43,44^). Conversely, IL10RA and GNAS (cAMP enhancer) were downregulated. This network is known to sustain apoptosis-resistant phagocytes^45^ that suppress TNF production^42,46^, harbor pathogens^45^ and sustain chronic inflammation.

Consistent with this model, *NOD2, CCDC88A, GNAI2,* and *GNAI3*, but not *GNAS*, were higher in phagocytes from treatment-refractory/inflamed IBD than in treatment-responsive/non-inflamed disease; patient-level distributions and exact statistical comparisons are shown in **Figure S1B**. Given that GIV potently suppresses cAMP by activating Gαi and inhibiting Gαs via its conserved C-terminal GEM motif (**Figure S1A**), we hypothesized that NOD2 leverages GIV•Gαi signaling to couple pathogen sensing to cAMP suppression. Because the cAMP–CREB axis can play both protective and pathogenic roles in inflammation^42^ ^5,47,48^ (summarized in **Figure S1A**), such a mechanism would require precise spatiotemporal gating.

Given GIV’s established ability to activate Gαi and suppress cAMP through its conserved GEM motif^34–36,49,50^, these observations led us to hypothesize that NOD2 leverages GIV•Gαi signaling to terminate pathogen-induced cAMP signaling. The human transcriptomic data are associative and do not establish that elevated GIV functionally restrains cAMP–CREB signaling in patient macrophages; rather, they provide the disease context for this hypothesis. We therefore tested the causal relationship experimentally in the studies that follow.

### GIV loss rewires infection-induced phosphosignaling toward sustained innate and cAMP–PKA signaling

To define how GIV shapes early signaling after microbial encounter, we performed time-resolved phosphoproteomic profiling of WT and GIV-KO TGPMs infected with *E. coli AIEC* LF82 (**Figure 1C–D**). Rather than uniformly amplifying signaling, GIV loss produced broad, time-dependent phosphoproteomic rewiring (**Figure 1E-F**; **Figure S2; Tables S1–S5**).

A defining feature of the GIV-KO state was increased phosphorylation-associated signal across innate-sensing and cAMP–PKA modules, including NOD2-associated proteins (Atg16l2, Erbin and Ripk1); NF-κB pathway components; and multiple PKA subunits (notably R1α, which primes PKA catalytic activity^51^) (**Figure 1F**). Gnas, Pde7B and multiple RGS-family proteins were also altered, pointing to broader remodeling of cAMP-control machinery. In contrast, selected MAPK pathway components showed lower relative phosphorylation-associated signal in GIV-KO macrophages, consistent with diversion away from a coordinated MAPK-dependent cytoskeletal and trafficking program rather than uniform amplification of infection-induced signaling (**Figure 1F**).

Unbiased genotype comparisons at 10 and 45 min confirmed distinct WT- and GIV-KO-enriched phosphoprotein sets (**Figure S2A–B**; **Tables S1–S4**). Their temporal organization (summarized in **Table S5**) suggested that WT macrophages transition from small-GTPase/cytoskeletal signaling toward endosomal trafficking and phagosome maturation, whereas GIV-KO macrophages show early dysregulated signaling followed by persistent PKA–Rap and phagolysosomal signaling.

This trajectory generated two experimentally testable predictions that organize the remainder of the study: failure to terminate cAMP–PKA signaling and failure to complete maturation of pathogen-containing compartments. Other GEM-family members (Ccdc88b, Nucb1/2) lacked notable phosphoproteomic shifts, arguing against compensatory remodeling from related GEMs, each of which is known to act in highly context-specialized ways^34,52^.

Together, the data position GIV as an organizer of the signaling-state transition that couples pathogen sensing to cAMP control and antimicrobial trafficking, consistent with our prior finding that myeloid GIV loss impairs microbial clearance and promotes dysbiosis^40^.

### Live microbes amplify, and GIV shapes, biphasic cAMP–CREB dynamics during pathogen clearance

We next asked whether GIV suppresses cAMP–CREB signaling during infection and whether this suppression is required for pathogen clearance. In THP1-derived macrophages (**Figure 2A-B**) loss of GIV impaired clearance of *AIEC* LF82 (**Figure 2C**) without affecting uptake/engulfment (**Figure S3A-B**) and similarly compromised clearance of multiple enteropathogenic strains (**Figure 2C**). In contrast, commensal *E. coli* (K12 and HS) and non-adapted lab strains (DH5α) were largely unaffected (**Figure 2D**), indicating selective vulnerability to pathogenic microbes.

**Figure 2:**
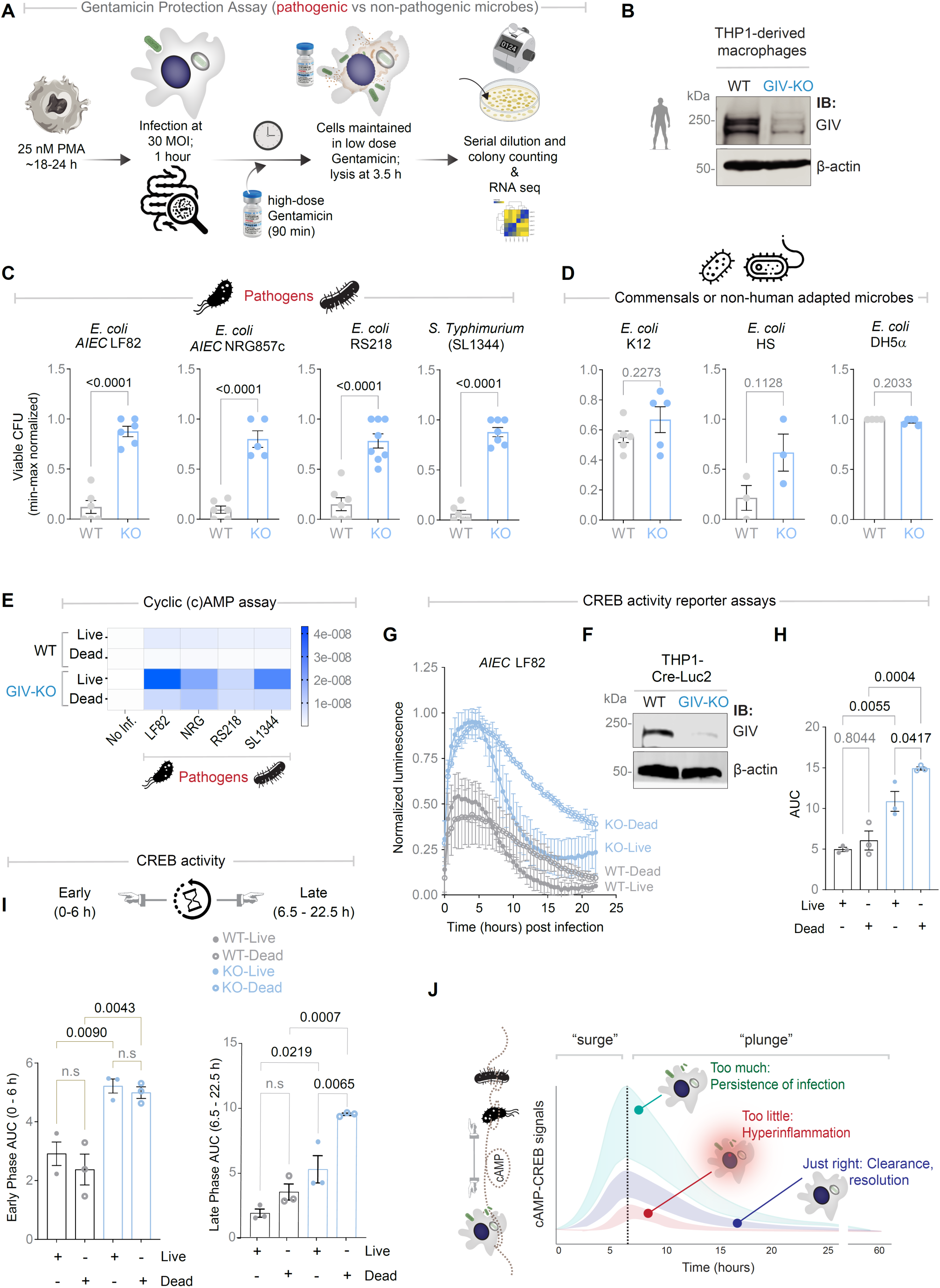
Live microbes and GIV cooperate to shape biphasic cAMP–CREB responses in macrophages. **A**. Workflow for the gentamicin protection assay used to assess intracellular bacterial clearance. **B**. Immunoblot (IB) confirming loss of GIV in GIV-knockout (GIV-KO) compared with control (WT) THP1-derived macrophages. β-actin serves as loading control. **C-D.** Viable intracellular bacterial counts in WT and GIV-KO THP1 macrophages infected with (C) pathogens (*E. coli AIEC* LF82, *E. coli AIEC* NRG857c, *E. coli* RS218, *S. Typhimurium* SL1344) or (D) commensals/non–human-adapted microbes (*E. coli* K12, HS, DH5α). See also **Figure S3A-B** for microbial uptake assay of *AIEC* LF82 by WT and GIV-KO THP1 macrophages. **E.** ELISA-based cAMP assay in control (WT) or GIV-knockout (GIV-KO) THP1 macrophages infected with either live or heat-inactivated (dead) *E. coli AIEC* LF82, *AIEC* NRG857c, RS218 and *S. Typhimurium* SL1344. **F-I**. Immunoblot (IB; F) confirming GIV depletion in WT and GIV-KO THP1-Cre Luc2 macrophages used in G-I. Line graphs (**G**) display time course of CREB-luciferase activity in WT and GIV-KO reporter macrophages infected with live or heat-inactivated (dead) *AIEC* LF82. Area-under-the-curve (AUC) quantification of the CREB responses shown in G: (H) total AUC, (I) early phase (0–6 h) and late phase (6.5–22.5 h) AUC. See also **Figure S3C-F** for CREB activity profiles stratified by microbial viability and genotype, and **Figure S3G-H** for analyses of the late plunge relative to the early surge and comparisons of live versus dead *AIEC* LF82. **J**. Schematic of biphasic cAMP–CREB dynamics during infection, comprising an early “surge” followed by a late “plunge,” and the associated macrophage response states. The full late ‘plunge’ is most pronounced with live microbes in GIV-proficient macrophages. Failure to appropriately terminate cAMP signaling during the late phase is associated with microbial persistence, whereas insufficient signaling is associated with hyperinflammation; appropriately timed surge-to-plunge dynamics favor clearance and resolution. *Statistics*: Data are represented as mean ± SEM (n = 3 independent biological replicates). Statistical significance was tested using an unpaired *t*-test for panels C–D, and one-way ANOVA followed by Tukey’s multiple-comparisons test for panel H and I. Exact p values are indicated in the corresponding panels. *p*-value ≤ 0.05 is considered as significant. ns, not significant.

Direct cyclic (c)AMP measurements revealed exaggerated surges in GIV-deficient macrophages across all pathogens (**Figure 2E**). However, the magnitude of this early rise did not distinguish the critical defect identified by the time-resolved studies below: failure to terminate cAMP signaling during the late phase. Live bacteria induced larger surges than their heat-killed counterparts, reflecting combined microbial and host contributions. Importantly, heat-inactivated bacteria still produced a partial cAMP response. This is likely because they retain structurally preserved microbial ligands, including peptidoglycan-derived NOD2 agonists and other pathogen-associated molecular patterns capable of activating pattern-recognition receptors. GIV’s presence counteracted both sources: it abolished host-induced surges elicited by dead microbes (heat-inactivated; **Figure 2E**), and blunted pathogen-induced surges elicited by live microbes (**Figure 2E**).

To resolve the temporal structure of these responses that are typically masked by the obligatory requirement of PDE inhibitors for cAMP assays (**Figure 2E**), we conducted time series studies in pathogen challenged WT and GIV-KO THP1 macrophages carrying a CREB-luciferase reporter (**Figure 2F**) as a surrogate readout of cAMP dynamics. In presence of GIV, live pathogens triggered a characteristic biphasic response in WT cells: an early surge (within 0-6 hours), followed by sustained suppression lasting 24 hours (**Figure 2G–I**; **Figure S3C-D**), consistent with prior reports^1, 2^. The late “plunge” was impaired in GIV-KO macrophages challenged with either live (**Figure 2G-I**; **Figure S3E-F**) or dead (**Figure S3G-H**) microbes, leading to sustained cAMP–CREB signaling and failed clearance.

These results establish that live microbes amplify, and GIV shapes, the biphasic cAMP–CREB response required for efficient pathogen clearance (**Figure 2J**). Together with prior work showing that GIV collaborates with NOD2 to temper NF-κB–driven inflammation (∼0–6 hours^40^), these data reveal how a transient cAMP surge is followed by a late phase in which GIV drives the collapse of cAMP, transforming an initially tolerogenic cAMP program^3,47,53^ into a microbicidal state.

Collectively, these findings show that GIV selectively enables clearance of pathogenic microbes by enforcing a **biphasic** signaling architecture. The early surge primes macrophages for activation and controlled inflammation, whereas the late GIV-dependent suppression is essential for resolution and pathogen-specific killing. Loss of GIV dismantles this architecture, leading to sustained cAMP–CREB signaling, impaired clearance, and prolonged inflammation, indicating that GIV-dependent control of cAMP–CREB is an essential determinant of pathogen-specific macrophage immunity.

### Biphasic NOD2•GIV assembly functions as a molecular timer for GIV•Gαi-dependent cAMP control

We next asked how the biphasic cAMP–CREB response is encoded at the molecular level. Building on our prior observation that GIV bridges TLR4^54^ and NOD2^40^, we noted distinct assembly dynamics: TLR4●GIV complexes exist constitutively at the cell surface and disassemble upon ligand (LPS) engagement, without pathogen-specificity (**Figure 3A**), whereas NOD2●GIV complexes assemble in response to ligand (MDP) engagement from intracellular pathogens (**Figure 3B**). This contrast suggested that the pathogen-specificity and the temporal behavior of NOD2●GIV interactions may function as an intracellular **molecular “timer”** that instructs biphasic, pathogen-specific cAMP–CREB responses (**Figure 3C**).

**Figure 3:**
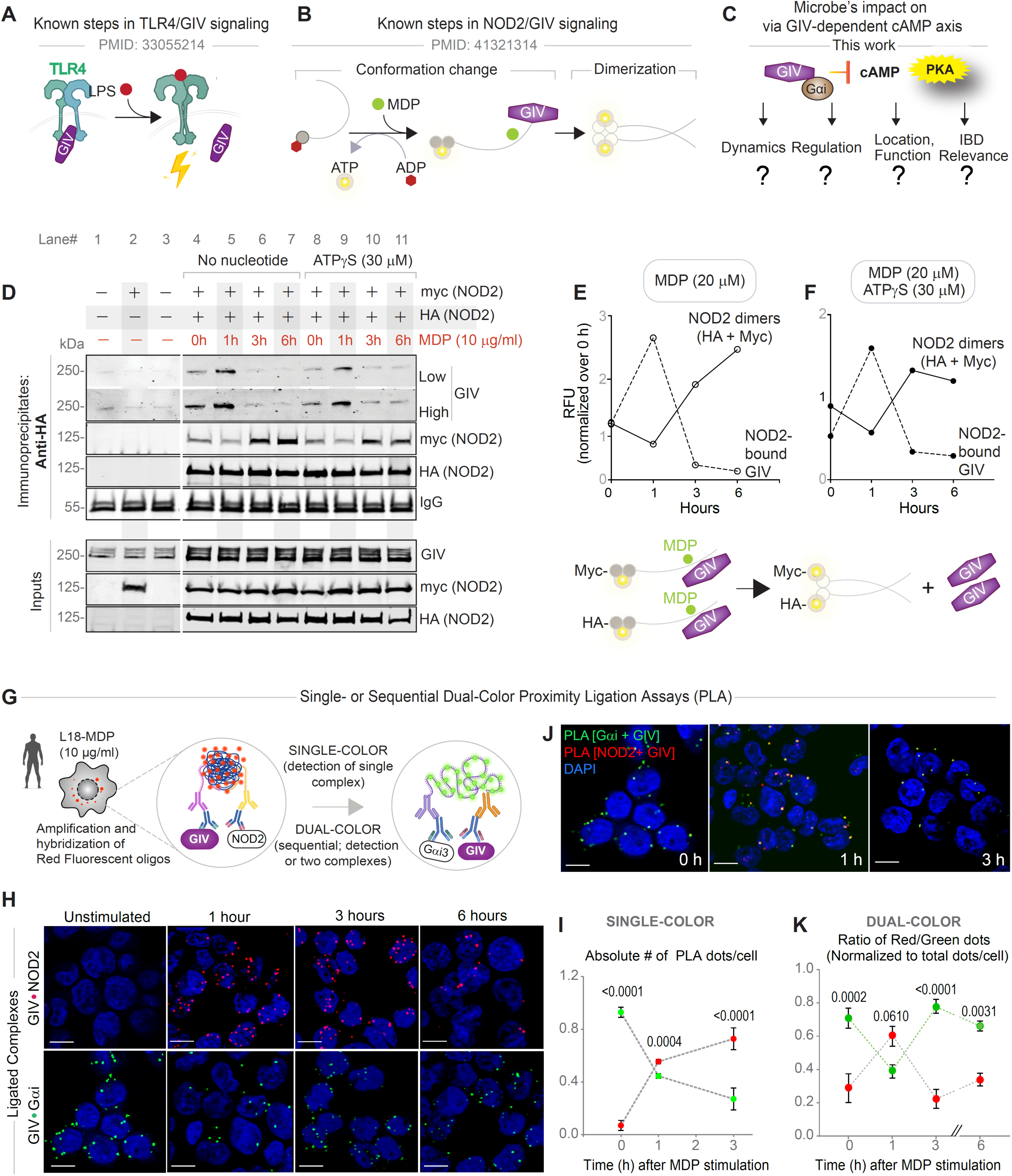
GIV•NOD2 complexes display biphasic assembly and disassembly. **A.** Schematic summarizing canonical LPS•TLR4•GIV signaling^54^. In the basal state, GIV binding to TLR4 prevents receptor dimerization and signaling; LPS stimulation triggers GIV dissociation, permitting TLR4 dimerization and downstream signaling. **B.** Schematic summarizing MDP-induced NOD2•GIV signaling^40^. Resting NOD2 is ADP-bound and autoinhibited; MDP triggers ADP→ATP exchange, conformational ‘opening’ of the LRR module, and NOD2 dimerization with recruitment of downstream partners. **C.** Conceptual framework highlighting this study’s focus on microbial impact via the GIV•Gαi ┤cAMP→PKA axis, which regulates cAMP and PKA suppression in IBD and lists the knowledge gaps (“?”). **D.** Coimmunoprecipitation (Co-IP) of HA- and myc-tagged NOD2 from HEK293 cells following MDP stimulation (0–6 h), with (+) or without (-) ATPγS pre-incubation. Immunoprecipitates (top) and inputs (bottom) were immunoblotted (IB) for endogenous GIV and tagged NOD2. **E-F.** Quantification of the experiments in D following (E) MDP stimulation or (F) ATPγS pre-incubation followed by MDP stimulation, showing NOD2 dimerization (HA•myc) and NOD2-bound GIV over time. Schematics below illustrate the corresponding NOD2•GIV assembly/disassembly states. **G.** Schematic of single-color and sequential dual-color proximity ligation assays (PLA) used to detect endogenous GIV•NOD2 and GIV•Gαi complexes in THP1-derived macrophages. **H-I**. Representative confocal images (H) of single-color PLA following MDP stimulation (0–6 h), showing GIV•NOD2 complexes (red, top) and GIV•Gαi complexes (green, bottom). Nuclei are stained with DAPI (blue). (I) Quantification of PLA puncta per cell from 0–3 h after MDP stimulation. Scale bar: 10 μm **J-K.** Representative confocal images (J) of sequential dual-color PLA following MDP stimulation, simultaneously detecting GIV•NOD2 (red) and GIV•Gαi (green) complexes; nuclei are stained with DAPI (blue). See **Figure S4A** for 3D surface reconstruction and line-scan analysis. **(K)** Quantification of the red-to-green PLA signal ratio, normalized to total PLA puncta per cell, from 0–6 h after MDP stimulation. Scale bar: 10 μm *Statistics*: Data are presented as mean ± SEM. Representative normalized RFU profiles from n=3 independent biochemical experiment. PLA analyses represent ∼20–30 randomly selected fields from *n* = 4–5 independent experiments. Statistical significance was assessed by two-way ANOVA followed by Tukey’s multiple-comparisons test for panels I and K; Exact p values are indicated in the corresponding panels. *p* ≤ 0.05 was considered significant.

Co-immunoprecipitation assays in cells co-expressing HA- and myc-tagged NOD2 revealed that MDP stimulation rapidly induces NOD2●GIV assembly, followed by timed disassembly (**Figure 3D**, *Lane# 4-7*) . Notably, the disassembly phase coincided with NOD2 dimerization (**Figure 3D**, *Lane #4-7* **and E**). Permeabilized cell assays using the non-hydrolysable ATP analogue, ATPγS (**Figure 3D**, *Lane #8-11)--*which mimics MDP-induced conformational activation, recapitulated this assembly/disassembly cycle, indicating that complex disassembly is associated with the nucleotide-dependent conformational transition accompanying NOD2 self-association and does not require ATP hydrolysis per se (**Figure 3D**, *Lane #8-11*, **Figure 3F**).

*In situ* proximity ligation assays (PLA; **Figure 3G**), which report endogenous protein proximity within ∼40 nm^55^, provided an orthogonal measure of the temporal relationship between NOD2•GIV and GIV•Gαi complexes at an *in situ* resolution and physiologic stoichiometry. Single-color PLA revealed temporally regulated NOD2•GIV proximity that was inversely related to GIV•Gαi PLA abundance (**Figure 3H**-**I**). Sequential dual-color PLA, which resolves both complexes within the same cell, revealed a stepwise partner-exchange pattern (**Figure 3J-K**): Early NOD2●GIV engagement coincides with the cAMP–CREB surge (**Figure 2G-I**). Later, GIV transitions back to form GIV●Gαi complexes, aligning with plunge phase.

Together, these findings reveal a biphasic, ligand- and nucleotide-driven cycle of NOD2•GIV assembly followed by oligomerization-associated disassembly, consistent with a molecular timer for GIV•Gαi-mediated cAMP suppression. If so, NOD2•GIV complexes could enable precise temporal control of macrophage antimicrobial responses via the cAMP-CREB axis.

### GIV colocalizes with NOD2 on hemifusome–MVB microdomains and enforces spatial cAMP control

Having established the temporal cycle of NOD2●GIV assembly, we next examined its spatial organization. Because both NOD2 and GIV are cytosolic pathogen sensors and signal transducers, respectively, we reasoned that their interactions at phagosomal membranes could spatially anchor the biphasic GIV•Gαi–cAMP cycle, coupling signal dynamics to bacterial clearance, and that compartmental pH and/or protein conformation might regulate the temporal dynamics.

Biochemical assays further supported this spatial model. GST pulldown and co-immunoprecipitation experiments demonstrated that NOD2•GIV complex stability is both nucleotide (**Figure 4A; Figure S5A**) and pH dependent (**Figure 4B; Figure S5B**), aligning with the progressive acidification and nucleotide exchange that accompany phagosomal maturation. In fact, dual-color PLA in THP1 macrophages revealed spatial proximity of NOD2•GIV and GIV•Gαi complexes, consistent with signaling microdomains where pH and nucleotide-dependent complex composition dynamically shuffle (**Figure S4A**).

**Figure 4.**
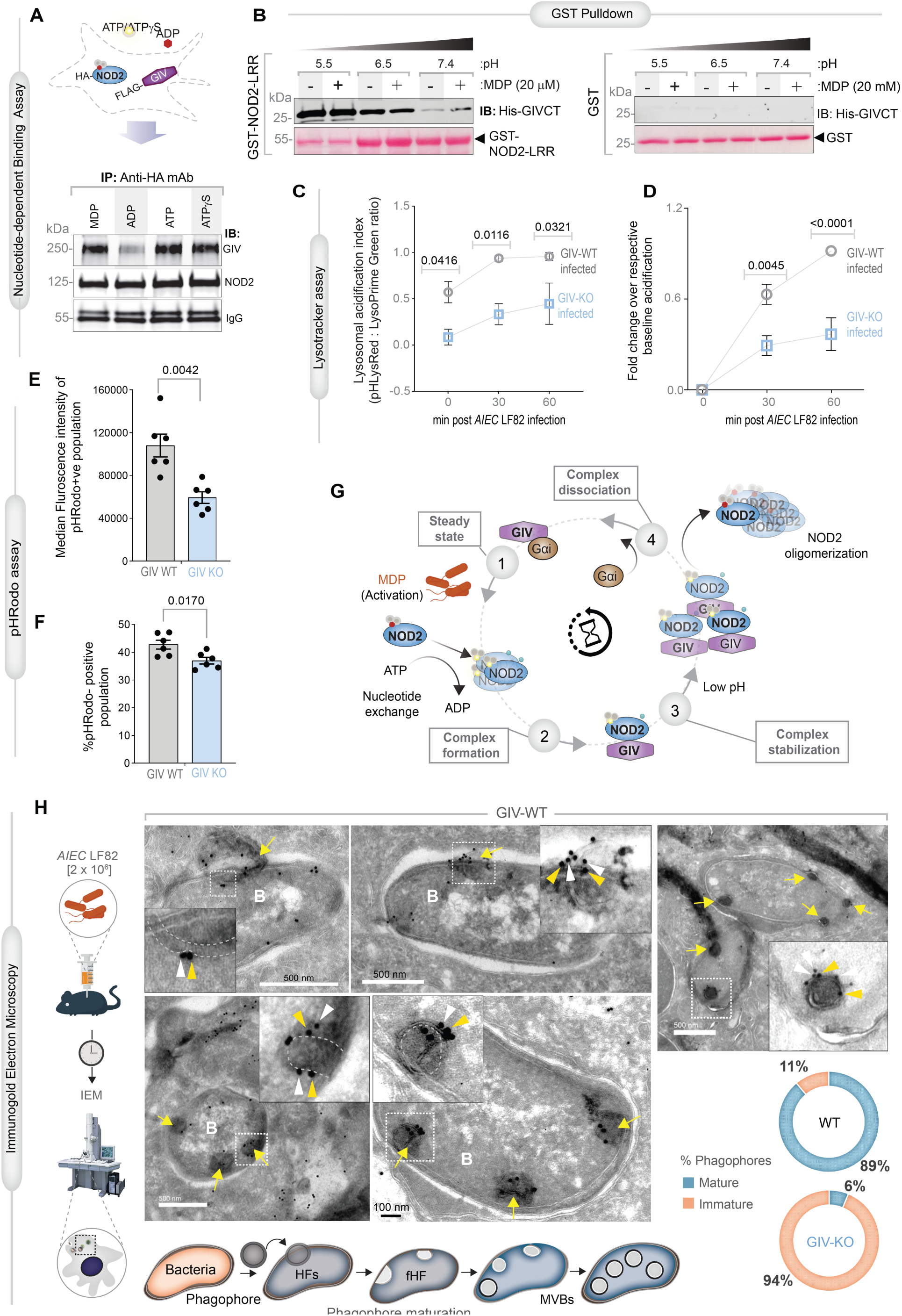
NOD2•GIV complexes couple microbial sensing to phagosome maturation. **A.** Nucleotide-dependent NOD2•GIV binding. HA-tagged NOD2 was immunoprecipitated from HEK293 cells following MDP stimulation or treatment with ADP, ATP, or ATPγS after permeabilization. Immunoprecipitates were immunoblotted (IB) for associated GIV and NOD2. See **Figure S5A** for replicate-level quantification. **B.** pH-dependent NOD2•GIV binding. *Left:* GST pulldown of recombinant His–GIV-CT with GST–NOD2-LRR at the indicated pH (5.5, 6.5, and 7.5), ± MDP. Bound His–GIV-CT was detected by immunoblotting; GST loading was visualized by Ponceau S. *Right:* GST alone serves as negative control. See **Figure S5B** for replicate-level quantification. **C-D**. Lysosomal acidification in GIV-WT and GIV-KO THP1 macrophages following *AIEC* LF82 infection, measured using the dual-fluorescent pHLys Red/LysoPrime Green assay. C. Lysosomal acidification index (pHLys Red/LysoPrime Green ratio) at 0, 30, and 60 min after infection. D. Acidification expressed as fold change relative to the respective 0-min baseline. See **Figure S4B** for representative live-cell fluorescence images and **Figure S4C** for relative lysosomal mass. **E-F.** pHrodo-based assessment of phagosomal acidification in *AIEC* LF82-infected GIV-WT and GIV-KO THP1 macrophages: (E) median fluorescence intensity (MFI) and (F) percentage of pHrodo-positive cells. See **Figure S4D** and **Videos S1–S2** for gating and high-content imaging, respectively. **G**. Mechanistic model of the NOD2•GIV signaling cycle. At steady state, NOD2 is inactive (1). MDP sensing triggers nucleotide exchange and formation of NOD2•GIV complexes (2), which are stabilized at low pH during phagophore maturation (3). NOD2 oligomerization is temporally associated with NOD2•GIV complex dissociation and re-engagement of GIV•Gαi (4). **H.** Immunogold electron microscopy of primary murine peritoneal macrophages exposed to *E. coli AIEC* LF82 (MOI 30, 60 min). NOD2 and GIV were detected using 12-nm (white arrowhead) and 18-nm (yellow arrowhead) gold particles, respectively. Representative images show GIV and NOD2 on phagophore-associated membrane structures (yellow arrows) at progressive stages of maturation, including hemifusomes (HFs), flipped hemifusomes (fHFs), and multivesicular bodies (MVBs); the schematic below summarizes these stages. Donut plots quantify mature and immature phagophores in WT and GIV-KO macrophages, classified by the presence or absence of HFs, fHFs, or MVBs. See **Figure S4E** for representative electron micrographs from GIV-KO macrophages. *Statistics*: Data are presented as mean ± SEM, where applicable. Statistical significance was assessed by two-way ANOVA followed by Tukey’s multiple-comparisons test for panels C–D (*n* =3 independent biological replicates), and unpaired t-test with Welch’s correction for panels E–F (*n* = 3 independent biological replicates). Donut plots represent descriptive percentages of mature and immature phagophores from independent preparations derived from *n* = 4–5 mice per genotype for panel H. Exact *p* values are indicated in the corresponding panels; *p* ≤ 0.05 was considered statistically significant.

The phosphoproteomic trajectory independently predicted that failure to terminate cAMP–PKA signaling in GIV-KO macrophages would be accompanied by defective progression into antimicrobial trafficking (**Table S5**). We therefore tested whether GIV loss impairs maturation of pathogen-containing compartments. We quantified lysosomal acidification using an established fluorescence based dual-probe approach ^56–58^. in which pHLys Red reports lysosomal acidity and LysoPrime Green provides a pH-insensitive measure of relative lysosomal mass. GIV-deficient THP1 cells exhibited reduced lysosomal acidification upon infection (**Figure 4C; Figure S4B**). When normalized to genotype-specific baselines, WT cells showed a markedly greater infection-induced increase in lysosomal acidification at 30 and 60 minutes (**Figure 4D; Figure S4B**). To rule out the possibility that this defect reflects altered lysosomal abundance rather than impaired acidification, we quantified lysosomal mass using LysoPrime Green alone (**Figure S4C**). Lysosomal mass remained comparable between WT and GIV-deficient cells across all time points, indicating that GIV loss does not affect lysosome mass or abundance. Together, these findings indicate that GIV is required for efficient lysosomal acidification in response to bacterial infection.

To further assess phagosomal acidification during *AIEC* LF82 infection, we performed a pHrodo-based assay, in which fluorescence increases as the pHrodo labeled cargo enters acidic phagosomal/phagolysosomal compartments^59,60^. GIV-deficient THP1 cells exhibited significantly lower pHrodo fluorescence compared with WT cells, as reflected by a reduction in median fluorescence intensity (**Figure 4E**). Quantification of the pHrodo-positive population similarly revealed a significant decrease in GIV-deficient cells compared with WT cells (**Figure 4F, Figure S4D**). These findings indicate that the loss of GIV impairs the acidification of bacteria-containing phagosomal/phagolysosomal compartments during *AIEC*-LF82 infection, further supporting a role for GIV in efficient phagolysosomal maturation and acidification.

Integrating the results, a model emerges (**Figure 4G**): MDP binding triggers NOD2 nucleotide exchange and dimerization, enabling GIV binding and formation of NOD2•GIV complexes while suppressing GIV•Gαi interactions, thereby permitting the early cAMP surge. As NOD2 progresses into its oligomerized/activated state, NOD2•GIV complexes dissociate, coincident with re-engagement of GIV•Gαi and cAMP suppression.

As for where in cells these complexes might rearrange dynamically, high resolution immunogold EM of primary peritoneal macrophages infected with live bacteria showed robust co-localization of GIV and NOD2 on phagophore-derived multivesicular bodies (MVBs) across stages of maturation (**Figure 4H**). These included flipped hemifusomes--recently described inverted vesicles that act as ESCRT-independent precursors to MVBs^61^. Hemifusome-associated structures were markedly less frequent in GIV-deficient than WT macrophages (89% WT vs. 6% KO; **Figure 4H, Figure S4E**), consistent with the broader defect in maturation of pathogen-containing compartments observed in GIV-deficient macrophages, without establishing whether their reduced abundance is a cause or consequence of impaired maturation. Because hemifusomes bypass canonical endocytic routes^61^, these findings associate GIV-dependent antimicrobial maturation with non-canonical endocytic routes.

Taken together, the high-resolution imaging places GIV and NOD2 in spatial proximity to hemifusome–MVB microdomains during antimicrobial maturation, without establishing whether GIV directly recruits NOD2 or controls formation of these structures. This localization places GIV and NOD2 near maturing pathogen-containing compartments, providing a spatial context in which GIV•Gαi partner exchange and the late cAMP plunge may occur.

### Structural basis of the molecular toggle: GIV’s GEM motif alternates between NOD2 and Gαi

To understand how GIV switches its binding between NOD2 and Gαi, we turned to structural modeling. Homology models based on resolved structures of rabbit NOD2 (PDB: 5IRN) and Gαi•GIV complex (PDB: 6MHF) revealed a discrete pocket on the 10^th^ LRR of NOD2 that accommodates GIV’s GEM motif (**Figure 5A**); the 10^th^ LRR of NOD2 was recently shown to be critical for binding to GIV^40^. It is noteworthy that both the 10^th^ LRR of NOD2^62^ and GIV’s GEM motif^63^ are known to be evolutionarily conserved. Leu998 (NOD2) within this pocket emerged as critical for binding, whereas Phe996 was dispensable. Notably, the GEM region of GIV, that is required for NOD2 binding, overlapped with the Gαi interface, predicting mutually exclusive interactions (**Figure 5A**)—a structural prerequisite for toggle-like behavior.

**Figure 5:**
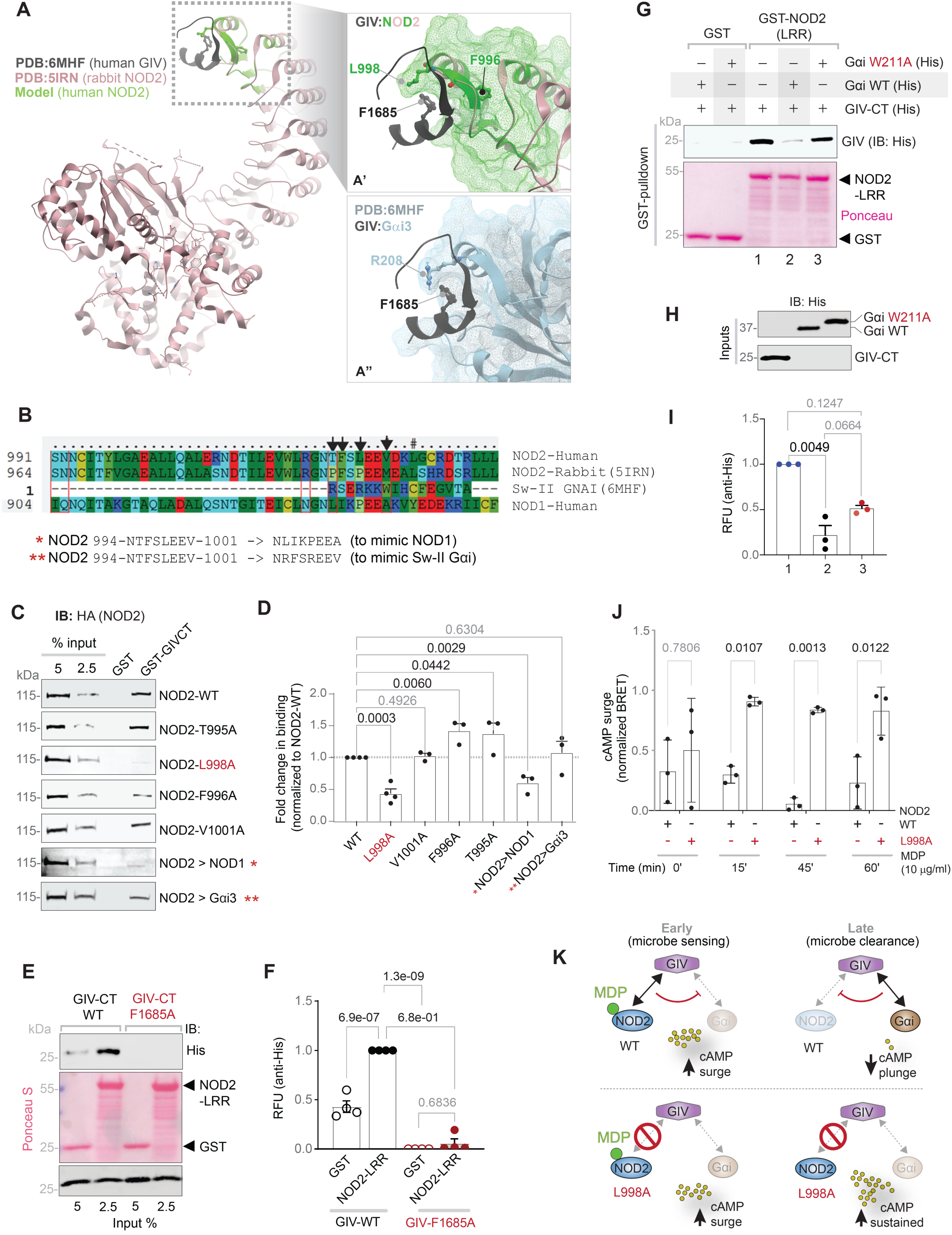
GIV’s GEM motif toggles NOD2 sensing to Gαi-mediated cAMP control. **A.** Structural model of the 10th LRR of human NOD2 (green; based on rabbit NOD2, PDB:5IRN^86^) docked to the GEM motif of GIV (black; PDB:6MHF). A’. Magnified NOD2•GIV interface highlighting NOD2 residues L998 and F996 and GIV residue F1685. L998 appears critical for binding, whereas F996 appears dispensable. A”. Crystal structure of GIV’s GEM motif (black; PDB:6MHF^87^) bound to SwII region of Gαi (blue), highlighting the GIV F1685 and Gαi R208 interface. **B**. Sequence alignment of human NOD1, NOD2, Gαi (SwII), and rabbit Gαi (PDB:6MHF). Arrows indicate residues selected for mutagenesis; asterisks denote NOD2 chimeras incorporating corresponding NOD1 (*) or Gαi (**) sequences. **C-D.** GST pulldown assays of GST or GST–GIV with lysates of HEK293 cells expressing WT or mutant NOD2 (panel C). Bound HA-NOD2 was visualized by immunoblot with anti-HA mAb (IB). Quantification of three independent biological repeats is shown in D. **E-F.** GST pulldown assays of recombinant His–GIV-CT or its F1685A mutant (deficient in binding to Gαi^64^ and predicted in A as deficient in binding NOD2) with GST–NOD2-LRR. Bound proteins were detected by immunoblot; equal loading confirmed by Ponceau S (GST) and immunoblotting (IB; anti-His mAb). Quantification from three independent repeats of the assay in panel F. **G-I.** GST pulldown assays testing competition between GIV and Gαi for NOD2 binding. Recombinant His–GIV-CT was incubated with GST–NOD2-LRR in the presence of His–Gαi WT or the GIV-binding-deficient W211A mutant^88^. Bound GIV-CT (G) and unbound/soluble His-tagged proteins (H) were detected by anti-His immunoblotting. GST proteins were visualized by Ponceau S. Quantification of GIV-CT binding is shown in I. **J.** BRET-based cAMP responses in HeLa cells expressing WT NOD2 or the GIV-binding-deficient NOD2-L998A mutant following MDP stimulation. cAMP responses were measured using the CAMYEL biosensor and normalized to baseline. See **Figures S6A–B** for CAMYEL-related schematics and experimental workflow, **Figure S6C** for AUC analysis, and **Figure S6D** for the corresponding cAMP time courses. **K.** Schematic of the NOD2●GIV●Gαi interface in cAMP control. *Top:* WT NOD2 engages GIV during early microbial sensing, followed by GIV•Gαi engagement during the late cAMP plunge. *Bottom:* NOD2-L998A fails to engage GIV, disrupting the toggle and sustaining cAMP signaling. *Statistics*: Data are presented as mean ± SEM (n = 3 independent biological replicates). Statistical significance was assessed using an unpaired Student’s *t*-test for panels D, one-way ANOVA followed by Tukey’s multiple-comparisons test for panels F and I, and two-way ANOVA followed by Tukey’s multiple-comparisons test for panel J. Exact p values are indicated in the corresponding panels. *p*-value ≤ 0.05 is considered as significant.

To experimentally validate this model, we generated a panel of structure-guided NOD2 mutants, including point mutations in the 10th LRR and domain swaps replacing this region with sequences from NOD1 (non-binder) or the SwII loop of Gαi (binder) (**Figure 5B**). These structure-rationalized NOD2 mutants help assess specificity of binding and enable rigorous dissection of interface specificity while leaving intact the domains required for nucleotide binding, hydrolysis, and MDP recognition. These mutants also serve as molecular “handles” to test whether NOD2 and Gαi engagement were indeed exclusive and functionally separable. We also created a GEM motif mutant in GIV (F1685A).

Pulldown assays confirmed the predicted binding patterns (**Figure 5C-D**). A GIV F1685A mutant, known to abolish Gαi binding^64^, also failed to bind NOD2-LRR (**Figure 5E-F**), underscoring the shared binding determinant. BRET-based cAMP assays in cells expressing NOD2 mutants (e.g., L998A) demonstrated loss of GIV-dependent cAMP suppression in response to muramyl dipeptide (MDP) (**Figure 5J, Figure S6**), functionally uncoupling MDP sensing by NOD2 from signaling. Competition assays using a Gαi mutant (W211A^64^), that is specifically unable to bind GIV demonstrated that NOD2 and Gαi directly compete for occupancy of the GEM motif (**Figure 5G**-**I**), consistent with a molecular toggle mechanism.

Together with the spatial–temporal dynamics observed earlier (**Figures 3–4**), these structural and biochemical findings define the GEM motif as a molecular toggle: a single overlapping binding interface that is alternately engaged by NOD2 or Gαi, supporting mutually exclusive partner selection. This arrangement provides the mechanistic basis for how macrophages choreograph rapid switching between microbial sensing (via NOD2) and cAMP suppression (via Gαi) (**Figure 5K**), temporally segregating these functions to control pathogen clearance.

### Pharmacogenomic dissection implicates the GIV•Gαi axis in PKA-dependent microbial clearance

To rigorously define the contribution of the GIV•Gαi–cAMP axis to bacterial clearance, we paired selective pharmacologic inhibitors with recombinant TAT-GIV constructs in gentamycin protection assays using primary TGPMs (**Figure 6A**). As observed in THP1 macrophages (**Figure 2**), TGPMs recapitulated the signaling defects: GIV loss triggered an early, exaggerated cAMP surge within 15 minutes of *AIEC* LF82 infection (**Figure 6B**), accompanied by enhanced and sustained phosphorylation of PKA substrates (**Figure 6C** and **Figure S7A-B**).

**Figure 6:**
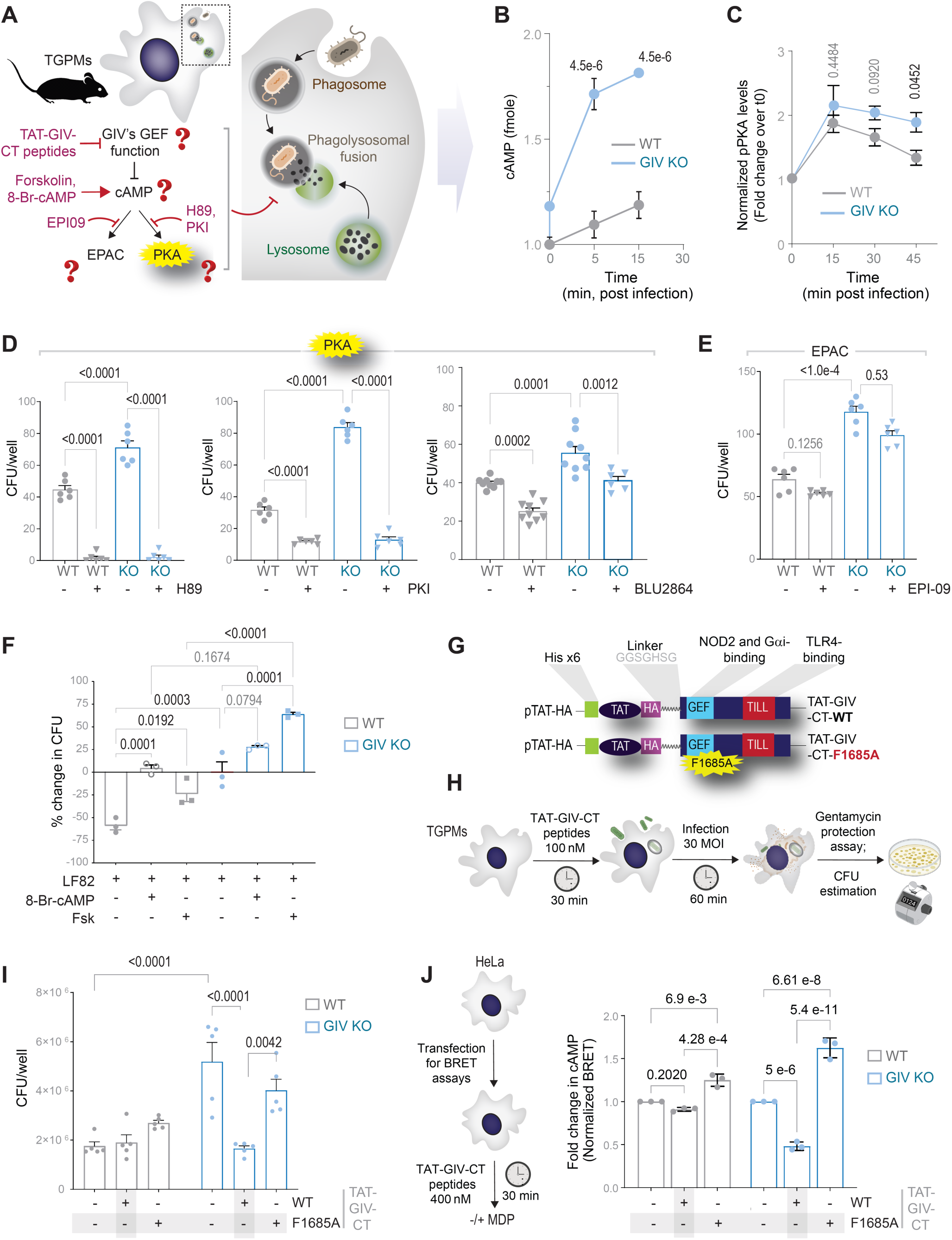
Pharmacogenomic dissection identifies PKA as the functional effector of GIV•Gαi-dependent cAMP control. **A.** Schematic of pharmacologic and genetic perturbagens used to dissect the cAMP cascade and GIV’s G protein regulatory function in phagolysosomal fusion and bacterial clearance. **B**. ELISA-based cAMP measurements in WT and GIV-KO TGPMs at 0, 5, and 15 min after *AIEC* LF82 infection. **C.** Time course of phospho-PKA substrate levels in WT and GIV-KO TGPMs following MDP pretreatment and *AIEC* LF82 infection, normalized to β-actin and expressed as fold change over baseline (0 min). See **Figure S7A-B** for representative immunoblots and quantification and **Figure S7F-G** for corresponding analyses in WT and GIV-KO HeLa cells. **D-E.** Viable bacterial counts in PMA-differentiated WT and GIV-KO THP1 macrophages, untreated (−) or pre-treated (+) with PKA inhibitors (H89, 30 µM; PKI 14–22, 10 µM; BLU2864, 30 µM) or EPAC inhibitor (ESI-09, 10 µM) for 30 minutes prior to *AIEC* LF82 infection (1 h, MOI 1:50). **F.** Percent change in viable bacterial counts in PMA-differentiated WT and GIV-KO THP1 macrophages, untreated (−) or pre-treated (+) with 8-Br-cAMP (10 mM) or Fsk (20 µM) for 30 minutes prior to *AIEC* LF82 infection (1 h, MOI 1:50). **G.** Domain architecture of TAT-GIV constructs^89^. The TAT transduction domain (TAT-PTD) fused to His- and HA-tags, linked to the C-terminal GIV segment (aa 1660–1870). Key binding sites are indicated for NOD2 and Gαi (blue) and TLR4 (red). WT TAT-GIV-CT (top) and the CT-F1685A mutant (bottom) are shown. See **Figure S7D-E** for TAT protein purification and cellular uptake. **H.** Workflow for the gentamicin protection assay in WT and GIV-KO TGPMs pretreated with WT or F1685A TAT-GIV CT mutant (100 nM, 30 min), followed by *AIEC* LF82 infection (MOI 1:50, 1 h). **I.** Viable bacterial counts in WT and GIV-KO TGPMs pretreated with WT or the F1685A TAT-GIV CT prior to *AIEC* LF82 infection. **J**. BRET-based cAMP assay using the CAMYEL biosensor in WT and GIV-KO HeLa cells. *Left*: experimental workflow: for treatment with 400 nM WT or F1685A TAT-GIV CT, followed by MDP stimulation (10 µg/ml). *Right*: normalized cAMP responses in WT and GIV-KO cells, ±WT or F1685A TAT-GIV CT. *Statistics*: Data are presented as mean ± SEM (n = 3 independent biological replicates). Statistical significance was tested using two-way ANOVA followed by Tukey’s multiple-comparisons test for panels B and I–J, an unpaired Student’s *t*-test for panel C, and one-way ANOVA followed by Tukey’s multiple-comparisons test for panels D–F. Exact p values are indicated in the corresponding panels; p ≤ 0.05 is considered significant.

To test causality, we inhibited PKA with three mechanistically distinct inhibitors: H89 (ATP competitive), PKI 14-22 (peptide-based, potent and selective) and BLU2864 (potent, PRKACA-specific). All three robustly rescued microbial clearance defect in GIV-KO macrophages (**Figure 6D**). Two orthogonal, high-specificity inhibitors (PKI, BLU2864) yielded identical results, making a shared off-target explanation unlikely. By contrast, EPAC inhibition with ESI-09 failed to restore clearance (**Figure 6E**), pinpointing PKA as the critical downstream effector of the GIV●Gαi–cAMP axis.

Manipulating cAMP levels directly supported this conclusion. Forskolin or 8-Br-cAMP impaired bacterial clearance in WT macrophages and further exacerbated clearance defects in GIV-KO cells (**Figure 6F**), supporting the conclusion that sustained cAMP signaling is incompatible with efficient phagolysosomal maturation and microbial clearance. Reintroduction of the GIV C-terminus via cell-permeable TAT-GIV rescued both microbial clearance (**Figure 6G-I**) and cAMP regulation (**Figure 6J**), whereas a mutant TAT-GIV lacking the G protein regulatory GEM motif failed to do so. These results establish that an intact GEM motif is required for the rescue activity of GIV-CT and that GIV-CT is sufficient to restore cAMP–PKA control and pathogen clearance in this setting.

Collectively, these pharmacogenomic perturbations define the GIV●Gαi–cAMP–PKA axis as a central, targetable checkpoint that governs phagolysosomal maturation and microbial clearance. By functionally linking the upstream toggle mechanism to its downstream effector arm, these studies reveal how precise disruption or restoration of individual nodes within this pathway can modulate macrophage antimicrobial capacity with high specificity.

### GIV•Gαi–dependent cAMP control preserves intestinal barrier integrity during infection

The intestinal barrier is a critical frontline in host defense, and its collapse is a hallmark of refractory IBD. Because refractory lamina propria macrophages exhibit defective microbial clearance and sustained cAMP–CREB signaling (**Figure 1B**), we next asked whether disrupting the GIV•Gαi axis compromises epithelial barrier integrity.

We first confirmed that clearance of multiple pathogens—*AIEC* LF82, *Salmonella* SL1344, *AIEC* NRG857c, and RS218—which was impaired in GIV-depleted THP1 macrophages, are restored by PKA inhibition (with PKI 14-22 validated earlier; **Figure 7B-C**). To phenocopy genetic loss of GIV, we used IGGi-11me, a selective inhibitor of the GIV•Gαi interaction^65^ (**Figure 7A**). Blocking this upstream interaction recapitulated the clearance defect, while downstream PKA inhibition restored it (**Figure 7D**), directly linking the molecular toggle to its effector cascade.

**Figure 7:**
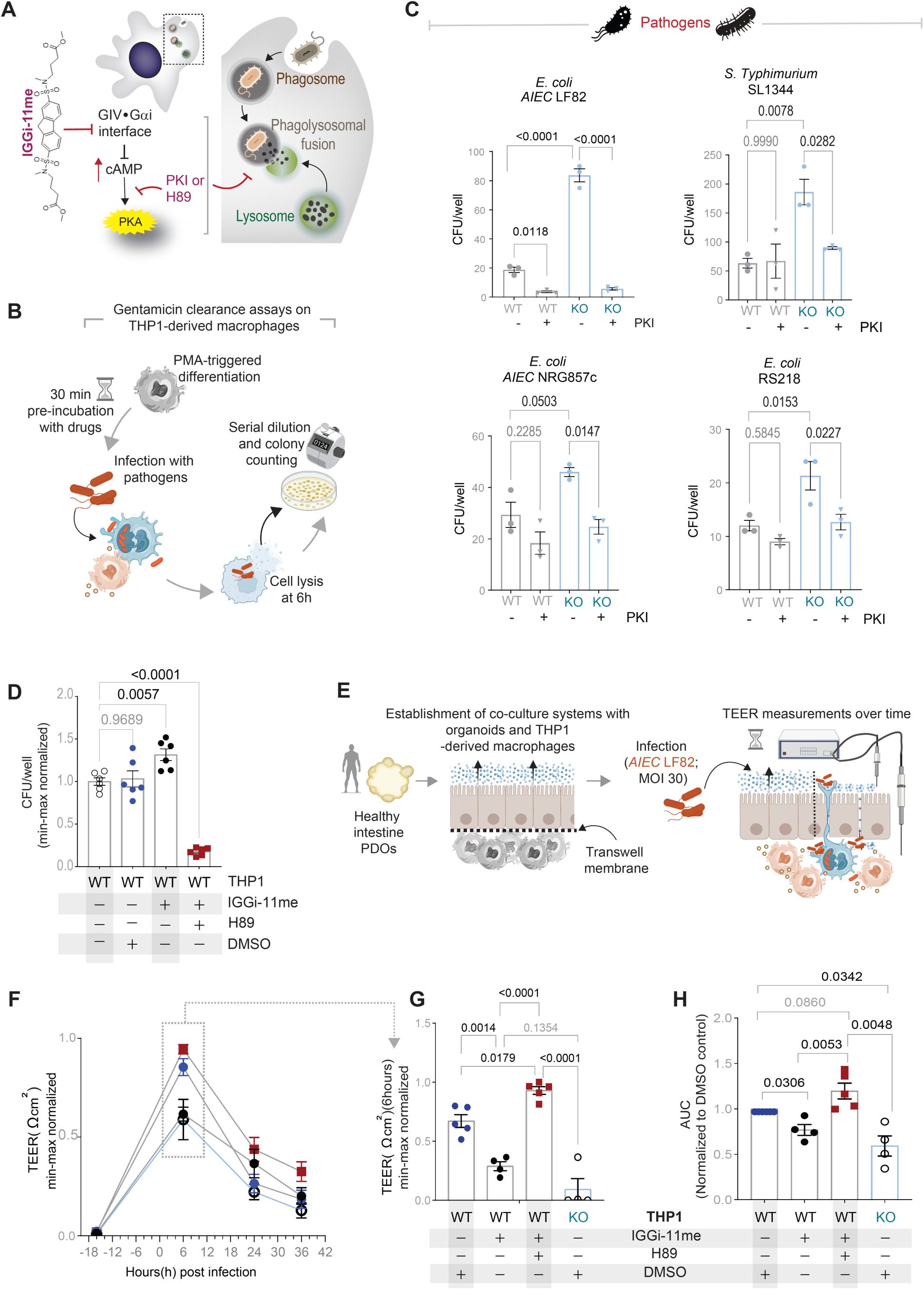
Perturbing GIV•Gαi–dependent cAMP signaling in macrophages highlights its control of microbial clearance and gut barrier integrity. **A.** Schematic of how pharmacologic perturbagens (used in C-H) modulate GIV’s Gαi-regulatory function and downstream cAMP effectors in phagolysosomal fusion. **B.** Workflow of the gentamicin protection assay in PMA-differentiated WT THP1 macrophages pretreated with GIV•Gαi inhibitor^65^ (IGGi-11me; 10 µM), PKA inhibitor H89 (30 µM), or DMSO, prior to *AIEC* LF82 infection (MOI 1:50, 1 hour). **C.** Viable bacterial counts in WT and GIV-KO THP1 macrophages ± PKA inhibition (PKI 14-22, 10 µM) during infection with multiple pathogens (*AIEC* LF82, *S. Typhimurium* SL1344, *AIEC* NRG857c, *E. coli* RS218; MOI 1:50, 1 hour). **D.** Viable bacterial counts in WT THP1 macrophages ± pharmacologic pretreatment (IGGi-11me, H89, or DMSO) prior to *AIEC* LF82 infection (MOI 1:30, 1 hour). **E.** Schematic of a trans well co-culture system with enteroid-derived monolayers (EDMs, apical side), WT or GIV-KO macrophages (basolateral side), and live microbes (apical side), used to model epithelium–microbe–macrophage crosstalk and barrier disruption during *AIEC* LF82 infection. **F.** Time course TEER (Ω·cm²) measurements of EDM-macrophage co-cultures upon *AIEC LF82* infection, shown as min–max normalized values (for comparison across experiments). See **Table S6** for raw TEER values (Ω·cm²). **G.** TEER at 6 h post-infection in EDM-macrophage co-cultures ± pharmacologic treatments (IGGi-11me, H89, or DMSO). **H.** Area under the curve (AUC) of TEER across infection time course, normalized to DMSO. *Statistics*: Data are presented as mean ± SEM, where applicable. Panel F presents TEER time-course data. Statistical significance was assessed using one-way ANOVA followed by Tukey’s multiple-comparisons test for panels C (*n* = 3 independent biological replicates), D and G (*n* = 4–5 independent biological replicates), and unpaired t-test with Welch’s correction for panel H (*n* = 4–5 independent biological replicates). Exact *p* values are indicated in the corresponding panels; *p* ≤ 0.05 was considered statistically significant.

To assess physiological consequences of this disruption, we turned to human enteroid-derived monolayers co-cultured with WT or GIV-KO macrophages and live *AIEC* LF82 (**Figure 7E**). In this transwell system—where 1.0 µm pores enable macrophage projections to directly sample microbes--transepithelial electrical resistance (TEER) measurements provided a quantitative readout of barrier integrity. Loss of GIV or pharmacologic disruption of the GIV●Gαi–cAMP axis led to premature epithelial barrier breakdown (**Figure 7F–H**), indicating a failure of macrophage-mediated barrier protection. Thus, disruption of the GIV•Gαi–cAMP–PKA checkpoint propagates from defective macrophage clearance to loss of epithelial barrier integrity, linking the molecular toggle to a tissue-level consequence relevant to refractory IBD.

## DISCUSSION

For fifty years, the host mechanism that terminates pathogen-induced cAMP surges has remained unresolved. Here, we identify GIV as that brake and show that it governs a broader signaling-state transition from microbial sensing to clearance. By alternately engaging NOD2 and Gαi, GIV converts pathogen sensing into a timed cAMP surge-to-plunge, terminates cAMP–PKA signaling, and enables phagolysosomal maturation. Time-resolved phosphoproteomics captures the same trajectory: WT macrophages progress from early signaling/cytoskeletal programs toward antimicrobial trafficking, whereas GIV-deficient cells remain trapped in a dysregulated PKA-associated state. Thus, GIV does more than restrain cAMP—it organizes the transition from microbial sensing to microbial destruction.

We highlight a few key advances emerging from this work:

### A host-encoded toggle switch reclaims control of cAMP

The NOD2●GIV●Gαi module functions as a biphasic molecular toggle switch that orchestrates the surge-and-plunge dynamics of cAMP in macrophages during microbial encounters. By alternately engaging NOD2 and Gαi, GIV permits the early surge but is required for efficient termination of cAMP–PKA signaling during the late plunge that drives phagolysosomal fusion and clearance. Live bacteria amplify the biphasic response, while heat-inactivated bacteria retain a partial response (**Figure S8A**). Acting from hemifusome–MVB microdomains, GIV provides a pathogen-proximal mechanism that complements global and compartmentalized regulation by PDEs (**Figure S8B**).

This mechanism also reveals a fundamentally different route to trimeric G-protein control in innate immunity (**Figure S8C**). Canonical extracellular cues, including chemokines, regulate cAMP through ligand-activated GPCRs at the plasma membrane. Here, an intracellular microbial sensor accesses the same G-protein/cAMP machinery without a GPCR: cytosolic GIV uses its GEM motif to couple NOD2-dependent pathogen sensing to Gαi activation. Thus, canonical GPCR and non-canonical GEM pathways converge on the cAMP–PKA axis from opposite sides of the membrane—one translating extracellular receptor activation, the other translating intracellular microbial sensing (**Figure S8C**). This expands the logic of G-protein signaling from receptor-encoded signaling at the cell surface to cytosolic, pathogen-proximal control of an antimicrobial state transition.

This discovery unites two previously separate domains: (i) microbial virulence mechanisms that elevate cAMP via toxins, effectors, or adenylyl cyclases, and (ii) host GEM (GIV–Gαi) modules that regulate cAMP in innate immunity. By bridging these, we identify GIV as a molecular nexus linking pathogen-induced cAMP surges to host recovery mechanisms that rebalance inflammation and clearance. In essence, immune resolution emerges from rapid, dynamic cAMP circuit rewiring at the host–microbe interface, orchestrated by GIV-centered hubs.

### GIV organizes the signaling-state transition from sensing to microbial clearance

The phosphoproteomic trajectory provides a systems-level view of what this molecular toggle accomplishes. GIV loss did not simply amplify infection-induced signaling; it rewired its temporal progression. In WT macrophages, early phosphorylation programs enriched in small-GTPase and cytoskeletal regulators gave way to endosomal trafficking and phagosome-maturation machinery. GIV-deficient macrophages instead entered an aberrant early state and progressed to persistent PKA–Rap and phagolysosomal signaling. Thus, the defining defect is not failure to initiate antimicrobial signaling, but failure to transition between signaling states. The subsequent functional experiments validate both predictions emerging from this trajectory: sustained cAMP–PKA activity and defective maturation of pathogen-containing compartments.

This temporal rewiring suggests that GIV operates above individual downstream pathways as an organizer of signaling-state transitions. By controlling when its GEM motif is occupied by NOD2 versus Gαi, GIV couples microbial recognition to the termination of cAMP–PKA signaling and progression into productive antimicrobial trafficking. In this view, the NOD2•GIV•Gαi toggle converts a biochemical timer into a cellular trajectory: sensing → cAMP–PKA restraint → trafficking → acidification → clearance. Loss of GIV uncouples these events, leaving macrophages signaling without successfully progressing to microbial destruction.

### A pathogen-proximal route to cAMP–PKA control

Our findings place GIV-dependent cAMP control in spatial proximity to hemifusome–MVB microdomains during maturation of pathogen-containing compartments. Whereas PDEs regulate cAMP lifetime—including at phagophore membranes^17^—prior work has linked this axis largely to cAMP–EPAC-dependent chemokine control^66^, rather than the PKA-dependent microbial clearance observed here. The NOD2•GIV•Gαi toggle instead acts upstream, coupling intracellular microbial sensing directly to inhibition of cAMP production. Localization of GIV and NOD2 near hemifusome–MVB structures places the molecular machinery for cAMP control in proximity to maturing pathogen-containing compartments, although compartment-restricted cAMP or PKA activity was not directly measured. Nucleotide- and conformation-dependent partner exchange adds temporal precision, linking microbial sensing to the late cAMP plunge and phagolysosomal maturation.

Although live-cell cAMP imaging could add spatial resolution, conventional BRET/FRET sensors require exogenous overexpression (which can be challenging in macrophages) and may perturb the native stoichiometry, partner exchange, or localization central to this mechanism. We therefore prioritized orthogonal readouts that preserve endogenous pathway organization—including cAMP measurements, CREB/PKA activity, endogenous proximity assays, and phagolysosomal acidification and maturation. Together, these approaches provide convergent support for biphasic cAMP regulation without perturbing the spatial and stoichiometric relationships that enable it.

### Structural basis and systems implications of conformation-dependent molecular toggle

Structural and mutagenesis analyses provide the mechanistic foundation for how GIV operates as a true molecular toggle. GIV typically scaffolds diverse surface^67^ and intracellular sensors—including ER stress (Grp78^68^), secretory flux (ER-Golgi intermediate compartment [ERGIC]/Arf1 GTPases^69,70^), double-stranded (ds)DNA-breaks (BRCA1^71^)—to trimeric Gαi, enabling GPCR-independent signaling. NOD2 uses this machinery differently: NOD2 and Gαi share the same binding determinant within GIV’s GEM motif. Leu998 in NOD2’s 10th LRR is essential for GIV binding; L998A disrupts this interaction and uncouples pathogen sensing from Gαi activation. Thus, mutually exclusive occupancy of a single motif creates toggle logic between NOD2-bound (sensing) and Gαi-bound (suppression) states.

We further identify the triggers governing this switch. MDP and ATP promote NOD2•GIV assembly; low pH–dependent conformational changes stabilize it; and NOD2 oligomerization accompanies dissociation of NOD2•GIV complexes and subsequent GIV•Gαi re-engagement. GIV preferentially recognizes ATP-bound NOD2 and GDP-bound Gαi, making the toggle conformation- and nucleotide-dependent. Consistent with established roles of terminal NLR LRRs in oligomerization and membrane localization^72–74^ , these findings position the shared GIV interface as a dynamic allosteric hub.

### System level implications of toggle switch control

Toggle switches are canonical motifs in systems biology^75,76^: they generate bistable, ultrasensitive, and rapid state transitions. Classical genetic toggles rely on transcriptional feedback and operate over minutes to hours^76,77^, whereas engineered phosphorylation-based protein toggles can respond within seconds^78^ ^75^.

Our findings reveal a similarly rapid, nucleotide-driven protein toggle embedded in innate immunity. The NOD2•GIV•Gαi module uses conformationally controlled partner selection to switch macrophages between cAMP surge and plunge states within minutes of microbial encounter. Phosphorylation of GIV by CDK5^79^ (enhancing) and PKCθ^80^ (disrupting) may further tune this interface; whether downstream PKA feeds back on the toggle remains an important question.

Thus, the NOD2•GIV•Gαi toggle provides a rapid decision-making mechanism that shifts macrophages from inflammatory restraint toward microbial clearance without transcriptional delay. Like engineered protein switches, it uses a single molecular interface to execute state transitions on the timescale of host–pathogen encounters.

### Translational implications in infection and IBD

This host-encoded toggle has broad translational implications. The NOD2-1007fs variant—a well-established CD risk allele and UC severity determinant^81,82^ truncates the 10th LRR, abolishing GIV binding and microbial clearance^40^. The mutation truncates NOD2, removes the 10^th^ LRR module, abolishes GIV binding, and is therefore expected to phenocopy NOD2-L998A mutant by uncoupling NOD2 from GIV•Gαi mediated cAMP suppression

Sustained cAMP signaling is a common microbial strategy, spanning *Mycobacterium*, *Salmonella, Vibrio* and pathoadapted *E. coli*. Our finding that PKA inhibition restores clearance despite GIV loss identifies PKA as a critical downstream effector and reveals two potentially druggable checkpoints: the NOD2•GIV•Gαi toggle and cAMP–PKA.

Therapeutically, restoring rather than disrupting this axis may be preferable. Emerging degradative molecular glues^83^ --monovalent small molecules that transiently induce protein–protein interactions--could, in principle, stabilize productive NOD2•GIV engagement and subsequent GIV•Gαi switching to restore physiologic cAMP control. In IBD, where NOD2 dysfunction can occur even without coding variants, targeting the GIV•Gαi–cAMP–PKA axis may restore microbial clearance and barrier protection without broad immunosuppression.

### Limitations of the study

This study has notable limitations. First, while structural modeling and mutagenesis identified the NOD2•GIV interface, high-resolution structural validation (e.g., cryo-EM) will be required to fully define the toggle mechanism. Second, phosphoregulation of the NOD2●GIV and GIV●Gαi interfaces was not systematically explored; future work should determine whether PKA or other kinases modulate toggle switching. Third, while our macrophage–enteroid co-cultures recapitulate key aspects of gut barrier physiology, allele-specific *in vivo* validation of the NOD2•GIV interface remains to be established, and *in vivo* infection and colitis models with GIV●Gαi inhibitors were not performed because IGGi-11me lacks bioavailability^65^. Nonetheless, prior evidence that PKA inhibition ameliorates colitis^84,85^ aligns with our conclusions and supports physiological relevance. Finally, the phosphoproteomic analysis was exploratory and was used primarily for hypothesis generation and pathway prioritization rather than definitive differential-phosphorylation inference. An important unresolved question is whether spatial PKA regulation is simply a downstream consequence of local cAMP control or constitutes an additional regulatory layer within pathogen-proximal signaling domains. Although the present study establishes cAMP–PKA as the functional effector axis downstream of GIV•Gαi, how local PKA substrates and feedback events shape phagosome maturation and toggle dynamics remains an important frontier.

In summary, our findings identify cAMP–PKA as a decisive checkpoint between microbial sensing and clearance. GIV, a cytosolic non-canonical GEM, couples intracellular NOD2 sensing to Gαi, enabling macrophages to terminate cAMP–PKA signaling and transition into antimicrobial trafficking. Thus, the NOD2•GIV•Gαi toggle determines when and where cAMP–PKA signaling must end for microbial destruction to begin.

## Resource Availability Lead Contact

Further information and requests for resources and reagents should be directed to and will be fulfilled by the lead contact, Pradipta Ghosh.

### Materials availability

Plasmids and other unique reagents generated in this study are available from the lead contact upon reasonable request, subject to applicable material transfer agreements and institutional policies of the University of California, San Diego.

### Data and Code Availability

- **Data:** The phosphoproteomic datasets generated in this study have been deposited with the ProteomeXchange Consortium via the PRIDE partner repository and are publicly available as of the date of publication. The datasets can be accessed at https://www.ebi.ac.uk/pride/archive/projects/PXD087532 The accession number is listed in the Key Resource Table. All data underlying the figures and tables are provided in the article and its supplemental information.
- **Code:** The code used for phosphoproteomic analyses is publicly available through the GitHub repository at <u>GitHub - sinha7290/NOD2-Encoded-Toggle-Switch</u>. The repository URL is provided in the Key Resources Table.
- **Additional information:** Additional information required to reanalyse the data reported in this paper is available from the Lead Contact upon request.

## Acknowledgments

This work was supported by the National Institutes of Health (NIH) grant (R01-AI141630, UG3TR003355, UG3TR002968, and R01-AI155696), a pilot award from the Propel a Cure Foundation and the Leona M. and Harry B. Helmsley Charitable Trust (to P.G.) and by NIH grants R21 AI149369, R21 AI156662, R01 AI161880, and R01 GM136202 (to I.K.). M.S.A. was supported by an American Heart Association Predoctoral Fellowship (25PRE1357971:https://doi.org/10.58275/AHA.25PRE1357971.pc.gr.227116). G.D.K. received support from the American Association of Immunologists (AAI) Intersect Fellowship Program for Computational Scientists and Immunologists and the American Heart Association Career Development Award (24CDA1268506: https://doi.org/10.58275/AHA.24CDA1268506.pc.gr.193658). S.S. was supported by an AAI Intersect Fellowship Program and M.M. was supported by a UC San Diego Agilent Center of Excellence Postdoctoral Fellowship. The authors would like to thank the University of California, San Diego - Cellular and Molecular Medicine Electron Microscopy Core (UCSD-CMM-EM Core, RRID:SCR_022039) for TEM sample preparation and microscope access. Data in this manuscript were generated at the UC San Diego Biomolecular and Proteomics Mass Spectrometry Facility, funded by NIH SIG grant S10 OD016234 (Synapt-HDX-MS) and S10 OD021724 (LUMOS Orbi-Trap). The authors acknowledge instrumentation resources at the UC San Diego Agilent Center of Excellence in Cellular Intelligence. The content is solely the authors’ responsibility and does not necessarily represent the official views of the Helmsley Charitable Trust or the NIH.

## Author contributions

M.A and P. G. conceptualized the project. M.A., D.C., K.C.P, G.D.K., M.N, S.R, J.S, K.Z, M.E, M.M, S.T.H and S.W were involved in data curation and formal analysis. S.S, with supervision from P.G, carried out phosphoproteomic analysis. K.C-P, with supervision from C.T. conducted all patient-derived organoid studies. S.R made the plasmid design, cloning and site-directed mutagenesis. A.S with supervision from J.Y, synthesized chemical matter. I.K. performed molecular modeling and NOD2 mutant design. M.A. and P.G. prepared figures for data visualization. P.G. wrote original draft; C.R.E, S.R, S.S, G.K and C.T helped in reviewing the manuscript. P.G supervised and acquired funding to support the study. All co-authors approved the final version of the manuscript.

## Inclusion and diversity

We worked to ensure sex balance in the selection of non-human subjects. One or more of the authors of this paper belongs to a minority group underrepresented in science.

## Declaration on use of generative AI

Generative AI has not been used to write the manuscript or generate figures de novo. AI-related tools were used to improve the readability and language of the text after it has been drafted.

## Key resources table

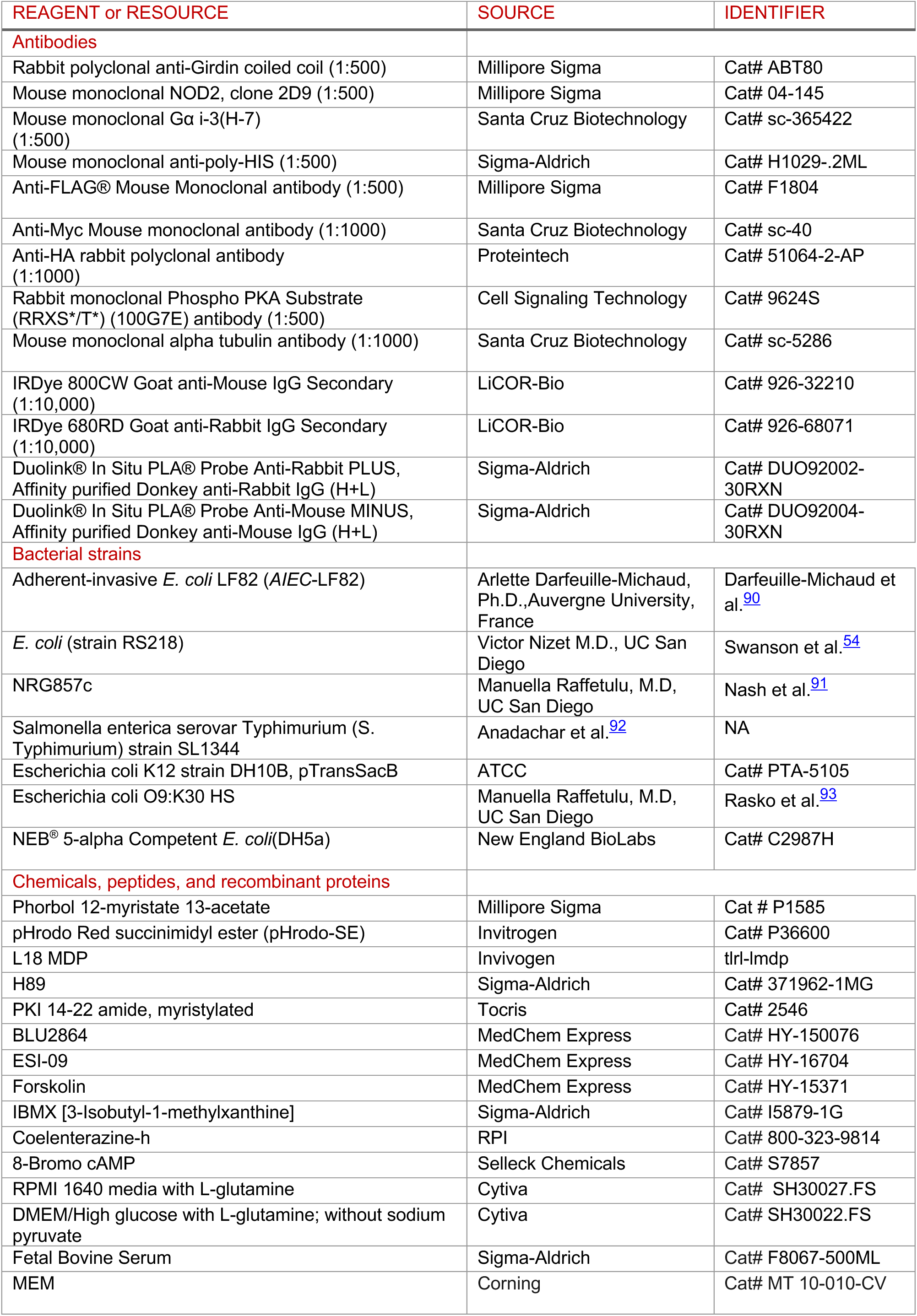
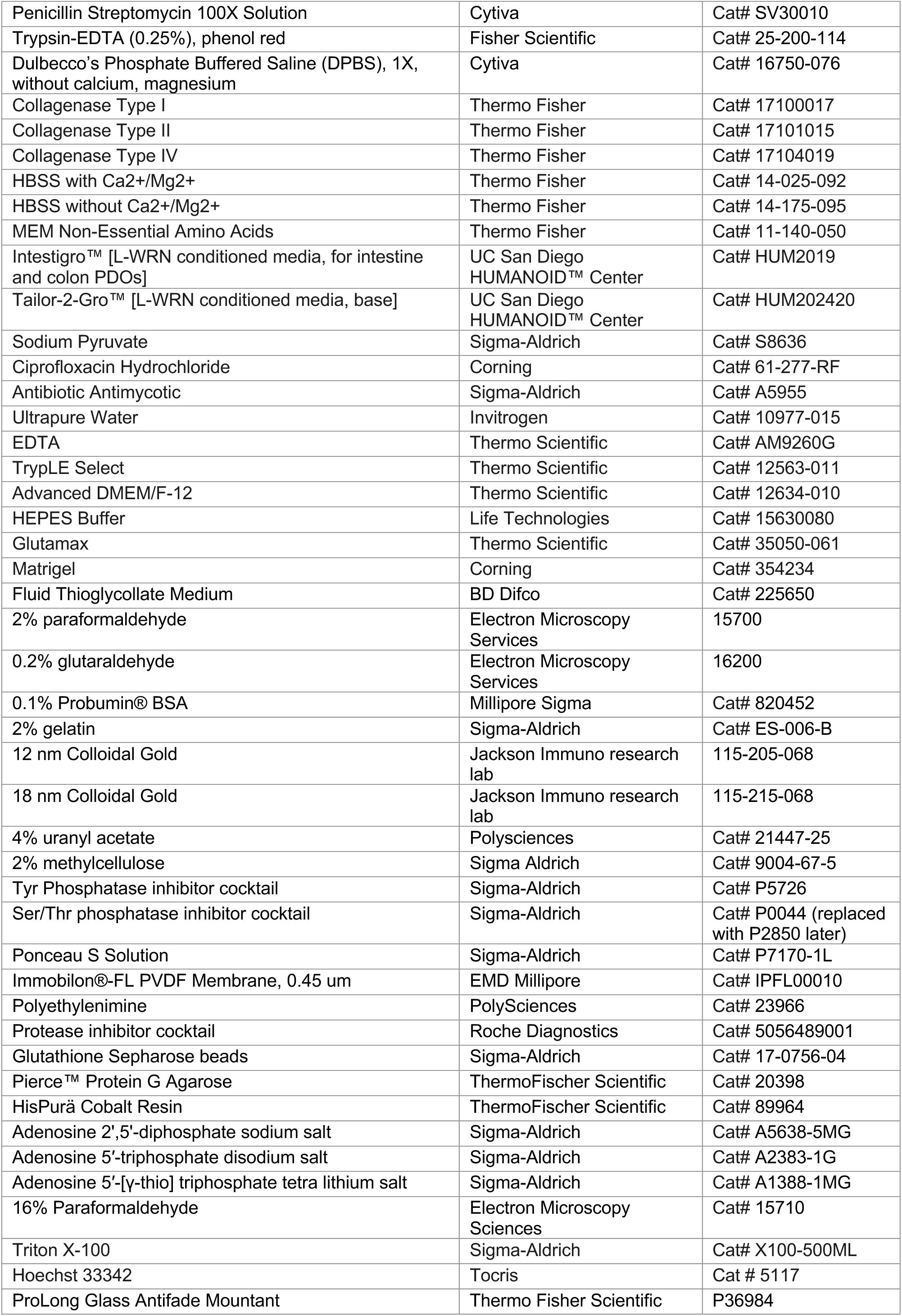
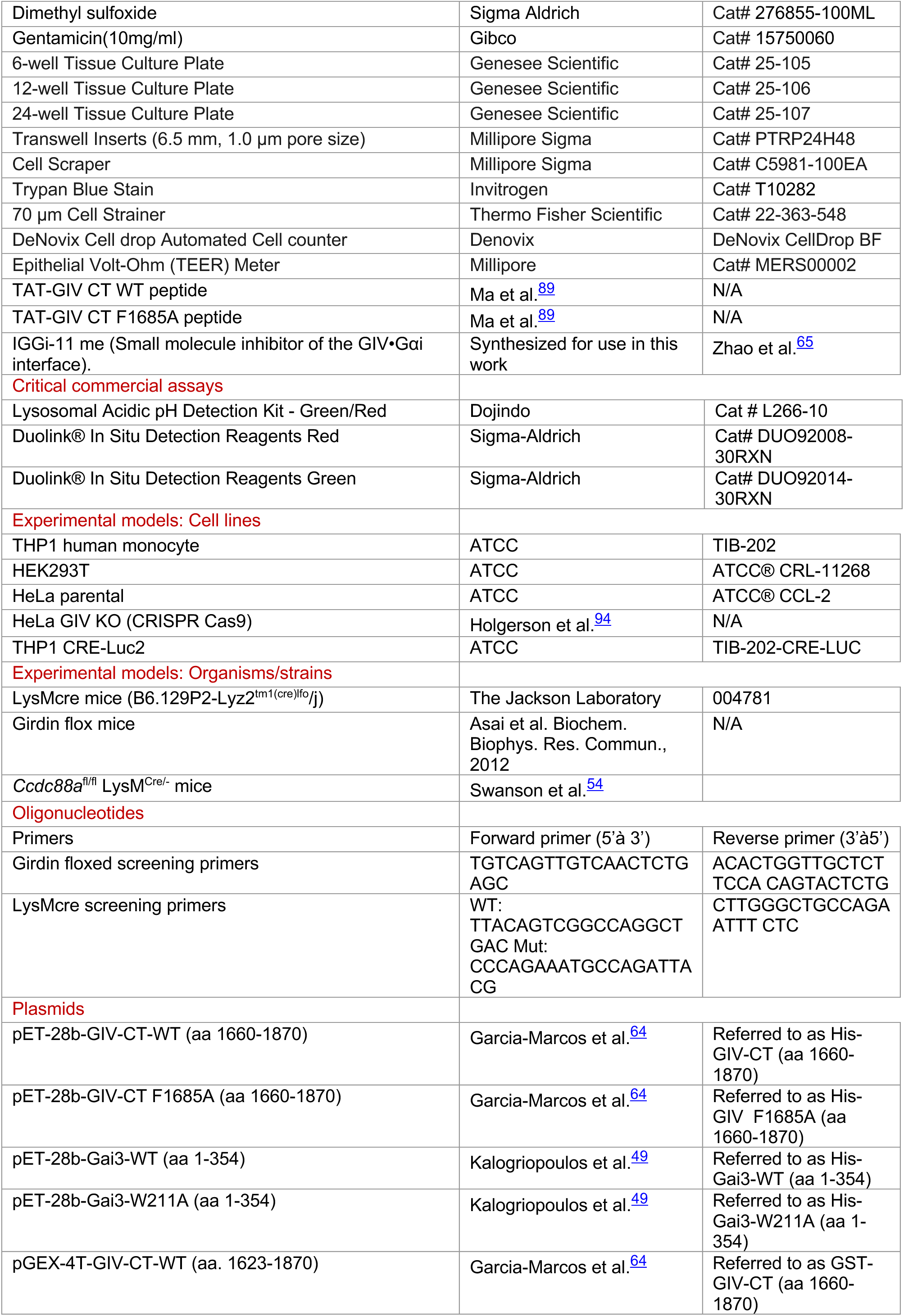
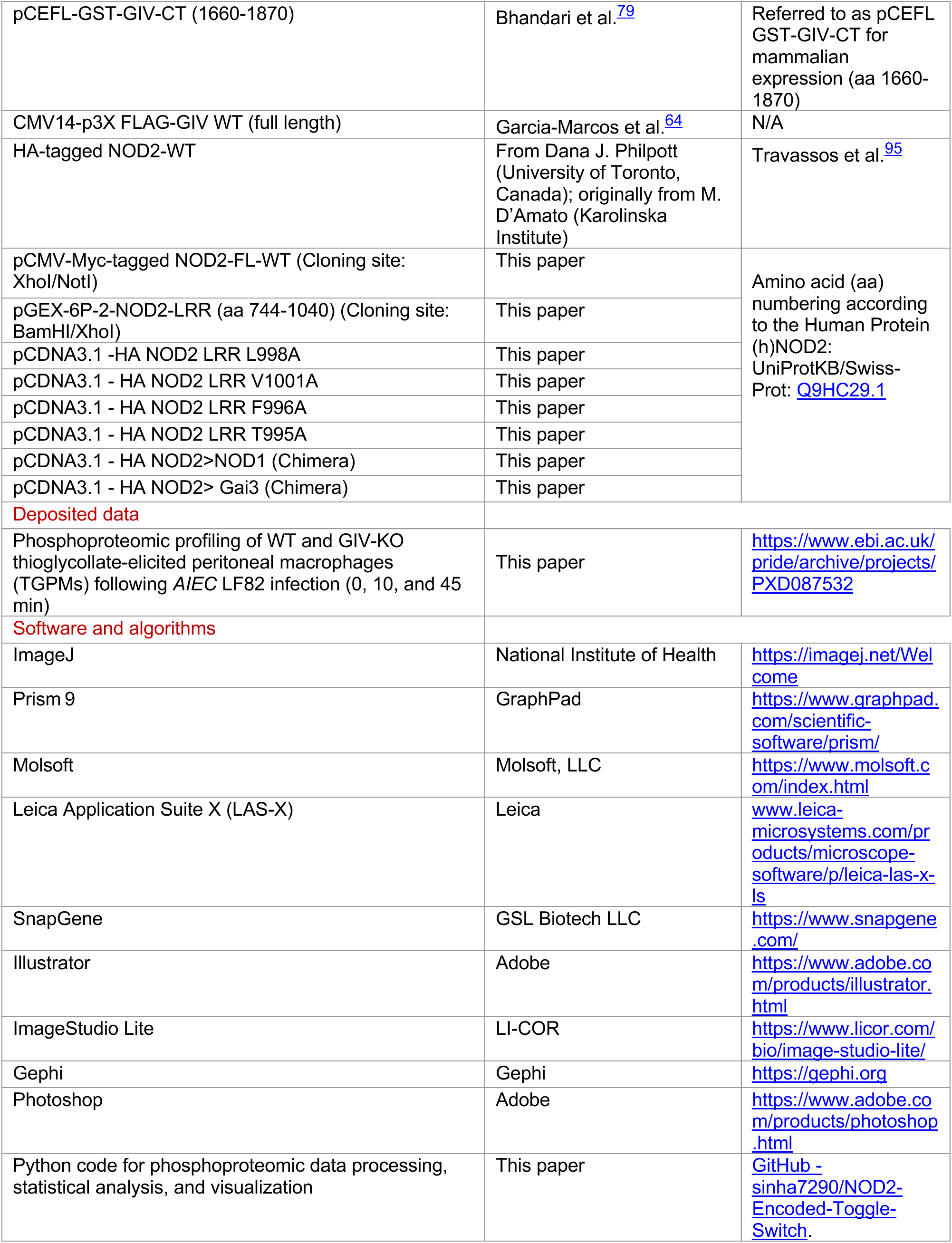

## STAR METHODS

### 1. Experimental models and Study Participant Details

#### Cell

Cell lines used in this study were obtained from either American Type Culture Collection (ATCC) or InvivoGen. All lines were routinely tested for mycoplasma contamination using commercially available PCR-based kits. Only validated, mycoplasma-free cells were expanded and used for experiments.

##### THP1

This human monocytic cell line was obtained from the ATCC and maintained in Roswell Park Memorial Institute 1640 (RPMI-1640) medium (Gibco-BRL). The medium was supplemented with heat-inactivated 10% fetal calf serum (Hyclone), 1% L-glutamine, and 1% penicillin/streptomycin (Gibco-BRL). The cells were incubated at 37°C in a 5% CO₂ incubator. Cultures were subcultured by centrifugation and resuspension at 2–4 × 10⁵ viable cells/ml, and maintained below 1 × 10⁶ cells/ml.

##### THP1-CRE-Luc

This CREB reporter derivative of the THP1 cell line was obtained from InvivoGen and maintained in RPMI-1640 medium supplemented with heat-inactivated 10-20% fetal calf serum, 1% L-glutamine, and 1% penicillin/streptomycin, as per the manufacturer’s direction. The cells were incubated at 37°C in a 5% CO₂ incubator. Cultures were subcultured by centrifugation and resuspension at 2–4 × 10⁵ viable cells/ml, and maintained below 1 × 10⁶ cells/ml.

##### HEK293T

This human embryonic kidney cell line was obtained from the ATCC and maintained in Dulbecco’s Modified Eagle Medium (DMEM; Gibco-BRL) supplemented with 10% fetal calf serum, 1% L-glutamine, and 1% penicillin/streptomycin. The cells were incubated at 37°C in a humidified atmosphere with 5% CO₂ and were routinely passaged at a dilution of 1:5 to 1:10 using 0.05% trypsin-EDTA when they reached approximately 80–90% confluency.

##### HeLa

This human cervical carcinoma-derived epithelial cell line was obtained from the ATCC and maintained in Dulbecco’s Modified Eagle Medium (DMEM; Gibco-BRL) supplemented with 10% fetal calf serum, 1% L-glutamine, and 1% penicillin/streptomycin. The cells were incubated at 37°C in a humidified atmosphere with 5% CO₂ and were routinely passaged at a dilution of 1:5 to 1:10 using 0.05% trypsin-EDTA when they reached approximately 80–90% confluency.

##### Thioglycolate-elicited murine peritoneal macrophages

Murine peritoneal macrophages were collected from peritoneal lavage of 8- to 12-wk-old C57BL/6 mice with 10ml of ice-cold RPMI (Roswell Park Memorial Institute medium) per mouse, 4 days after intraperitoneal injection of 3 ml of aged, sterile 3% thioglycolate broth (Becton Dickinson Difco™ Detroit, MI, USA) and cultured as described previously^96^. Peritoneal macrophages were passed through 70 µm filter to remove possible tissue debris contamination during harvesting. They were counted, centrifuged, and resuspended in RPMI-1640 containing 10% FBS and 1% penicillin/streptomycin and plated with the required cell density. The media was changed after 4 h to remove non-adherent cells. Cells were allowed to equilibrate overnight before various stimulations and/or treatments, as indicated in the figure legends.

#### Animal model

Girdin floxed mice were a generous gift from Dr. Masahide Takahashi (Nagoya University, Japan) and were generated as previously described^97^ LysM*^Cre^* mice (B6.129P2-Lyz2^tm1(cre)lfo/j^) were obtained from The Jackson Laboratory. Girdin*^fl/fl^*;LysM*^Cre+^*^/-^ mice were previously generated and characterized in our laboratory^54^ and maintained as homozygous for the Girdin floxed allele (fl/fl) and heterozygous for LysM*^Cre^*. Primers used for genotyping are listed in the *Key Resource Table (KRT)*. Mice were housed in the UC San Diego animal facility under a 12-hour light/dark cycle (30–70% humidity, room temperature maintained at 68–75 °F) with ad libitum access to standard chow and water. All animal studies were approved by the University of California, San Diego Institutional Animal Care and Use Committee (IACUC).

Male mice aged 8–12 weeks were used for the isolation of Thioglycolate-elicited peritoneal macrophages (TGPMs), with 6–9 mice used per experimental group. All animal experiments used male mice because this study did not assess sex as a biological variable.

#### Bacteria and bacterial culture

*Adherent Invasive Escherichia coli* strain LF82 (*AIEC-*LF82)^90^, *E. coli* K1 (RS218)^54^, *E. coli* O83:H1 (NRG857C)^91^, *Salmonella enterica* serovar Typhimurium (*SL1344*)^92^, *E. coli* K12 (commensal), *E. coli* O9 HS^93^ (commensal) and *E. coli* non-human adapted (DH5α) were cultured from a single colonies in LB broth for ∼6-8 hours on shaking, followed by overnight incubation under oxygen-limiting conditions without shaking to preserve pathogenicity as done previously ^98–100^. Bacterial density was estimated by measuring absorbance at 600 nm (OD_600_), followed by PBS washes and infection at the multiplicity of infection (MOI) indicated in figure legends.

### 2. Method details

#### Transcriptomic analysis

Publicly available transcriptomic data from the Gene Expression Omnibus (GEO) database (accession **GSE183620**) were analyzed using the *GEOquery* R package (Bioconductor). Supplementary *.csv.gz* files were retrieved and processed using *GEOquery* and the *readr* package, which allows for decompression and conversion to R data frames

Samples were filtered to include only ileal samples from individuals with Crohn’s disease and the genes *NOD2, CCDC88A, GNAI2, GNAI3, and GNAIS* using platform-level data about the samples. Samples were then grouped based on inflammation status. Log2 transformation was applied to the expression data, with a pseudocount of 1 added to avoid undefined values. Data visualization was performed using the *ggplot2* package to generate a grid of violin plots. Statistical significance was assessed using Welch’s paired t-test. P-values less than or equal to 0.05 were considered statistically significant.

#### Sample preparation for phosphoproteomics studies

WT and GIV-KO TGPMs were infected with *AIEC* LF82 (MOI 1:10) for 0, 10, and 45 minutes. After each time point of infection, cells were scraped, collected as pellets, snap-frozen in liquid nitrogen, stored at −80 °C, and submitted to the core facility for phosphoproteomic sample processing.

#### Phosphoproteomics Workflow

##### Sample Preparation and Mass Spectrometry

WT and GIV-KO TGPMs were infected with *AIEC* LF82 (MOI 1:10) for 0, 10, and 45 min. At each time point, cells were scraped, pelleted, snap-frozen in liquid nitrogen, stored at −80 °C, and processed at the core facility. Samples were lyophilized, reconstituted in 6 M guanidine-HCl (200 µL), and subjected to three boil–cool cycles. Proteins were precipitated with 90% methanol, pelleted, and resuspended in 8 M urea (100 mM Tris, pH 8.0). Reduction and alkylation were performed with 10 mM TCEP and 40 mM chloroacetamide, followed by dilution to 2 M urea. Proteins were digested with trypsin (1:50, 37 °C, 12 h), acidified (0.5% TFA), desalted using C18-StageTips, and quantified (BCA). Phosphopeptides (200 µg input) were enriched using High-Select Fe-NTA Phosphopeptide Enrichment (Thermo, A32992).

##### Mass Spectrometry Acquisition

Enriched phosphopeptides (0.6 µg/sample) were analyzed on a TimsTOF Pro 2 (Bruker) coupled to an EVOSEP ONE UPLC system. Peptides were separated on a 15 cm × 150 µm PepSep C18 column (1.5 µm beads, 58 °C) using an 88 min Evosep gradient (0.1% FA in water as solvent A; 0.1% FA/99.9% ACN as solvent B). Data-dependent acquisition (DDA) was performed using PASEF with a scan range of m/z 100–1700, 14 MS/MS scans per cycle, and charge states 2+ to 8+. Peptide identifications were made using PEAKS Server (Bioinformatics Solutions Inc.) with FDR <1% to construct a spectral library. For data-independent acquisition (DIA), identical chromatographic conditions were used, with 10 MS/MS scans per cycle and mobility-dependent collision energy ramping. DIA quantification was performed with PEAKS Server.

##### Phosphoproteomics data processing and visualization

Quantitative phosphopeptide Area values exported from PEAKS Server were analyzed in Python. UniProt accession identifiers were parsed from the protein-assignment field and mapped to mouse gene symbols using the UniProt ID-mapping system. For exploratory protein-level analysis, available phosphopeptide rows assigned to the same UniProt accession were averaged within each sample. This aggregation provides a protein-level phosphorylation-associated signal and does not quantify total protein abundance or phosphorylation at an individual phosphosite.

Biological replicates comprised independent TGPM samples, with n = 1 per genotype at 0 min and n = 2 per genotype at 10 and 45 min. For genotype comparisons at 10 and 45 min, replicate means were calculated separately for WT and GIV-KO samples. A pseudocount equal to one-half of the smallest positive Area value in the comparison dataset was added before log2 transformation. Log2 fold change was calculated as log2[(mean GIV-KO Area + pseudocount)/(mean WT Area + pseudocount)]. Welch’s two-sample t-test was applied to log2-transformed sample values, and P values were adjusted across all tested UniProt accessions using the Benjamini-Hochberg procedure. Proteins lacking sufficient replicate measurements for statistical testing were excluded from inferential comparisons rather than imputing missing measurements.

Because only one biological sample per genotype was available at 0 min, no inferential test or P value was calculated for the baseline comparison. Given the limited replication at 10 and 45 min (n = 2/genotype), nominal P-value-based genotype comparisons were considered exploratory and hypothesis-generating; Benjamini–Hochberg-adjusted P values are reported to provide multiple-testing context.

For the curated heatmap in Figure 1F, proteins representing signaling modules implicated by the phosphoproteomic analysis and relevant to the mechanistic questions addressed in this study were displayed using genotype–time-point mean phosphorylation-associated Areas. Heatmap selection was used for pathway-level visualization rather than statistical discovery. Remaining missing protein-by-sample values were set to zero for heatmap visualization only and were not replaced by zero during differential-abundance testing. Each protein row was standardized across the six genotype–time-point means by subtracting that protein’s row mean and dividing by its row standard deviation. Thus, red and blue indicate values above and below, respectively, the mean for the same protein across the displayed conditions; colors do not indicate statistical significance or permit quantitative comparison between proteins.

#### Data visualization

Previously reported upstream regulators identified using Ingenuity Pathway Analysis (IPA)<u>^41^</u>were plotted using R’s ggplot2 package. On the x-axis was bias-corrected z-score and on the y-axis was their adjusted p-value, with points colored by group as outlined in the original dataset. Network visualizations were based on Host-Host and Host-Microbe interaction data in *.csv* format. Native JavaScript was used to parse the data into edge and vertex arrays. Using the *Cytoscape.js* library, these were rendered as a multi-layer, interactive network in the user’s browser, with draggable nodes that could be repositioned to highlight specific patterns, relationships, or narrative insights within the data.

#### Gentamicin protection assay

THP1 monocytes were seeded at a density of 2.5 × 10⁵ cells per well in 24-well plates and differentiated into macrophage-like cells by treatment with 25 nM phorbol 12-myristate 13-acetate (PMA; Sigma-Aldrich, St. Louis, MO) for 16-18 hours prior to infection. Unless otherwise indicated, the same gentamicin protection assay protocol was applied to both PMA-differentiated THP1 macrophages and thioglycolate-elicited peritoneal macrophages (TGPMs). Cells (THP1 or TGPMs) were infected with the indicated bacterial strains for 1 hour in antibiotic-free RPMI 1640 supplemented with 10% FBS at 37 °C, 5% CO₂. After infection, cells were washed once with PBS and incubated with high-dose gentamicin (200 µg/mL, 90 min) to eliminate extracellular bacteria. The medium was then replaced with fresh RPMI containing low-dose gentamicin (50 µg/mL), and cells were incubated for an additional 3.5 hours. At the end of the low-dose gentamicin incubation, cells were washed with PBS and lysed in 1% Triton X-100. Cell lysates were serially diluted in PBS and plated on LB agar plates. Colony-forming units (CFUs) were enumerated following overnight incubation at 37°C to quantify viable intracellular bacteria. To test effect of MDP, cells were pre-treated overnight with L-18 MDP (10 µg/ml).

To test the impact of NOD2 activation, cells were pre-treated overnight with muramyl dipeptide (L18-MDP, Invivogen) (10 µg/mL), followed by gentamycin protection assays as above.

For pharmacologic studies, cells were pre-treated for 30 minutes with one of the following: PKA inhibitors H89 (30 μM), PKI 14–22 (10 μM), or BLU2864 (30 μM); EPAC inhibitor ESI-09 (10 μM); GIV●Gαi inhibitor IGGi-11me (10 uM); cAMP analog 8-Br-cAMP (10 mM); or adenylyl cyclase activator forskolin (20 μM). All compounds were maintained during infection and assay duration.

For cell penetrable peptide studies, cells were treated for 30 minutes with 100 nM cell-permeable TAT-GIV-CT peptides: wild-type (WT) to mimic GEF activity, or F1685A mutant to block it. All peptides were stored in small aliquots at -80 °C prior to use during the assay. These peptides have been expressed and purified exactly as described previously^54,89^, and mentioned briefly below (see later).

#### Measurement of intracellular cAMP via LANCE^®^ Ultra assay

Intracellular cyclic AMP (cAMP) levels were quantified using the LANCE® Ultra cAMP Assay Kit (PerkinElmer, Waltham, MA), according to the manufacturer’s instructions. Primary peritoneal macrophages (5,000 cells/well) were seeded into a 96-well low-volume white assay plates and infected with live or heat killed (dead) *AIEC* LF82 or NRG857c or RS218 or *SL134*4 with MOI 1:50 for 5 or 15 minutes. Heat inactivation was performed by incubating bacterial cultures at 65 °C for 30 minutes, followed by confirmation of sterility via plating on LB agar. After infection, Eu-cAMP tracer and ULight™ anti-cAMP antibody were added to each well, and plates were incubated at room temperature for 1 hour in the dark. Fluorescence resonance energy transfer (FRET) was measured using a Spark® 20M multimode microplate reader (Tecan, Männedorf, Switzerland). Cyclic AMP concentrations were calculated based on a standard curve generated in parallel using known concentrations of cAMP provided in the kit.

#### Plasmid constructs

HA-tagged full-length human NOD2 construct^95^ was obtained from Dr. Dana Philpott (University of Toronto, Canada), and the His-Myc–tagged full-length human NOD2 (His-Myc-NOD2^101^) was a generous gift from Dr. Santanu Bose (The University of Texas Health Science Center at San Antonio, San Antonio, Texas, USA). HA-NOD2 was cloned into pCDNA3.1 vector and served as the template for generating all mutant constructs. Site-directed mutants were created using the QuickChange II Site-Directed Mutagenesis Kit (Stratagene) following the manufacturer’s instructions. The sequences of mutagenesis primers are available upon request.

#### Transfection, lysis, and co-immunoprecipitation

HEK293T and HeLa cells were cultured in DMEM media containing 10% FBS and antibiotics, following ATCC guidelines. Cells were transfected with DNA plasmids using polyethyleneimine (PEI) following manufacturers’ protocols^102^. N-terminal HA- or Myc-tagged NOD2 and C-terminal FLAG-tagged GIV full length human proteins were co-expressed in HEK293T cells. 48hours post-transfection, cells were stimulated with 10 µg/mL MDP (L18-MDP, Invivogen) for 1, 3 and 6 hours and lysed in buffer containing 20 mM HEPES, pH 7.2, 5 mM Mg-acetate, 125 mM K-acetate, 0.4% Triton X-100, 1 mM DTT, 0.5 mM sodium orthovanadate, and phosphatase (for both Tyr and Ser/Thr) and protease inhibitor cocktails.

For immunoprecipitation, equal aliquots of clarified cell lysates were incubated with 2 μg of appropriate antibody [either anti-HA mAb or anti-FLAG M2Ab (Sigma Monoclonal ANTI-FLAG® M2, Clone M2)] for 3 hours at 4°C, followed by addition of protein G Sepharose beads (GE Healthcare; 40 µl 50% v:v slurry) and incubation for 1 hour at 4°C. Beads were washed four times (1 mL volume each wash) in PBS-T buffer [4.3 mM Na2HPO4, 1.4 mM KH2PO4, pH 7.4, 137 mM NaCl, 2.7 mM KCl, 0.1% (v:v) Tween 20, 10 mM MgCl2, 5 mM EDTA, 2 mM DTT, 0.5 mM sodium orthovanadate] and immune complexes were eluted by boiling in Laemmli’s sample buffer. Elutes were resolved on SDS PAGE and analyzed by immunoblotting with anti-HA, anti-Myc and anti-FLAG antibodies.

For nucleotide modulation studies, cells were first permeabilized with 4 μg/mL of streptolysin-O (Sigma-Aldrich, Budapest, Hungary) for 30 minutes at 37°C, then incubated with nucleotides (ADP, ATP, or ATPγS) as indicated.

#### Single and sequential DuoLink® Proximity Ligation Assay (PLA)

PMA-differentiated THP1 wild-type (WT) cells were seeded on sterile glass coverslips at a density sufficient to reach ∼60–70% confluency and treated with 25 nM phorbol 12-myristate 13-acetate (PMA; Sigma-Aldrich, St. Louis, MO) for 16–18 hours to induce differentiation. The next day, cells were stimulated with 10 µg/mL MDP for 1, 2, 3 or 6 hours. Following stimulation, cells were washed twice with phosphate-buffered saline (PBS) and fixed with 4% paraformaldehyde (PFA, in PBS) for 20 minutes at room temperature (RT), quenched with 0.1 M glycine in PBS for 10 minutes, permeabilized with 0.2% Triton X-100 in PBS for 1 hour at RT and blocked with 1% bovine serum albumin (BSA) and 0.1% Triton X-100 in PBS for 1 hour, as described previously^103^. Protein–protein interactions were assessed using the Duolink® In Situ Red and Green PLA kits (Sigma-Aldrich) following the manufacturer’s instructions.

For single PLA assay, coverslips were incubated overnight at 4°C with primary antibodies against GIV (rabbit, Millipore; ABT80, 1:150 dilution) and NOD2 (mouse, Santa Cruz, sc-56168,1:50 dilution), followed by incubation with species-specific PLA probes (PLUS and MINUS) for 1 hour at 37°C in a humidified chamber. Ligation and rolling circle amplification were performed. Red PLA detection reagents (Ex/Em: 566–594 nm) visualized GIV●NOD2 interactions. For GIV●Gαi3 detection, rabbit anti-GIV (1:150) and mouse anti-Gαi3 (1:50) were used, followed by Green PLA detection reagents (Ex/Em: ∼488 nm). Each single PLA interaction was conducted on different cover slips—in parallel---from the same experiment but imaged, analyzed and quantified independently.

For the dual PLA assay, GIV●NOD2 interaction was visualized using the red PLA protocol. After PBS washes, coverslips were incubated overnight with mouse anti-Gαi3, and the green PLA protocol was repeated to detect GIV●Gαi3 interactions

In all assays, nuclei were counterstained with 1 µg/mL DAPI to visualize nuclei and coverslips were mounted using ProLong™ Gold. Imaging was performed using Leica Stellaris confocal microscope or BioTek Cytation 10. PLA signal quantification (puncta/cell) was done in ImageJ using background subtraction, thresholding, and the “Analyze Particles” tool.

#### Transmission electron microscopy (TEM) and immunogold EM staining

Cells are fixed with 2% paraformaldehyde + 0.2% Glutaraldehyde in 0.1M sodium cacodylate buffer pH 7.4 for 5 min at room temperature and for 1 hour on ice, then pelleted in 10% gelatin. The pellet in gelatin was then cut into 0.5 mm pieces and incubated overnight in 2.3 M sucrose in PBS. The pieces were mounted onto aluminium pins and frozen into liquid nitrogen. 70 nm cryosections were produced at -120⁰C using a Leica cryo-ultramicrotome and collected on formvar/carbon coated 300 mesh gold grids. For immunolabelling, the sections on grids were washed with PBS and blocked with 1% BSA in PBS for 2 min. Next, they were incubated with mouse anti-NOD2 antibody (Santa Cruz, sc-56168, 1:50 dilution in 1% BSA in PBS) and rabbit anti-GIV antibody (Millipore sigma, ABT80; 1:50 dilution in 1% BSA in PBS) followed by secondary antibodies 18 nm colloidal gold of donkey anti-rabbit IgG and 12 nm gold of donkey anti-mouse IgG (Jackson ImmunoResearch Laboratories, Inc.). After washing with 0.1% BSA in PBS and PBS, the sections on the grids were fixed with 1% glutaraldehyde, washed extensively with water, stained for 5 minutes with 2% uranyl acetate, and coated with 0.2% uranyl acetate in 1.8% methylcellulose.

Images were acquired with a transmission electron microscope JEOL 1400 plus, operated at 80 KeV and equipped with a bottom-mounted 4k x 4k camera Gatan One View.

#### Protein expression and purification

His-tagged and GST-tagged recombinant proteins were expressed in *E. coli* strain BL21 and purified as previously described^104^ ^105^ ^106^. Bacterial cultures were induced with 1 mM isopropyl β-D-1-thio-galactopyranoside (IPTG) either overnight at 25°C. Bacterial culture (1l) were pelleted after induction and resuspended in 20 mL GST-lysis buffer (25 mM Tris·HCl, pH 7.5, 20 mM NaCl, 1 mM Ethylenediaminetetraacetic acid (EDTA), 20% [v/v] glycerol, 1% [v/v] Triton X-100, 2 × protease inhibitor mixture [Complete EDTA-free; Roche Diagnostics]) or in 20 mL His-lysis buffer (50 mM NaH_2_PO_4_ [pH 7.4], 300 mM NaCl, 10 mM imidazole, 1% [vol/vol] Triton X-100, 2 × protease inhibitor mixture [Complete EDTA-free; Roche Diagnostics]) for GST or His-fused proteins, respectively. Bacterial lysates were sonicated (three cycles, with pulses lasting 30 s/cycle, and with 2 minutes interval between cycles to prevent heating), centrifuged at 12,000×*g* for 20 minutes at 4°C. Supernatant (solubilized proteins) were affinity purified on glutathione-Sepharose 4B beads (GE Healthcare) or HisPur Cobalt Resin (Pierce), dialyzed overnight against PBS, and stored at −80°C.

#### GST pulldown assays

Recombinant GST-tagged proteins including GST alone (control), GST-NOD2 LRR wild-type, and GST-NOD2 LRR mutants were expressed in *E. coli* strain BL21 (DE3) and purified as previously described^54^. Bacterial cultures were induced with 1 mM IPTG and incubated overnight at 25 °C. Cells were pelleted and resuspended in GST lysis buffer [25 mM Tris-HCl (pH 7.5), 20 mM NaCl, 1 mM EDTA, 20% (v/v) glycerol, 1% (v/v) Triton X-100, and 2× protease inhibitor cocktail], then briefly sonicated (30-second pulses on/off for 5 minutes). Lysates were clarified by centrifugation at 13,000 rpm for 45 minutes. Protein concentrations were determined using BSA standards, and aliquots were stored at –80 °C.

Equimolar concentrations of GST-tagged proteins were immobilized onto glutathione-Sepharose beads by incubation in binding buffer [50 mM Tris-HCl (pH 7.4), 100 mM NaCl, 0.4% (v/v) Nonidet P-40, 10 mM MgCl₂, 5 mM EDTA, 2 mM DTT, and protease inhibitors] for 1 hour at 4 °C or overnight with gentle tumbling. After washing, beads were incubated with equimolar amounts of purified His-tagged wild-type or mutant F1685A GIV-CT proteins or with His Gai WT or His Gai W211A mutant protein (resuspended in the same buffer) for 4 hours at 4 °C. Ten percent of the His-tagged input was retained as control.

Following incubation, beads were washed four times with phosphate wash buffer [4.3 mM Na₂HPO₄, 1.4 mM KH₂PO₄ (pH 7.4), 137 mM NaCl, 2.7 mM KCl, 0.1% (v/v) Tween-20, 10 mM MgCl₂, 5 mM EDTA, 2 mM DTT, 0.5 mM sodium orthovanadate], then eluted twice in Laemmli buffer [5% SDS, 156 mM Tris base, 25% glycerol, 0.025% bromophenol blue, 25% β-mercaptoethanol] by boiling at 95 °C for 5 minutes. Eluates were pooled, resolved by 10% SDS-PAGE, and transferred to PVDF membranes.

Membranes were stained with Ponceau S to confirm transfer efficiency and equimolar loading of GST-tagged proteins, then blocked in PBS containing 5% non-fat milk. Immunoblotting was performed using mouse anti-His antibody (Sigma; 1:1000 dilution) and a custom rabbit polyclonal anti-GIV-CT antibody (1:1000 dilution; epitope: C-terminal 18 amino acids of human GIV, validated previously^107^). Detection and quantification were carried out using a LI-COR Odyssey imaging system with dual-color infrared detection. Final figures were assembled using Adobe Photoshop and Illustrator (Adobe, San Jose, CA, USA).

#### Homology modeling and protein docking

To illustrate the putative structural basis of the GIV•NOD2 interaction, we generated a comparative model of the 10th leucine-rich repeat (LRR) of human NOD2 (aa sequence 986-KILKLSNNCITYLGAEALLQALERNDTILEVWLRGNTFSLEEVDKLGCRDTRLLL-1040) with the GEM motif of GIV (aa sequence 1674-GSPGSEVVTLQQFLEESN-1691) Modeling and structural analyses were performed in Internal Coordinate Mechanics (ICM-Pro) software^108,109^. We noticed that in the crystal structure of the homologous rabbit NOD2 (PDB 5IRN^86^), the corresponding LRR10 region shares structural and sequence homology with the Switch-II region of human Gαi3 complexed with GIV GEM motif (PDB 6MHF^49^). Capitalizing on this observation, a model of human NOD2 LRR10 was built *ab initio* and assigned to a conformation of its rabbit counterpart observed in the crystal structure. The model was superimposed with the Sw-II region of Gαi3 from PDB 6MHF. The GIV GEM molecule was then combined with the model of NOD2 LRR10, the backbone atoms of the two molecules tethered to the conformations observed in PDB 6MHF and PDB 5IRN, respectively, and harmonic distance restraints set to retain the backbone hydrogen bonds that mediate the anti-parallel β-sheet between the two molecules in PDB 6MHF. Global side-chain Monte Carlo conformational search (4×10^4^ steps) with concurrent local backbone minimization was performed on the system to find the optimal relative geometry of the two molecules, with the objective function including steric complementarity, electrostatics, hydrogen bonding, tethers, and distance restraint terms. Top-scoring conformation was selected for visualization and used to propose interface-disrupting mutations.

#### Sequence alignment

Amino acid sequences corresponding to the leucine-rich repeat (LRR) domain of human and rabbit NOD2 (wild-type and 1007fs mutant), human NOD1, and the Switch II (Sw-II) region of human Gαi3 were retrieved from the Ensembl Genome Browser (https://www.ensembl.org). Multiple sequence alignments were performed using MolSoft ICM. Residue annotations and structural references were guided by crystal structures available in the Protein Data Bank: rabbit NOD2 (PDB: 5IRN) and the GIV–Gαᵢ3 Switch II complex (PDB: 6MHF).

#### Quantitative immunoblotting

All immunoblots were imaged and quantified using a LI-COR Odyssey imaging system with dual-color infrared detection using its densitometry feature. Bound proteins were normalized to soluble proteins and % bound was expressed as fold change compared to internal controls on the same immunoblot to avoid confounding variables such as antibody potency, transfer efficiency, etc.

For the assessment of PKA activity in cells, WT and GIV KO TGPMs or WT and GIV KO HeLa cells^94^ were pre-stimulated with muramyl dipeptide (L18-MDP; InvivoGen, 10 µg/mL) for 30 minutes, followed by infection with *AIEC* strain LF82 at a multiplicity of infection (MOI) of 1:50 for 15, 30, or 45 minutes. Cells were lysed in HEPES lysis buffer [20 mM HEPES (pH 7.2), 5 mM magnesium acetate, 125 mM potassium acetate, 0.4% Triton X-100, 1 mM DTT] supplemented with 500 µM sodium orthovanadate, phosphatase inhibitor cocktail (Sigma-Aldrich), and protease inhibitor cocktail (Roche, California, USA). Lysates were passed through a 28G needle on ice and centrifuged at 10,000 × *g* for 10 minutes at 4°C. Phospho-PKA activity was assessed by immunoblotting with a rabbit monoclonal anti-phospho-PKA substrate (RRXS*/T*) antibody (Cell Signaling Technology; 1:1000 dilution), and signal was quantified as total lane intensity. α-Tubulin (mouse monoclonal; 1:1000 dilution) served as a loading control.

#### CREB reporter assays using THP1 -derived macrophages

To assess signal transduction downstream of the cAMP/PKA/CREB and innate immune pathways, two THP1 reporter lines were used: THP1-CRE-Luc2 (ATCC Cat# TIB-202-CRE-LUC2): This monocytic cell line stably expresses firefly luciferase under the control of a cAMP response element (CRE) promoter, enabling quantification of CREB-mediated transcription. Cells were differentiated with 25 nM PMA for 18–24 hours, rested, and then infected with live or heat inactivated bacteria at the indicated MOI. Luciferase activity was measured using the D-Luciferin substrate and normalized to untreated control.

These assays allowed time-resolved profiling of CREB dependent transcriptional responses to microbial signals and pharmacological perturbations.

#### Expression and purification of TAT-GIV-CT

Cloning of TAT-GIV-CT wild-type (WT) has been described elsewhere^89^. The GEF-deficient mutant TAT-GIV-CT-FA was generated using QuikChange II (Stratagene) following the manufactureŕs instructions. Constructs were expressed using BL21(DE3)-pLysS (Invitrogen) and Terrific Broth (BioPioneer) supplemented with additives as per auto-induction protocols outlined by William F. Studier^110^. Briefly, cultures of bacteria were grown at 300 rpm at 37°C for 5 hours, and subsequently at 25°C overnight. Cells were lysed in 10 mL of buffer [20 mM Tris, 10 mM Imidazole, 400 mM NaCl, 1% (vol:vol) Sarkosyl, 1% (vol:vol) Triton X-100, 2 mM DTT, 2 mM Na3oV4 and protease inhibitor mixture (Roche Diagnostics) (pH 7.4)], sonicated (3 × 30 s), and clarified by centrifugation (12,000 × g for 20 minutes at 4°C). His-tagged proteins were affinity-purified on Ni-NTA agarose resin (Qiagen) for 4 hours at 4 °C and eluted in buffer [20 mM Tris, 300 mM Imidazole, 400 mM NaCl, pH 7.4]. As a final step, size exclusion chromatography (SEC) was performed using a 60 cm column packed with Superdex 200 pg (34 µm) to remove residual microbial LPS and ensure endotoxin-free preparations suitable for cell-based assays. Proteins were stored at −80 °C in TBS containing 400 mM NaCl.

#### CAMYEL Gαi-directed cAMP response assay

These assays were carried out exactly as detailed elsewhere^111^. Briefly, on day 1, HeLa cells were plated at 600,000 cells per well in 6-well plates using DMEM supplemented with 10% FBS. On day 2, media was replaced with fresh DMEM + 10% FBS. Cells were transfected with a total of 3 µg DNA per well using TransIT-X2 reagent, comprising: 1 µg V2-CAMYEL, 1 µg WT Gαi and 1 µg NOD2 or NOD2 mutant construct (excluded in TAT peptide assay sets). On day 3, cells were trypsinized, resuspended in DMEM + 10% FBS at 700,000 cells/mL, and 100 µl aliquots (70,000 cells) were re-plated into 96-well white/clear bottom plates. Cells adhered for 5–6 hours at 37°C/5% CO₂. For MDP-treatment, media was replaced with 100 µl DMEM + 10% FBS containing 10 µg/mL MDP, and cells were incubated overnight. On day 4, media was replaced with 100 µl DMEM containing 0.2% FBS and 1 µg/mL LPS. For TAT peptide experiments, cells were preincubated with 400 nM TAT peptides for 30 minutes prior to MDP treatment. Before reading, media was carefully removed and replaced with 70 µl serum-free assay buffer (PBS + 0.1% D-Glucose + 0.05% BSA).

Luciferase substrate Coelenterazine-h (final 10 µM) and IBMX (final 100 µM) were added (10 µl each from stock solutions of 0.1 mM and 1 mM, respectively). Plates were incubated at room temperature for 5 minutes. Luminescence was measured using a Tecan SPARK 20M plate reader at 485 nm and 515 nm emission wavelengths for 3 minutes to establish a basal CAMYEL signal. Forskolin (FSK) was then added (final concentration 10 µM), and luminescence was measured for an additional 15 minutes.

##### Data Analysis

cAMP levels were calculated as the inverse BRET ratio (485 nm emission / 515 nm emission). Net BRET values were computed by subtracting the average inverse BRET during 3–6 minutes pre-FSK from the average during 10–12 minutes post-FSK. The assay was repeated in four independent biological replicates with three technical replicates each. Further data normalization was done by minimum maximum scoring. Data were graphed using GraphPad Prism 9, and main text figures show averages of three to four biological replicates.

#### Lysosomal pH and mass measurements using pHLys Red and LysoPrime Green

WT and GIV-KO THP1s were seeded in black, clear-bottom 96-well plates (Agilent) at 5 × 10^4 cells per well in 100 μL RPMI 1640 supplemented with 10% fetal bovine serum (FBS) and differentiated with 25 nM PMA for 24 h. Cells were then washed and cultured overnight in RPMI 1640 containing 0.2% FBS prior to infection.

Lysosomal acidification and relative lysosomal mass were measured using the pHLys Red/LysoPrime Green Lysosomal pH Indicator Kit (Dojindo, Kumamoto, Japan; Cat. No. L226-10), following established fluroscence-based approaches for assessing lysosomal acidity^56,57^. pHLys Red, a pH-sensitive lysosomal probe, and LysoPrime Green, a pH-insensitive lysosomal probe used to account for lysosomal mass, were diluted in prewarmed HBSS to final working dilutions of 1:1,000 and 1:2,000, respectively. Cells were washed twice with HBSS and incubated with LysoPrime Green for 30 min at 37°C protected from light. Deep Red-labeled, heat-inactivated *AIEC* LF82 bacteria were added at a multiplicity of infection (MOI) of 50 together with pHLys Red and Hoechst nuclear stain (1:2,000 dilution), and cells were incubated for an additional 30 min at 37°C. Cells were subsequently washed twice with HBSS and maintained in phenol red-free imaging medium.

Live-cell fluorescence imaging was performed immediately using a BioTek Cytation imaging system (Agilent). Images were acquired using excitation/emission settings of 456/540 nm for LysoPrime Green, 562/586 nm for pHLys Red, and 630/660 nm for Deep Red-labeled *AIEC* LF82. Identical acquisition settings were maintained across experimental conditions.

Images were analyzed using BioTek Gen5 software. Individual regions of interest (ROIs) were identified, and mean fluorescence intensities for pHLys Red and LysoPrime Green were quantified within corresponding ROIs. A lysosomal acidification index was calculated as the pHLys Red/LysoPrime Green fluorescence ratio, thereby normalizing the pH-sensitive signal to relative lysosomal mass. Ratios were normalized to the corresponding uninfected control. LysoPrime Green fluorescence was analyzed separately as a measure of relative lysosomal mass.

#### Phagosomal acidification assay using flow cytometry

Phagosomal acidification was assessed using an established pHrodo-based flow cytometric approach^59,60^. Briefly, heat-inactivated *AIEC* LF82 were labeled with pHrodo Red succinimidyl ester (pHrodo-SE; Thermo Fisher Scientific) according to the manufacturers’ recommendations. THP1 WT and GIV-KO cells were seeded at 1.5 × 10⁶ cells/well in 12-well plates in RPMI containing 10% FBS, PSG, and 25 nM PMA. Following overnight differentiation, the medium was replaced with phenol-red-free RPMI containing 0.2% FBS without PSG.

On the day of the assay, THP1 cells were challenged with pHrodo-labeled LF82 at an approximate MOI of 50. Cells were incubated for 30 min at 37°C, after which plates were immediately placed on ice to stop further phagocytosis. Infection medium was removed, and cells were collected by scraping in 500 μL ice-cold FACS buffer. Cells were resuspended to obtain a single-cell suspension and transferred to a 96-well plate for flow cytometric analysis.

Flow cytometry was performed using a NovoCyte Quanteon (Agilent Technologies, Santa Clara, CA), and data were analyzed using NovoExpress (Agilent Technologies) and FlowJo. Cells were identified based on FSC-A and SSC-A characteristics, and pHrodo fluorescence was subsequently assessed within the gated cell population. Unstained controls were used to establish the pHrodo-positive gate. Both the percentage of pHrodo-positive cells and pHrodo median fluorescence intensity (MFI) were determined. Because pHrodo fluorescence increases in acidic environments, these measurements were interpreted as readouts of bacterial delivery to and/or acidification within acidic intracellular compartments rather than as pH-independent measures of bacterial uptake.

#### Phagosomal acidification assay using high-content imaging

Phagosomal acidification was assessed using an established pHrodo-based fluorescence imaging approach^112^. Briefly, WT and GIV-KO THP1 cells were seeded at 5 × 10⁴ cells/well in black, clear-bottom 96-well imaging plates in RPMI supplemented with 10% FBS and PSG and differentiated with 25 nM PMA overnight. The following day, medium was replaced with RPMI containing 0.2% FBS, and cells were cultured overnight. On the day of infection, cells were washed and maintained in phenol red-free RPMI supplemented with 10% FBS.

Heat-inactivated, FITC-conjugated *AIEC* LF82 were additionally labeled with pHrodo Red-SE (Thermo Fisher Scientific). pHrodo Red-SE was added to the bacterial suspension at a final concentration of 0.1 mg/mL and incubated for 30–45 min at room temperature with gentle rotation, protected from light. Labeled bacteria were washed three times with 1× PBS to remove unbound dye and resuspended in phenol red-free RPMI containing 10% FBS.

For nuclear labeling, cells were incubated with Hoechst 33342 (1:2,000) for 20 min at 37°C before bacterial addition. Cells were then challenged with dual-labeled pHrodo Red/FITC *AIEC* LF82 at an MOI of 50.

Live-cell images were acquired using a Cytation 10 Cell Imaging Multi-Mode Reader (Agilent BioTek), with identical acquisition and analysis parameters across experimental conditions. pHrodo Red fluorescence served as a pH-sensitive readout of bacteria localized within acidic intracellular compartments, whereas FITC fluorescence was used to visualize labeled bacteria. Combined fluorescence imaging enabled assessment of bacterial association and acidification-associated changes.

#### General notes for the synthesis and characterization of IGGi11-Me

Deuterated solvents were purchased from Cambridge Isotope Laboratories, Inc. All ^1^H nuclear magnetic resonance (NMR) spectra were recorded on a Jeol-ECZ spectrometer operating at 400MHz. Chemical shifts were reported in parts per million (ppm) in the scale relative to the residual solvent signals. Multiplicities are abbreviated as: s, singlet; d, doublet; t, triplet; p, pentet. Coupling constants are measured in Hertz (Hz). Low resolution MS analysis was performed on a Micromass Quattro Ultima triple quadrupole mass spectrometer with an electrospray ionization (ESI) source and an atmospheric pressure chemical ionization source (APCI) by Molecular Mass Spectrometry Facility (MMSF) in the department of chemistry and biochemistry at University of California, San Diego. Liquid chromatography–mass spectrometry was obtained on a Waters ACQUITY UPLC M-Class Chromatography System. Preparative HPLC was performed on a Waters LC Prep 150 System using SunFire C18 OBD Prep column (100Å, 19 mm × 50 mm, 5 μM) with water containing 0.05% trifluoroacetic acid (TFA) as eluent A and methanol as eluent B. A linear gradient of 15−95% eluent B is applied at a flow rate of 20 mL/minutes over 28 minutes per run.

#### Synthetic procedure for IGGi11-Me

##### Synthesis of sodium 9H-fluorene-2,7-disulfonate

Chlorosulfonic acid (2.0 mL, 45 mmol, 3 equiv) was added dropwise at 0 °C to a stirred solution of fluorene (1.7g, 10 mmol, 1 equiv) in acetic acid (15 mL). The reaction mixture was heated at 160°C for 16h. After cooling to room temperature, the mixture was poured into 30 mL of an aqueous solution of saturated NaCl containing NaOH (1.0 g, 2.5 equiv). The solid was filtered, washed three times with saturated brine, and dried under vacuum at 60 °C to afford sodium 9H-fluorene-2,7-disulfonate as brown solid (3.4 g, 92%). The compound was taken on to the next step without further purification.

^1^H NMR (400 MHz, D_2_O) δ = 7.88 (s, 2H), 7.77 (d, *J* = 8.1 Hz, 2H), 7.69 (d, *J* = 8.0 Hz, 2H), 3.65 (s, 2H). ESI-MS: 162.41 [M-2Na]^2-^

##### Synthesis of 9H-fluorene-2,7-disulfonyl dichloride

Sodium 9H-fluorene-2,7-disulfonate (1.0 g, 2.75 mmol, 1 equiv), PCl_5_ (1.72 g, 8.25 mmol, 3 equiv) and POCl_3_ (3.0 mL, 33 mmol, 12 equiv) were heated neat to reflux for 16 hours. After cooling to room temperature, the reaction was poured into iced water (300 mL), and the slurry was sonicated to break up the aggregates. The solid was filtered to obtain 9H-fluorene-2,7-disulfonyl dichloride as brown solid (0.82g, 84%). The compound was taken on to the next step without further purification.

^1^H NMR (400 MHz, CDCl3) δ = 8.31 (s, 2H), 8.19 (d, *J* = 8.2 Hz, 2H), 8.12 (d, *J* = 8.3 Hz, 2H), 4.21 (s, 2H). APCI-MS: 361.11[M-H]^-^

##### Synthesis of 4-(methylamino) methyl butanoate

Hydrochloric acid (7 mL) was added to N-methylpyrrolidone (5 g, 50 mmol, 1 equiv) and the mixture was refluxed for 16 h. The HCl was removed under reduced pressure. As the yellow solid starts to crystallize, cold acetone was added. The mixture was stirred thoroughly, put into the freezer overnight to recrystallize. The solid was then filtered, washed two times with cold acetones, and dried in high vacuum to obtain 4-(Methylamino)butanoic acid as white solid. (6g, 100%). The reaction was taken on to the next step without further purification.

Thionyl chloride (2.5 mL, 4.0 g, 34 mmol, 4 equiv) was added dropwise at 0 °C to a stirred solution of 4-(Methylamino) butanoic acid (1.0 g, 8.5mmol, 1 equiv) in dry methanol (15 mL). The solution was allowed to warm to room temperature, stirred for 3 hours, and the solvent was removed under reduced pressure. The residue was washed with a mixture of hexane/DCM (13:1) and dried in high vacuum. 4-(methylamino) methyl butanoate was obtained as grey-white crystal (0.78g, 70%). The reaction was taken on to the next step without further purification.

^1^H NMR (400 MHz, CD_3_OD) δ = 3.70 (s, 3H), 3.06 (t, *J* = 8.0 Hz, 2H), 2.71 (s, 3H), 2.50 (t, *J* = 7.2 Hz, 2H), 1.98 (p, *J* = 8.0, 7.1Hz, 2H)

##### Synthesis of dimethyl 4,4’-((9H-fluorene-2,7-disulfonyl)bis (methylazanediyl)) dibutyrate (IGGi11-Me)

4-(methylamino) methyl butanoate (22 mg, 0.17 mmol, 2 equiv) and N, N-Diisopropylethylamine (43 μL, 0.25 mmol, 3 equiv) were added to a stirred solution of 9H-fluorene-2,7-disulfonyl dichloride (30 mg, 0.08 mmol, 1 equiv) in dimethylamine (1 mL). The mixture was stirred at room temperature overnight, then quenched with 1M HCl (1.2 mL) prior to extraction three times with dichloromethane and washing three times with brine. The crude was dried over Na_2_SO_4_, and the solvent was evaporated in vacuo and purified by column chromatography on silica gel using petroleum ether/ethyl acetate (6:4 to 3:7) as the eluent. Final purification was done by preparative HPLC using a gradient of 15−95% methanol in water containing 0.05% TFA (injection volume is 1mL). The final product IGGi11-Me was obtained as a white solid (10 mg, 22%).

^1^H NMR (400 MHz, CDCl3) δ = 8.01 (s, 2H), 7.97 (d, *J* = 8.2 Hz, 2H), 7.87 (d, *J* = 8.0 Hz, 2H), 4.07 (s, 2H), 3.69 (s, 6H), 3.10 (t, *J* = 6.8Hz, 4H), 2.77 (s, 6H), 2.44 (t, *J* = 7.3Hz, 4H), 1.89 (p, *J* = 7.0Hz, 4H). LC-MS: 553.32 [M+H]^+^ ESI-MS: 553.35 [M+H]^+^

##### Human subjects

Healthy patient-derived organoids (PDOs) were generated from colonic biopsies collected during routine colonoscopies at the University of California, San Diego IBD Center. Patients were enrolled and consented under a research protocol approved by the Institutional Review Board (IRB Project ID# 1132632; PI: Boland) and compliant with the Human Research Protection Program (HRPP). Histologically normal colon tissue was obtained from patients undergoing screening colonoscopy or evaluation for irritable bowel syndrome. All participants provided informed consent for the collection of colonic biopsies to generate 3D organoids.

Isolation and biobanking of organoids were performed under a separate IRB-approved protocol (Project ID# 190105; PI: Ghosh) at the UC San Diego HUMANOID Center of Research Excellence (CoRE).

De-identified clinical data—including age, gender, and disease history—were extracted from medical records in accordance with HIPAA regulations. The study adhered to the ethical principles outlined in the Declaration of Helsinki.

##### Isolation of patient derived enteroids

Intestinal crypts containing crypt-base columnar cells were isolated from human colonic tissue following established protocols^113^. Briefly, tissues were digested with collagenase type I (2 mg/mL) supplemented with gentamicin (50 μg/mL) at 37°C in a CO₂ incubator. Every 10 minutes, samples were vigorously pipetted and monitored by light microscopy to assess release of epithelial units. Digestion was stopped by adding wash media (DMEM/F12 with HEPES and 10% FBS). The cell suspension was filtered through a 70 μm strainer, centrifuged at 200 × g for 5 minutes, and the supernatant discarded. Viable intestinal stem cells were quantified using Trypan Blue exclusion with a DeNovix Celldrop cell counter. Epithelial units were resuspended in Matrigel, and 25 μL of the cell–Matrigel mixture was plated per well of a 12-well plate on ice. Plates were inverted and incubated at 37°C for 10 minutes to allow Matrigel polymerization. Epithelial units were resuspended in Matrigel, and 25 μL of the cell–Matrigel mixture was plated per well of a 12-well plate on ice. Plates were inverted and incubated at 37°C for 10 minutes to allow Matrigel polymerization.

Following gelation, 1,000 μL of 50% conditioned medium (purchased from the UC San Diego HUMANOID Center; Intestigro; Cat#HUM2019), prepared from L-WRN cells^113^ [ATCC CRL-3276] which produce biologically active Wnt3a and R-Spondin. This ‘base media’ was supplemented with 20 ng/mL EGF, 10 μM SB202190 (p38 MAPK inhibitor), 10 μM Y27632 (ROCK inhibitor), and 10 μM SB431542 (TGFβ/SMAD inhibitor). Media were refreshed every 2 days, and enteroids were expanded and biobanked for subsequent use.

##### Creation of gut-in-a-dish model (microbe-organoid-macrophage co-cultures)

EDMs were prepared using single cells dissociated from enteroids, seeded onto 24-well transwells with 1 μm pore polyester membranes precoated with diluted Matrigel (1:40 in cold PBS) in 5% conditioned media, as previously described. Cells were seeded at ∼2x10^5^ cells/well and differentiated for 24 hours in 5% conditioned media, prepared by diluting Intestigro with Tailor-2-Gro (HUMANOID Center; cat# HUM202402).

Simultaneously, WT and GIV KO THP1 monocytes were seeded onto a 6-well tissue culture plate at 1e^6^ cells/mL density in RPMI, supplemented with 10% FBS and 25nM PMA to induce macrophage differentiation overnight. After 24hours, THP1 monocyte-derived macrophages were detached via trypsin and placed basolaterally (1e^5^ cells/well) on the flipped trans well membrane to incubate for 2 hours. Once set, trans wells were flipped again with epithelial monolayer media applied apically, and RPMI+10%FBS applied basolaterally to feed the co-culture model.

Adherent invasive *Escherichia coli* strain LF82 was cultured aerobically in LB broth for 5 hours, then sub-cultured overnight anaerobically for 14 hours. Cocultures were infected apically at a multiplicity of infection (MOI) of 1:30 (epithelial cells: *AIEC* LF82) for 36 hours, with transepithelial electrical resistance (TEER) monitored throughout.

##### Assessment of trans-epithelial electrical resistance (TEER)

TEER was measured manually using STX2 electrodes and an EVOM2 volt/ohm meter (World Precision Instruments) on 24-well transwell plates. Measurements were taken the night before infection and at 6-, 24-, and 36-hours post-infection. Raw TEER values (ohms) were normalized by multiplying by the transwell surface area (0.33 cm²).

## Ethics Statement

All animal studies were conducted in strict accordance with the NIH *Guide for the Care and Use of Laboratory Animals* and were approved by the Institutional Animal Care (#S17223; PI Ghosh). Healthy colon patient-derived organoids (PDOs) were obtained from UC San Diego HUMANOID™ Center, a biorepository and research center dedicated to organoid-based disease modeling. Colon PDOs were derived from individuals undergoing routine colonoscopy for cancer screening at UC San Diego, with informed consent under an IRB-approved protocol (#190105; PI Ghosh).

### 3. Quantification and statistical analysis

All images were processed using ImageJ, Gen5, LAS X, or iStudio (LiCOR, Lincoln, NE, USA) software, and assembled into figure panels using Adobe Photoshop and Illustrator (Adobe Creative Cloud). All graphs were generated using GraphPad Prism v9.3.1. Unless otherwise indicated, experiments were performed with at least three independent biological replicates, and data are presented as mean ± standard error of the mean (SEM). The exact sample size (n), definition of n (e.g., independent biological replicates, individual mice, or independently analyzed samples), statistical tests used, and any deviations from the stated data presentation are specified in the corresponding figure legends. Comparisons between two groups were performed using unpaired Student’s t-tests, unpaired t-tests with Welch’s correction, or the nonparametric Mann–Whitney U test, as appropriate. Comparisons involving multiple groups or experimental factors were performed using one-way or two-way analysis of variance (ANOVA), followed by Tukey’s multiple-comparisons test, as indicated in the corresponding figure legends. P ≤ 0.05 was considered statistically significant. Phosphoproteomic data were analyzed using Welch’s *t*-tests and Benjamini–Hochberg multiple-testing correction as described under Phosphoproteomics data processing and visualization. Additional details of the statistical analyses are provided in the corresponding figure legends.

## SUPPLEMENTAL INFORMATION INDEX

### Supplemental videos (Videos S1-S2)

Video S1. Visualization of pHrodo signal indicating phagolysosomal acidification upon *AIEC* LF82 (Green) infection in GIV-WT THP1 macrophages, related to Figure 4.

Video S2. Visualization of pHrodo signal indicating failure of phagolysosomal acidification upon *AIEC* LF82 (Green) infection in GIV-KO THP1 macrophages, related to Figure 4.

