## Supplementary text for "A NOD2-Encoded Toggle Switch Resolves the Host–Microbe Battle Over Cyclic AMP Control"

### SUPPLEMENTAL INFORMATION

#### CATALOG OF SUPPLEMENTAL INFORMATION


1. *Supplementary Figures and Legends (S1-S8)*
2. *Supplementary Tables (S1-S6)*
3. *Supplementary References (9)*

#### SUPPLEMENTARY FIGURES AND LEGENDS

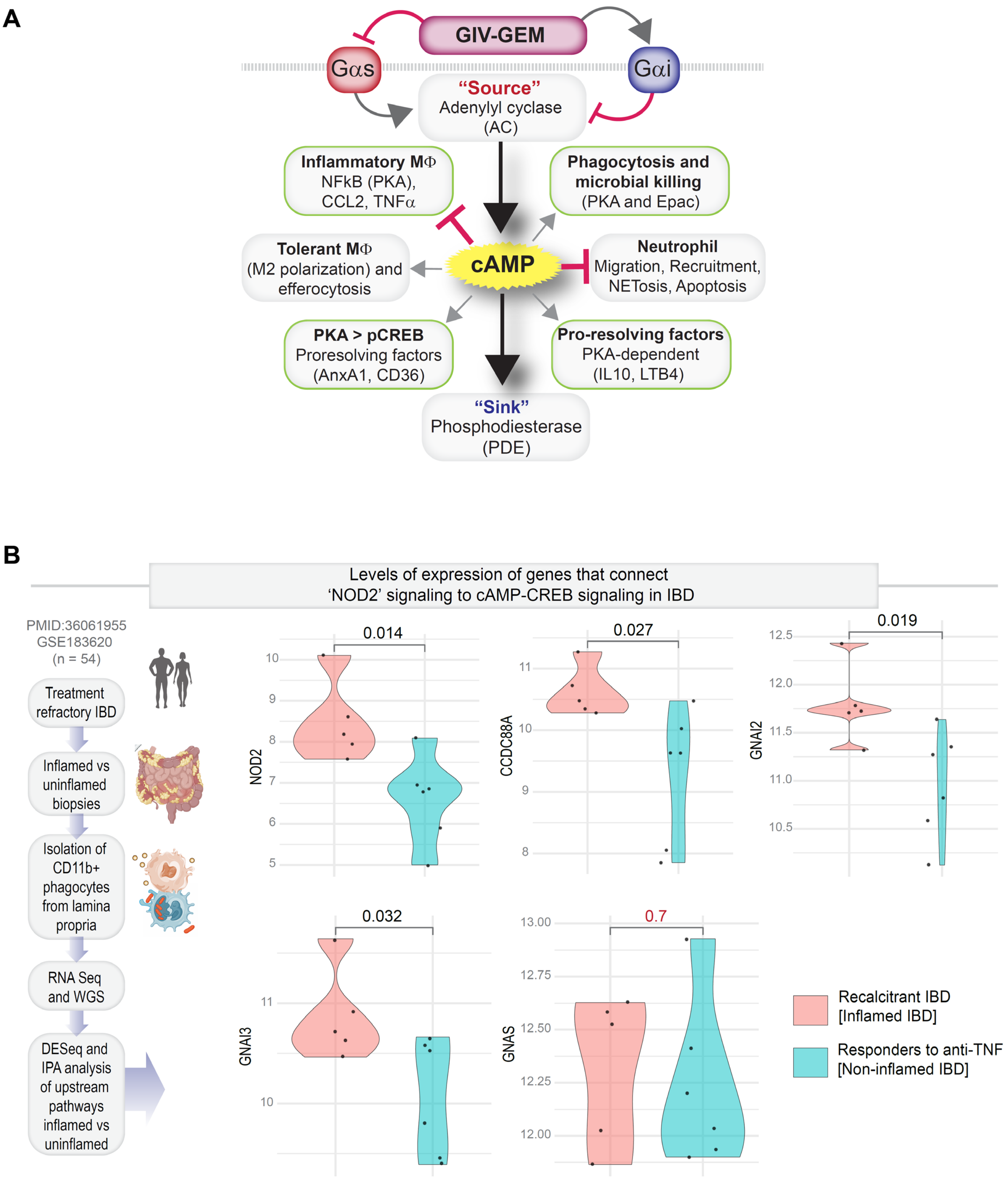


**Figure S1. Patient transcriptome reveals *CCDC88A* as a node linking NOD2 to cAMP signaling in IBD, related to Figure 1**

**A.** Schematic summarizing major sources, sinks, regulators, and immune consequences of cyclic AMP (cAMP) during infection [References S1,2]. Adenylyl cyclase (AC) generates cAMP, whereas phosphodiesterases (PDEs) degrade it. GIV, a prototypical guanine-nucleotide exchange modulator (GEM), suppresses cAMP [References S3-5] through its conserved C-terminal GEM motif by activating inhibitory Gαi [Reference S6] and inhibiting stimulatory Gαs [Reference S7].

**B.** TPM-normalized expression of NOD2, CCDC88A, GNAI2, GNAI3, and GNAS in CD11b$^{+}$ lamina propria cells from patients with inflammatory bowel disease (IBD), stratified as anti-TNF-responsive/non-inflamed (teal) or treatment-refractory/inflamed (salmon). Violin plots show the distribution of expression values; points indicate individual samples. Exact P values are shown above comparisons and were calculated Welch's t-test. The source dataset and analytical workflow are depicted at left; corresponding IPA upstream-pathway analysis is shown in **Figure 1B**.

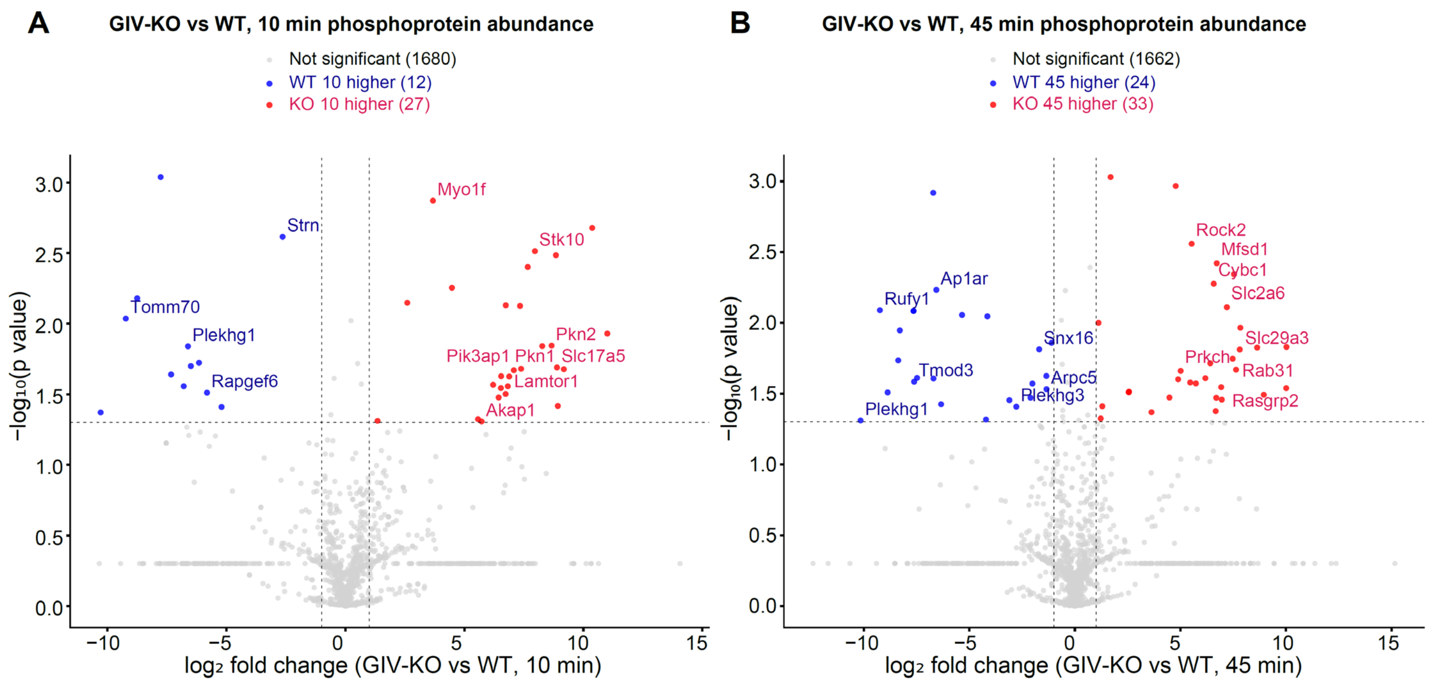


**Figure S2. Unbiased phosphoproteomic profiling reveals time-dependent signaling rewiring following loss of GIV, related to Figure 1**

**A–B.** Volcano plots comparing protein-level phosphorylation-associated abundance between GIV-knockout (KO) and wild-type (WT) thioglycollate-elicited peritoneal macrophage (TGPMs) following adherent-invasive *Escherichia coli* (*AIEC*) LF82 infection for **10 min (A)** and **45 min (B)**. The x-axis represents $\log_2$ fold change of phosphoprotein abundance (GIV-KO/WT), and the y-axis represents $-\log_{10}$ *P* value. Proteins with greater phosphorylation-associated abundance in GIV-KO macrophages are shown in red, those with greater abundance in WT macrophages are shown in blue, and proteins not meeting the indicated exploratory thresholds are shown in gray. Selected proteins are labeled to illustrate representative genotype-dependent differences. Dashed vertical and horizontal lines indicate the exploratory classification thresholds of |$\log_2$ fold change| +/- 1 and nominal $P < 0.05$. See **Tables S1–S5** for the complete time- and genotype-resolved phosphoproteomic results. Each point represents a protein-level phosphorylation-associated signal aggregated from detected phosphopeptides and should not be interpreted as total protein abundance or phosphorylation of an individual site. Biological replicates were n = 2/genotype/time point; therefore, nominal P-value-based classifications are exploratory and hypothesis-generating. Benjamini–Hochberg-adjusted results and complete protein-level values are provided in **Tables S1–S5**.

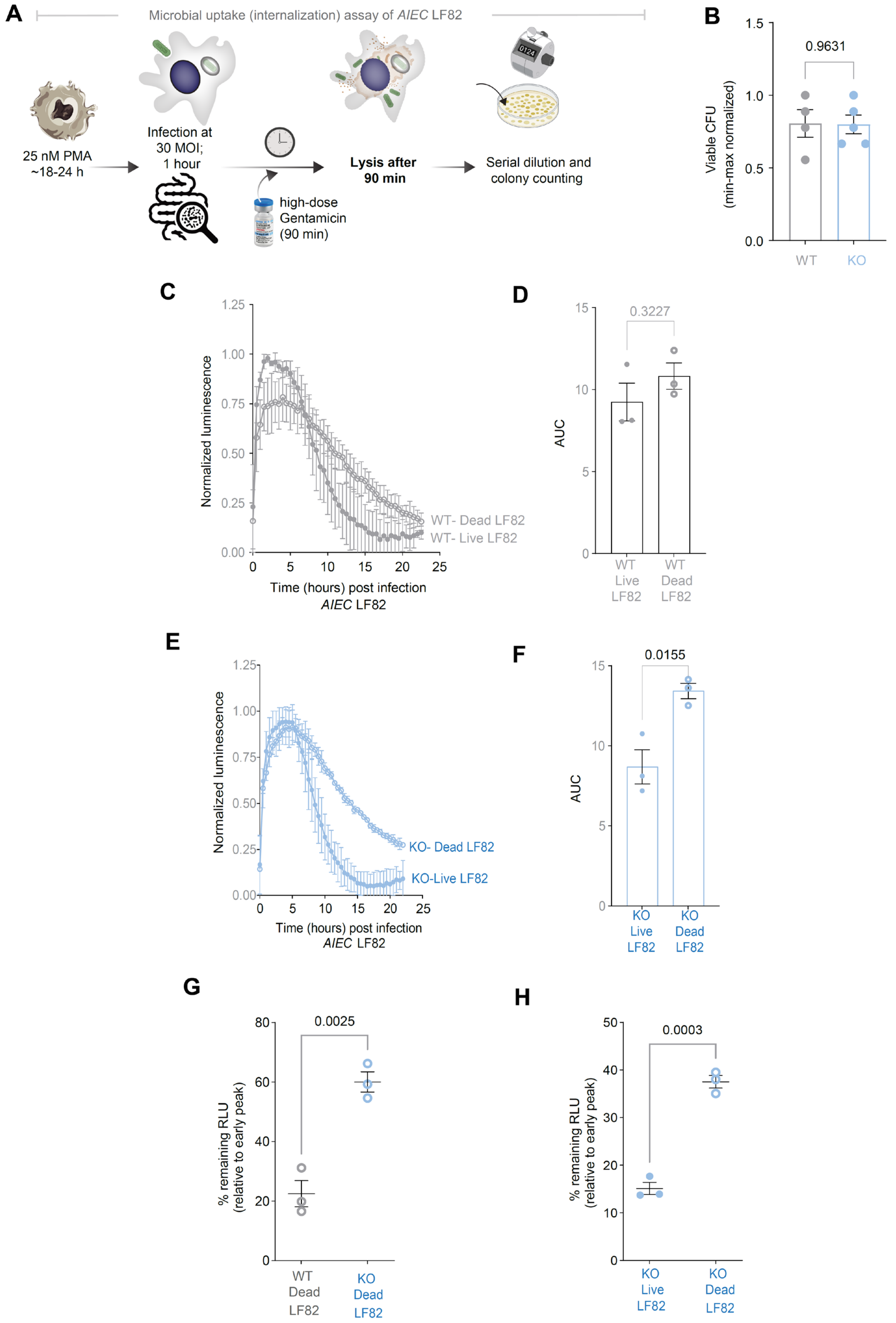


**Figure S3. GIV is dispensable for initial bacterial internalization but shapes pathogen-induced cAMP–CREB dynamics, related to Figure 2.**

**A-B.** Workflow (A) and quantification (B) of bacterial internalization in PMA-differentiated WT and GIV-KO THP-1 macrophages. Cells were differentiated with 25 nM PMA for ~18–24 h, infected with *AIEC* LF82 at MOI 30 for 1 h, and extracellular bacteria were eliminated with high-dose gentamicin for 90 min. Cells were then lysed, and viable intracellular bacteria were quantified by serial dilution and colony-forming unit (CFU) counting. In B, CFU values were min–max normalized across the displayed samples within each biological replicate. Individual symbols represent biological replicates; bars show mean ± SEM.

**C-F**. CREB-luciferase responses to live or heat-inactivated (“dead”) *AIEC* LF82, displayed separately within each genotype to resolve the contribution of microbial viability. Time courses show normalized CREB-dependent luminescence in WT (C) and GIV-KO (E) reporter macrophages; corresponding area-under-the-curve (AUC) values are shown in D and F, respectively. Within each genotype, luminescence was normalized to permit comparison of response kinetics between live and heat-inactivated bacteria. Lines and error bars indicate mean ± SEM; individual symbols in D and F represent independent biological replicates. See **Figure 2G–I** for direct cross-genotype comparisons.

**G-H.** Persistence of CREB signaling during the late phase, expressed as the percentage of relative luminescence units (RLU) remaining at the late time point (~20 h) relative to the early response peak for each condition. G compares WT and GIV-KO macrophages challenged with heat-inactivated *AIEC* LF82; H compares live versus heat-inactivated *AIEC* LF82 within GIV-KO macrophages. Higher values indicate less complete termination of the early CREB response.

*Statistics:* Data are mean ± SEM; individual symbols represent independent biological replicates (n = 4 for panel B; n=3 for Panel D,F,G and H). Statistical significance was assessed using unpaired *t*-tests for panels D, F, G, and H and an unpaired t-test with Welch’s correction for panel B. Exact *P* values are indicated above the corresponding comparisons. $P \leq 0.05$ was considered significant.

*Abbreviations*: CREB, cAMP response element-binding protein; RLU, relative luminescence units; AUC, area under the curve; *AIEC*-LF82, adherent-invasive Escherichia coli; MOI, multiplicity of infection; CFU, colony-forming units; WT, wild type; KO, GIV knockout.

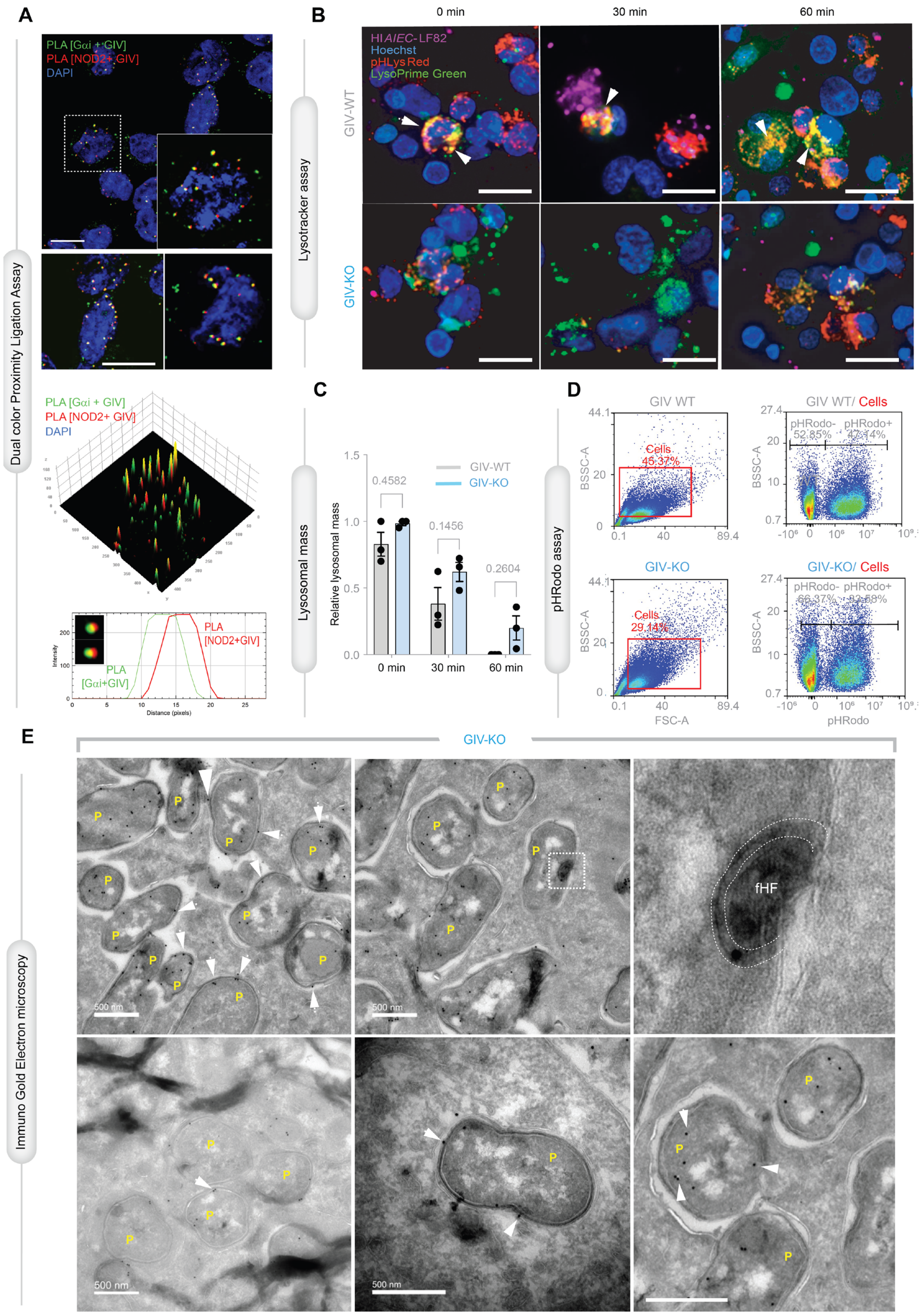


**Figure S4. GIV coordinates spatial NOD2/Gαi signaling and promotes lysosomal acidification and phagolysosomal maturation during *AIEC* LF82 infection, related to Figures 3 and 4**

**A.** Sequential dual-color proximity ligation assay (PLA; upper panel) resolving endogenous Gαi•GIV (green) and NOD2•GIV (red) complexes within the same cell following MDP (10 µg/ml, 1 h) stimulation. DAPI (blue) marks nuclei. Yellow puncta represent spatial overlap of red and green PLA signals, not a distinct PLA species. **Inset**: Representative foci. Three-dimensional intensity mapping (lower panel) depicts the spatial distribution and relative intensity of individual red and green PLA puncta. The line scan across a representative overlapping punctum shows fluorescence intensity as a function of distance (pixels) for Gαi•GIV (green) and NOD2•GIV (red), illustrating their spatial proximity. Scale bars, 10 µm. See **Figure 3G-K** for quantitative analysis of partner exchange.

**B-C**. Live-cell assessment of lysosomal acidification and relative lysosomal mass in GIV-WT and GIV-KO THP1 macrophages infected with *AIEC* LF82. B, representative images at 0, 30, and 60 min after infection. *AIEC* LF82 is shown in magenta; pHLys Red, an acidity-sensitive lysosomal probe, in orange/red; LysoPrime Green, a relatively pH-insensitive lysosomal-mass probe, in green; and Hoechst nuclear stain in blue. White arrowheads indicate bacteria-containing acidic compartments showing accumulation/overlap of the bacterial and lysosomal signals. Scale bars, 100 µm C, relative lysosomal mass derived from LysoPrime Green fluorescence and min–max normalized across the displayed measurements for comparative visualization; these values do not represent absolute lysosomal mass. Individual symbols represent independent biological replicates; bars show mean ± SEM (n = 3). Exact *P* values are shown and were determined by two-way ANOVA followed by Tukey's multiple-comparisons test. Corresponding lysosomal acidification indices (pHLys Red/LysoPrime Green) are quantified in **Figure 4C-D.**

**D**. Representative flow-cytometric gating for pHrodo-labeled *AIEC* LF82 in GIV-WT and GIV-KO THP1 macrophages. Cells were first identified on forward-scatter area (FSC-A) versus side-scatter area (SSC-A) plots (red gates; percentage of total acquired events indicated), followed by quantification of pHrodo-negative (pHrodo$^{-}$) and pHrodo-positive (pHrodo$^{+}$) populations within the cell gate. Percentages above the pHrodo gates indicate the fraction of gated cells within each population. Because pHrodo fluorescence increases in acidic environments, this assay reports bacterial delivery to and/or acidification within acidic intracellular compartments and is not interpreted as a pH-independent measure of bacterial uptake. Quantitative median fluorescence intensity and pHrodo$^{+}$ cell frequencies are shown in **Figure 4E–F**.

**E.** Representative immunogold electron microscopy of *AIEC* LF82-infected GIV-KO primary murine peritoneal macrophages. **P** denotes pathogen-containing compartments; **HF**, hemifusome; **fHF**, flipped hemifusome; and MVB, multivesicular body. White arrowheads indicate 12-nm gold particles labeling NOD2; GIV was detected with 18-nm gold particles as described in **Figure 4H**. The dashed box identifies the region enlarged in the adjacent high-magnification image; dashed outlines delineate the fHF structure. Scale bars are 500 nm where indicated. Representative GIV-KO fields illustrate the reduced frequency of hemifusome-associated structures quantified relative to WT in **Figure 4H**.

*Statistics:* Data are mean ± SEM; individual symbols represent independent biological replicates (n = 3, as shown). Statistical significance was assessed using two-way ANOVA for panel C; Exact *P* values are indicated above the corresponding comparisons. $P \leq 0.05$ was considered significant.

*Abbreviations*: PLA, proximity ligation assay; DAPI, 4′,6-diamidino-2-phenylindole; *AIEC*-LF82, adherent-invasive *Escherichia coli*; WT, wild type; KO, GIV knockout; FSC-A, forward-scatter area; SSC-A, side-scatter area; HF, hemifusome; fHF, flipped hemifusome; MVB, multivesicular body.

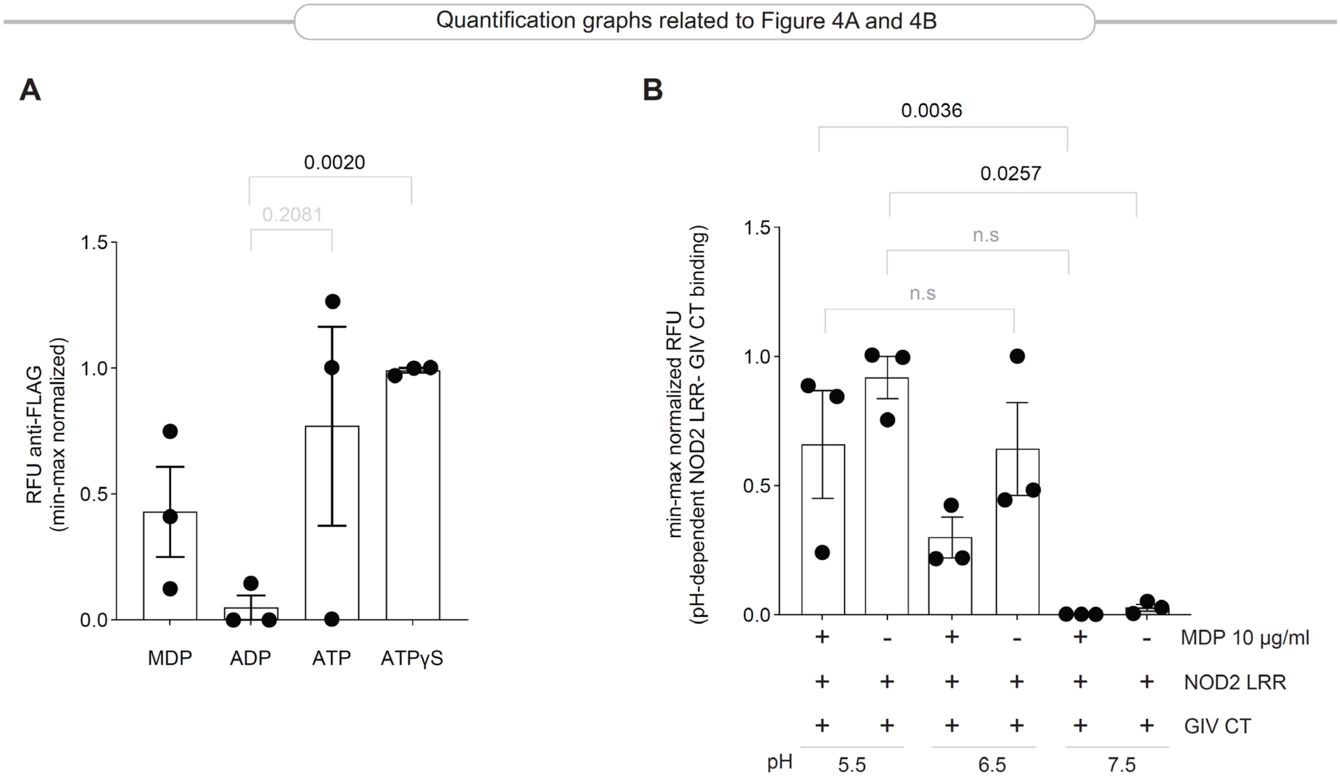


**Figure S5. Nucleotide state and acidic pH favor NOD2•GIV complex formation, related to Figure 4**

**A.** Densitometric quantification of nucleotide-dependent NOD2•GIV association corresponding to **Figure 4A**. GIV binding to NOD2 was quantified following incubation with MDP, ADP, ATP, or the non-hydrolysable ATP analog ATPγS and normalized by min–max scaling. Both ATP and ATPγS enhanced NOD2•GIV association relative to ADP. The hydrolysable nucleotide ATP showed greater inter-experimental variability, consistent with sampling a dynamic ensemble of nucleotide-dependent conformational states during binding and hydrolysis, whereas non-hydrolysable ATPγS produced a more consistent increase, consistent with stabilization of a pre-hydrolytic nucleotide-bound state. Individual symbols represent independent experiments (n = 3); bars show mean ± SEM. Statistical significance was assessed using Welch's t-test; indicated P values are shown.

**B.** Densitometric quantification of pH-dependent NOD2•GIV binding corresponding to **Figure 4B**. Recombinant NOD2-LRR and GIV-CT were incubated at the indicated pH (5.5, 6.5, or 7.5) in the presence or absence of MDP. GIV-CT binding was quantified and normalized by min–max scaling. NOD2•GIV association was favored at acidic pH and markedly reduced at pH 7.5 quantified relative to WT in **Figure 4H**.

*Statistics:* Data are mean ± SEM; individual symbols represent independent biological replicates (n = 3, as shown). Statistical significance was assessed using one-way ANOVA, followed by Tukey's multiple-comparisons test for all comparisons; Exact *P* values are indicated above the corresponding comparisons. $P \leq 0.05$ was considered significant. ns, not significant.

***Abbreviations:*** MDP, muramyl dipeptide; ADP, adenosine diphosphate; ATP, adenosine triphosphate; ATPγS, adenosine 5′-[γ-thio] triphosphate; RFU, relative fluorescence units; NOD2- LRR, NOD2 leucine-rich repeat domain; GIV CT, GIV C-terminal domain;

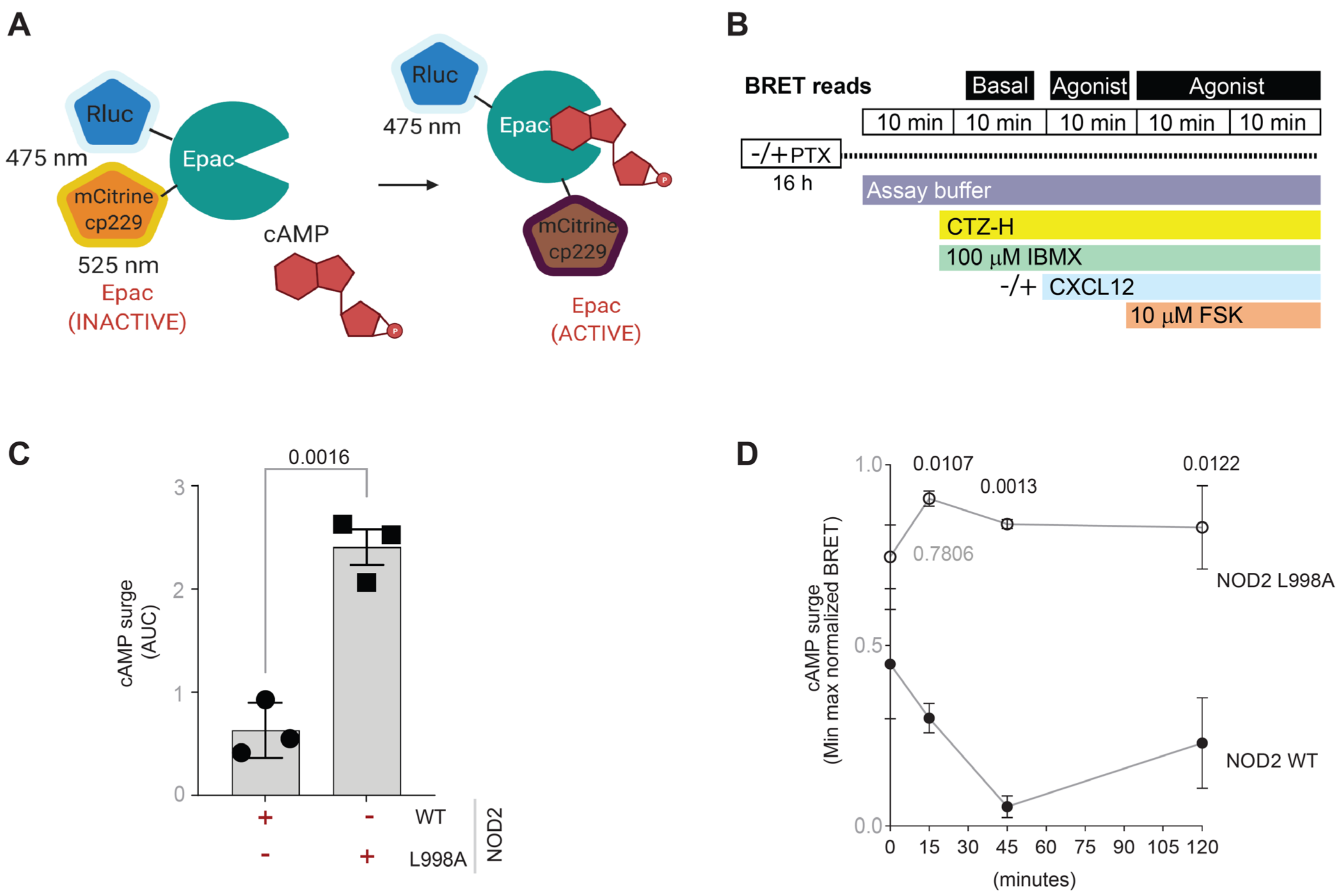


**Figure S6. Disrupting the NOD2•GIV interface uncouples NOD2 sensing from the GIV•Gαi-dependent cAMP plunge, related to Figure 5**

**A-B.** Schematic and workflow of the CAMYEL bioluminescence resonance energy transfer (BRET)-based cAMP reporter assay. **A**, CAMYEL (cAMP sensor using YFP–Epac–RLuc) contains *Renilla* luciferase (RLuc) and the fluorescent acceptor mCitrine cp229 linked through the cAMP-binding domain of Epac (exchange protein directly activated by cAMP). Binding of cAMP induces a conformational change that decreases BRET; thus, **lower BRET corresponds to higher intracellular cAMP**. **B**, general assay workflow and pharmacologic controls used to validate dynamic range and Gαi-dependent cAMP regulation. CTZ-H (coelenterazine-H) is the RLuc substrate; IBMX (3-isobutyl-1-methylxanthine) inhibits phosphodiesterases; forskolin (FSK) activates adenylyl cyclase; CXCL12 serves as a canonical Gαi-coupled GPCR agonist; and pertussis toxin (PTX) inhibits Gαi signaling. These controls establish reporter performance and are shown as the general CAMYEL assay framework; the NOD2 experiments in C–D instead used muramyl dipeptide (MDP) stimulation and did not use CXCL12/PTX as experimental stimuli.

**C**. Integrated quantification (AUC) of the CAMYEL cAMP responses shown in **Figure 5J**, comparing cells expressing NOD2-WT or the GIV-binding-deficient NOD2-L998A mutant. Individual symbols represent independent biological replicates (n = 3); bars show mean ± SEM.

**D**. Time-resolved analysis of the same NOD2-WT and NOD2-L998A responses, min–max normalized to resolve response kinetics. Whereas NOD2-WT supports an early cAMP surge followed by a late plunge, NOD2-L998A retains elevated cAMP, indicating that disruption of the NOD2•GIV interface selectively uncouples NOD2 sensing from termination of the cAMP response. See **Figure 5J** for the corresponding primary CAMYEL analysis.

*Statistics:* Data are mean ± SEM; individual symbols represent independent biological replicates (n = 3 as shown). Statistical significance was assessed using unpaired t-test with Welch's correction for panel C and two-way ANOVA followed by Tukey's multiple-comparisons test for panel D. Exact P values are indicated above the corresponding comparisons. $P \leq 0.05$ was considered significant.

*Abbreviations:* CAMYEL, cAMP sensor using YFP–Epac–RLuc; BRET, bioluminescence resonance energy transfer; RLuc, *Renilla* luciferase; Epac, exchange protein directly activated by cAMP; CTZ-H, coelenterazine-H; IBMX, 3-isobutyl-1-methylxanthine; FSK, forskolin; PTX, pertussis toxin; CXCL12, C-X-C motif chemokine ligand 12; MDP, muramyl dipeptide; AUC, area under the curve; WT, wild type.

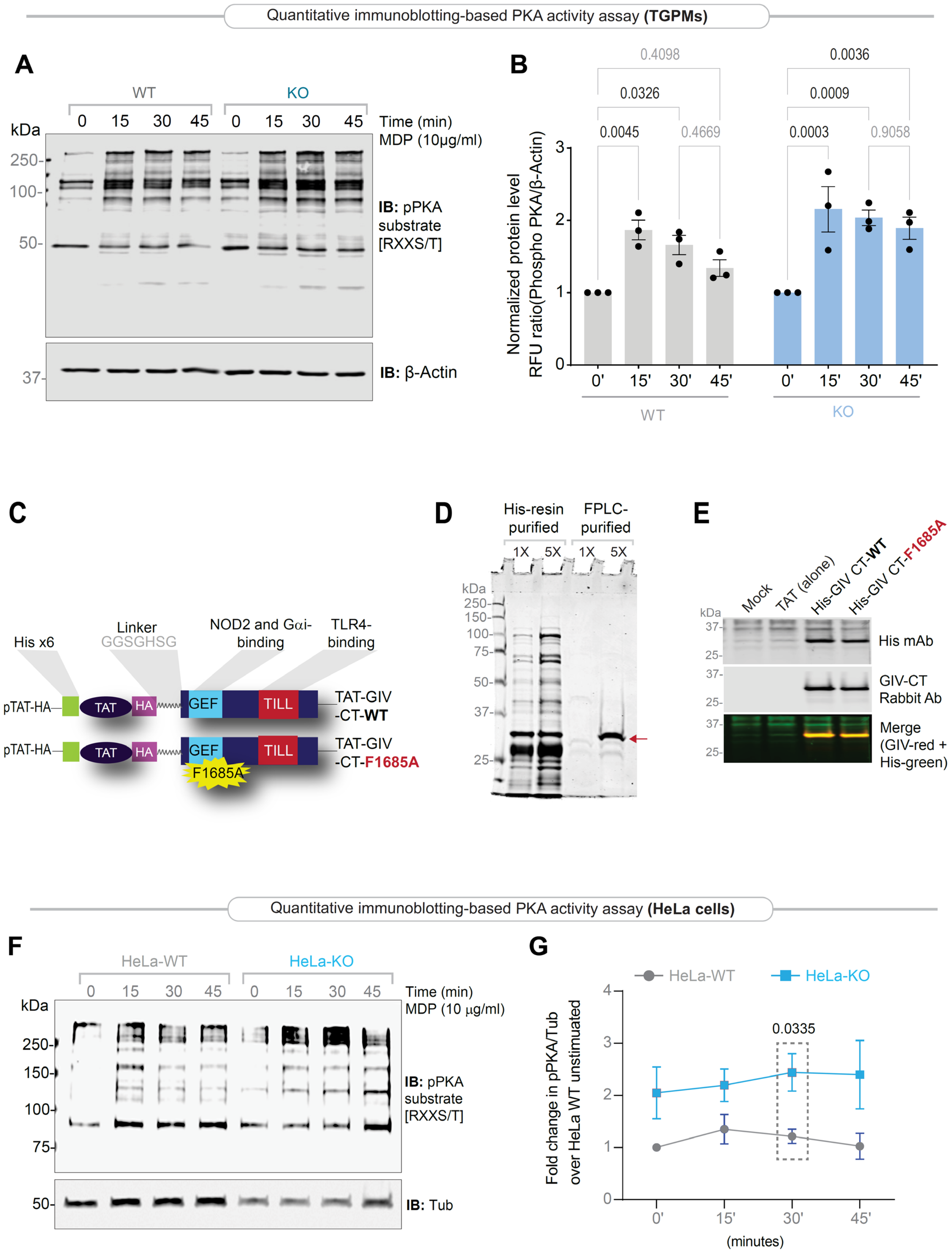


**Figure S7. Functional dissection of GIV's role in regulating PKA activity, related to Figure 6.**

**A-B.** Time-resolved analysis of PKA-dependent substrate phosphorylation in WT and GIV-KO thioglycollate-elicited peritoneal macrophages (TGPMs) following challenge with live *AIEC* LF82 for 0, 15, 30 and 45 min after MDP pre-treatment (10 µg/ml). A, representative immunoblot (IB) using an antibody recognizing phosphorylated PKA consensus substrate motifs [RRXS*/T*, where S*/T* denotes the phosphorylated Ser/Thr residue]. β-actin serves as loading control. B, densitometric quantification of total phospho-PKA-substrate signal, normalized to β-actin and expressed relative to the respective 0-min baseline. Individual symbols represent independent biological replicates (n = 3); bars show mean ± SEM. Exact *P* values are indicated. Same data is visualized as time-resolved line graphs in **Figure 6C**.

**C.** Domain map of cell-permeant TAT–GIV C-terminal (GIV-CT) constructs used for rescue experiments. Recombinant proteins contain an N-terminal polyhistidine ($His_6$) tag, TAT protein-transduction domain, hemagglutinin (HA) tag, and the GIV C-terminal region containing the GEM/guanine-nucleotide exchange factor (GEF) motif and TILL region. WT TAT–GIV-CT retains Gαi-binding/GEF activity, whereas TAT–GIV-CT-F1685A contains Phe1685-to-Ala(A) substitution (yellow star) that disables GEM-dependent Gαi activation while preserving the remainder of the construct. Indicated regions mediating NOD2/Gαi and TLR4 interactions are shown [Reference S8].

**D.** Purification of recombinant His–TAT–GIV-CT. Coomassie-stained gels show His-resin-purified material before and after fast protein liquid chromatography (FPLC). 1× and 5× indicate relative amounts of protein loaded for visualization. The red arrowhead marks the migration position of purified His–TAT–GIV-CT.

**E.** Immunoblot validation of cellular uptake of purified TAT–GIV-CT proteins. Mock and TAT-alone samples serve as negative controls; His–GIV-CT-WT and His–GIV-CT-F1685A were detected with an anti-His monoclonal antibody (His mAb; upper panel) and an antibody recognizing the GIV C terminus (GIV-CT; lower panel). The merged image overlays GIV-CT and His-tag detection channels, confirming identity of the recombinant proteins.

**F-G.** Time-resolved PKA-substrate phosphorylation following MDP stimulation (10 µg/mL) in HeLa-WT and GIV-deficient HeLa-KO cells (used here as orthogonal validation for TGPM studies in **Figure 6C**). F, representative immunoblot of phospho-PKA-substrate motifs [RRXS*/T*], with α-tubulin as loading control. G, densitometric quantification of total phospho-PKA-substrate signal normalized to α-tubulin and expressed as fold change relative to the WT unstimulated condition.

*Statistics*: Data are mean ± SEM; individual symbols represent independent biological replicates (n = 3). Statistical significance was assessed using two-way ANOVA followed by Tukey's multiple-comparisons test to evaluate genotype/time-dependent differences in Panel B, and unpaired Students' t -test for panel G. Exact *P* values are indicated above the corresponding comparisons; $P \leq 0.05$ was considered significant.

*Abbreviations:* PKA, protein kinase A; TGPM, thioglycollate-elicited peritoneal macrophage; MDP, muramyl dipeptide; IB, immunoblot; RFU, relative fluorescence units; TAT, trans-activator of transcription protein-transduction domain; HA, hemagglutinin; His, polyhistidine; GEM, guanine-nucleotide exchange modulator; GEF, guanine-nucleotide exchange factor; TILL, TIR-like BB-loop; FPLC, fast protein liquid chromatography; WT, wild type; KO, knockout.

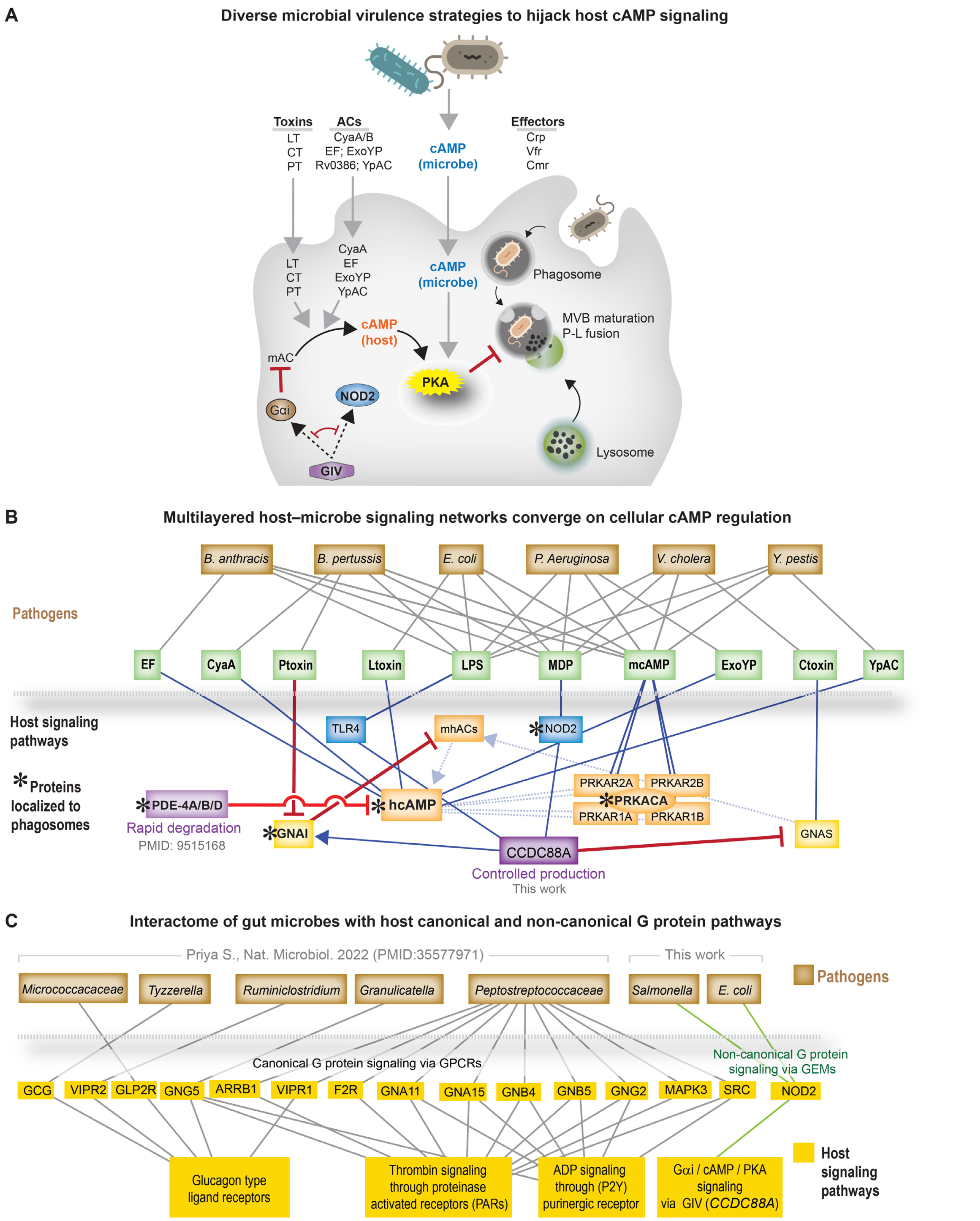


**Figure S8. Summary and working model: Dynamic GIV complex switching shapes microbial sensing and clearance**

**A**. Schematic summarizing diverse microbial mechanisms that elevate host cAMP and the NOD2–GIV–Gαi pathway that counteracts sustained cAMP–PKA signaling.

**B**. Multilayered networks connect specific gut microbes (brown) and their cAMP-modulating microbial genes (green) to host proteins (PRRs, signaling molecules), highlighting disease-specific host pathways regulating cellular cAMP. Strong feedback inhibitory loops (red) are orchestrated by proteins that are localized to phagophore membranes (* asterisk).

**C**. Multilayered networks linking gut microbes (brown) to host genes reveal both canonical GPCR-dependent shown before [Reference S9] and non-canonical GIV-dependent (this work) G protein signaling pathways enriched for disease-specific host pathways in IBD.

#### SUPPLEMENTARY TABLES

**Table S1: Phosphoproteins preferentially elevated in WT macrophages at 10 min after *AIEC* LF82 infection, related to Figure 1E-F.**

| Gene | PKA substrate | Phagosome / lysosome association | Assessment |
|---|---|---|---|
| **Ddx42** | — | — | RNA helicase with roles in RNA processing and nuclear/cytoplasmic RNP biology; no strong phagosome/lysosome connection. |
| **Tomm70** | possible PKA-linked mitochondrial biology | MITOCHONDRIAL — strong, not phagosome | Outer mitochondrial membrane receptor of the TOM complex controlling mitochondrial protein import. Strong mitochondrial compartment marker rather than lysosomal/phagosomal protein. |
| **Pdcd11** | — | — | Nucleolar/ribosome-biogenesis protein; no compelling phagosome/lysosome connection. |
| **Reep4** | — | ENDOMEMBRANE — weak/moderate | ER membrane-shaping protein involved in membrane organization; potentially relevant to membrane supply/trafficking but not a canonical phagosome or lysosome component. |
| **Cnot1** | — | — | CCR4–NOT scaffold controlling RNA turnover and transcription; no strong compartment-specific phagosome signal. |
| **Ddx51** | — | — | Nucleolar RNA helicase involved primarily in ribosome biogenesis. |
| **Plekhg1*** | — | PHAGOCYTIC/CYTOSKELETAL — moderate/strong | Dbl-family RhoGEF that drives Rho-dependent actin rearrangement. Rho activity is directly required for efficient FcγR phagocytosis, making PLEKHG1 a credible upstream cytoskeletal regulator rather than a phagosome resident. |
| **Ranbp10** | — | CYTOSKELETAL — weak/moderate | Microtubule-associated signaling/scaffold protein; potential contribution to cytoskeletal trafficking but no strong direct phagosome/lysosome assignment. |
| **Eif4e** | PKA-related only indirectly | — | Major translation-initiation factor regulated principally through mTOR/MNK pathways rather than being a convincing canonical PKA substrate. |
| **Rapgef6*** | — | PHAGOCYTIC SIGNALING — strong functional link | GEF for **Rap1/Rap2**. This is notable because Rap1 is rapidly activated downstream of Fcγ receptors, localizes near forming phagosomes, and is required for efficient macrophage phagocytosis. |
| **Mycbp2** | — | VESICLE/CYTOSKELETAL — weak/moderate | Giant E3 ligase involved in axonal/cytoskeletal and membrane-trafficking regulation; not a canonical phagosome/lysosome protein. |
| **Strn** | PKA/kinase-phosphatase scaffold — noteworthy | CYTOSKELETAL/SIGNALING — moderate | Striatin is the organizing scaffold of the **STRIPAK kinase–PP2A phosphatase complex**. It is not best interpreted as a PKA substrate, but as a spatial kinase/phosphatase signaling organizer. |

**Table S2: Phosphoproteins preferentially elevated in GIV-KO macrophages at 10 min after *AIEC* LF82 infection, related to Figure 1E-F.**

| Gene | PKA substrate? | Phagosome / lysosome? | Assessment |
|---|---|---|---|
| **Dok3** | — | PHAGOCYTE SIGNALING — moderate | Myeloid adaptor expressed prominently in macrophages; regulates DAP12/TLR signaling. More a regulator of phagocyte activation than a phagosome resident. |
| **Macf1** | — | PHAGOCYTIC/CYTOSKELETAL — moderate | Giant actin–microtubule crosslinker; well positioned to coordinate membrane trafficking and cytoskeletal rearrangements needed for engulfment, but not a canonical phagosome resident. |
| **Myo1f** | — | PHAGOSOME — very strong | Directly recruited to Fc receptor adhesions and **phagocytic cups** in macrophages; links plasma membrane to cortical actin and is required for efficient FcR-mediated phagocytosis and cup closure. |
| **Anxa1** | indirect/possible | PHAGOCYTOSIS/EFFEROCYTOSIS — strong functional link | Annexin A1 is strongly associated with macrophage engulfment, resolution and apoptotic-cell clearance; better considered phagocytosis/efferocytosis machinery than a lysosomal resident. |
| **Nktr** | — | — | NK-cell-associated protein with peptidyl-prolyl isomerase-related biology; no strong direct phagosome/lysosome assignment. |
| **Akap1** | PKA SCAFFOLD — very strong | — | Not simply a PKA substrate: **AKAP1 physically anchors PKA** at the outer mitochondrial membrane and creates a localized PKA signaling compartment, including phosphorylation of DRP1. This is potentially much more mechanistically informative than an isolated PKA substrate. |
| **Sf3b1** | — | — | Core spliceosome component; predominantly nuclear. |
| **Axin1** | possible PKA-regulated | — | Signaling scaffold in Wnt/β-catenin and stress pathways; phosphorylation-rich but not a strong canonical PKA hit for this analysis. |
| **Lamtor1** | — | LYSOSOME — very strong | Bona fide lysosomal membrane scaffold and anchor of the Ragulator complex; organizes **Rag GTPase–mTORC1 signaling on late endosomes/lysosomes**. Particularly strong lysosomal hit. |
| **Pik3ap1 / BCAP** | — | PHAGOCYTE SIGNALING — strong | Major myeloid PI3K adaptor connecting immune receptors to PI3K–AKT; directly relevant to macrophage activation and phagocytic responses, although not a resident phagosomal protein. |
| **Q8C708** | uncertain | uncertain | Mouse UniProt identifier requires exact protein mapping before assigning confidently; I would leave this unclassified rather than overinterpret it. |
| **Rbm8a** | — | — | Exon-junction complex/RNA-processing protein; no strong phagosome/lysosome link. |
| **Osbp** | possible PKA-regulated | ENDOMEMBRANE/ LIPID TRAFFICKING — moderate | Cholesterol/PI4P lipid-transfer protein operating at membrane-contact sites; relevant to organelle lipid homeostasis but not specifically lysosomal. |
| **Pkn1** | — | CYTOSKELETAL/RHO — moderate | Rho-regulated serine/threonine kinase affecting actin organization and membrane dynamics; plausible phagocytic machinery but not a phagosome resident. |
| **Cdk4** | — | — | Cell-cycle kinase; no compelling phagosome/lysosome relationship. |
| **Prdx1** | — | PHAGOCYTE REDOX — moderate | Cytosolic antioxidant enzyme relevant to ROS handling in macrophages; functionally compatible with phagocytic oxidative biology but not a phagosome resident. |
| **Slc28a2** | — | — | Concentrative nucleoside transporter primarily associated with plasma membrane transport rather than lysosomes. |

| | | | |
|---|---|---|---|
| **Lrrc25** | — | INNATE/ENDOLYSOSOMAL — moderate/strong | Myeloid-enriched innate immune regulator linked to turnover of signaling proteins through autophagic/lysosomal pathways; fits inflammatory endolysosomal regulation more than structural phagosome machinery. |
| **Stk10** | no established PKA | CYTOSKELETAL — moderate | STE20-family kinase regulating actin organization, migration and immune-cell morphology; potentially relevant to phagocytic cytoskeletal dynamics. |
| **Psma3** | — | — | Proteasome core component; proteasomal rather than lysosomal degradation. |
| **Pkn2** | — | CYTOSKELETAL/RHO — moderate | Rho-regulated kinase controlling actomyosin and membrane dynamics; plausible contributor to engulfment machinery but not a canonical phagosome protein. |
| **Phf6** | — | — | Nuclear chromatin/transcription regulator. |
| **Slc17a5 / Sialin** | — | LYSOSOME — very strong | Bona fide **lysosomal membrane transporter** exporting free sialic acid and other acidic metabolites from the lysosome. One of the cleanest lysosomal proteins in the set. |
| **Cep170b** | — | — | Centrosome/microtubule-associated protein; possible cytoskeletal relevance but insufficient evidence for phagosome assignment. |
| **Rcc1** | — | — | Nuclear Ran-GEF controlling nucleocytoplasmic transport/mitosis. |
| **Slk** | — | CYTOSKELETAL — moderate | STE20-like kinase regulating focal adhesions, actin and cell motility; indirect fit with phagocytic membrane remodeling. |
| **Faf1** | no established PKA | — | Apoptosis/NF-κB-associated phosphoprotein; no compelling phagosome/lysosome localization. |

**Table S3: Phosphoproteins preferentially elevated in WT macrophages at 45 min after *AIEC*-LF82 infection**, related to Figure 1E-F

| Gene | PKA substrate | Phagosome / lysosome association | Assessment |
|---|---|---|---|
| **Plekhg1** * | — | PHAGOCYTIC/CYTOSKELETAL — moderate | RhoGEF controlling Rho-family signaling and actin remodeling; plausible phagocytic-cup machinery but not a canonical phagosome resident. |
| **Rufy1** | — | ENDOSOME/PHAGOSOME — strong | FYVE-domain Rab4/Rab5 effector involved in PI3P-dependent early-endosomal trafficking; strong fit with early phagosome trafficking/maturation. |
| **Nfkb2** | indirect | — | Major inflammatory transcriptional regulator; cAMP/PKA can influence NF-κB signaling, but NFKB2 is not a convincing canonical direct PKA substrate. |
| **Cnot1** | — | — | CCR4-NOT complex scaffold involved primarily in mRNA turnover and transcriptional regulation. |
| **Faf1** | no established PKA | — | Phosphoprotein involved in apoptosis/NF-κB regulation; phosphorylation by other kinases is better established. |
| **Jun** | indirect | — | AP-1 transcription factor; can be influenced by cAMP/PKA signaling but not a strong direct PKA-substrate hit. |
| **Jund** | indirect | — | AP-1 transcription factor; downstream signaling connection rather than established direct PKA substrate. |
| **Tmod3** | no established PKA | PHAGOCYTIC/CYTOSKELETAL — moderate | Actin pointed-end capping protein controlling cortical actin dynamics; relevant to membrane remodeling but not a phagosome resident. |
| **Tjp3** | — | — | Tight-junction scaffold; primarily associated with epithelial junctions rather than phagosome/lysosome biology. |
| **Cgnl1** | — | CYTOSKELETAL — weak/moderate | Junctional scaffold regulating Rho-family signaling and actomyosin organization; indirect connection to phagocytic cytoskeleton. |
| **Map3k2** | — | — | MAP kinase signaling protein; no strong phagosome/lysosome localization or established PKA-substrate assignment. |
| **Ap1ar** | — | ENDOLYSOSOMAL — strong | Regulates AP-1-dependent TGN/endosome trafficking and Arp2/3; links membrane sorting with actin remodeling. |
| **Bptf** | — | — | Nuclear chromatin-remodeling protein; no compelling phagosome/lysosome connection. |
| **Pphln1** | — | — | Cytoskeletal/nuclear protein with limited evidence for direct phagosome/lysosome involvement. |
| **Gnl3** | — | — | Predominantly nucleolar GTP-binding protein involved in proliferation/ribosome biology. |
| **Arid1b** | — | — | SWI/SNF chromatin-remodeling component; predominantly nuclear. |
| **Des** | — | — | Intermediate-filament protein; not a canonical phagosome/lysosome protein. |
| **Plekhg3** | — | PHAGOCYTIC/CYTOSKELETAL — moderate/strong | RhoGEF with strong actin-remodeling activity; mechanistically well positioned to regulate phagocytic membrane dynamics. |
| **Api5** | — | — | Nuclear anti-apoptotic protein; no strong phagosome/lysosome connection. |
| **Hspb1** | no established PKA | CYTOSKELETAL — moderate | HSP27 regulates actin polymerization/remodeling; highly phosphorylated, but MAPKAPK2/3 rather than PKA is the major established kinase pathway. |
| **Snx16** | — | ENDOSOME/LYSOSOME — strong | PI3P-binding sorting nexin on early/late endosomes; participates in endosomal maturation and endosome-to-lysosome trafficking. |
| **Nup160** | — | — | Nuclear pore complex component; no compelling phagosome/lysosome connection. |
| **Arpc5** | — | PHAGOSOME — strong | Arp2/3 complex component; actin nucleation is central to phagocytic-cup formation and particle engulfment. |
| **Rprd2** | — | — | Primarily nuclear transcription/RNA-processing-associated protein; no strong phagosome/lysosome connection. |

**Table S4**: **Phosphoproteins preferentially elevated in GIV-KO macrophages at 45 min after *AIEC*-LF82 infection**, **related to Figure 1E-F**

| **Gene** | **PKA substrate** | **Phagosome / lysosome association** | **Assessment** |
|---|---|---|---|
| **Rasgrp2** | YES — strong | — | Direct PKA substrate; reported sites include S116/S117, S554/S587 [Reference S10]. |
| **Rab31** | — | PHAGOSOME — strong | Recruited specifically to **newly forming/early phagosomes in macrophages**, then disappears before LAMP1+ phagolysosome maturation. |
| **Slc29a3 / ENT3** | not established | LYSOSOME + PHAGOSOME — very strong | Bona fide lysosomal nucleoside transporter; strikingly, recent work shows recruitment to **phagosomes after bacterial capture** and control of phagosomal acidification/signaling. |
| **Mfsd1** | — | LYSOSOME — very strong | Bona fide **lysosomal membrane transporter**, exporting degradation-derived dipeptides. |
| **Slc2a6 / GLUT6** | — | LYSOSOME — strong | Annotated lysosomal/lysosomal-membrane glucose transporter; particularly relevant in hematopoietic cells. |
| **Cybc1 / EROS** | — | PHAGOCYTE machinery — strong functional link | Required for CYBA/CYBB stability and the **NOX2 respiratory burst after phagocytosis**. Protein itself is primarily ER-localized, so I would call this phagosome-*functional*, not phagosome-localized. |
| **Synj1** | not convincingly established | ENDOSOME/LYSOSOME — moderate | Strong endocytic trafficking connection; loss alters lysosomal morphology/function, but I would not classify it as a canonical resident lysosomal protein. |
| **Man1b1** | — | weak/indirect | Primarily ER/Golgi quality-control protein; can target substrates toward lysosomal degradation, but **not a lysosomal resident protein**. |

**Table S5**: **Temporal organization of GIV-dependent phosphosignaling during *AIEC* LF82 infection, related to Figure 1E-F.**

| Signaling phase | WT / GIV-intact macrophages | GIV-KO macrophages | Biological interpretation |
|---|---|---|---|
| 10 min — Early microbial response | **Signal initiation and small-GTPase/cytoskeletal control:** RAPGEF6, PLEKHG1, STRN, TOMM70; additional early membrane/cytoskeletal regulators identified in the WT-enriched dataset | **Aberrant early engagement of signaling and trafficking machinery:** MYO1F, PIK3AP1, PKN1/2, STK10, LAMTOR1, SLC17A5, AKAP1 (*controls where activated PKA is localized*) | WT cells preferentially engage signaling modules positioned to coordinate small-GTPase activity, actin remodeling, and controlled phagosome formation. In GIV-KO cells, infection instead produces an early phosphosignaling state enriched for kinase-, trafficking-, lysosomal-, and PKA-compartmentalization machinery, consistent with premature or misdirected engagement of downstream effector systems. |
| 45 min — Late microbial response | **Transition toward membrane remodeling and phagosome maturation:** PLEKHG1/3, ARPC5, RUFY1, SNX16, AP1AR, TMOD3 | **Persistent phagolysosomal and PKA/Rap-associated signaling:** RAB31, CYBC1, SLC29A3/ENT3, MFSD1, SLC2A6/GLUT6, RASGRP2† (*direct PKA-regulated Rap1 GEF*), ROCK2, PRKCH | WT macrophages progress toward an organized actin–endosome–phagosome maturation program. GIV-KO macrophages retain a distinct phosphoproteomic state enriched for phagosome/lysosome machinery and cAMP/PKA-linked Rap signaling, consistent with failure to transition normally through the infection response. |
| Dominant temporal trajectory | **Small-GTPase signaling → actin remodeling → endosomal trafficking → phagosome maturation** | **Early dysregulated signaling → persistent/misdirected PKA–Rap and phagolysosomal signaling** | GIV therefore appears to govern **signal-state transition**, rather than globally increasing or decreasing phosphorylation. Its loss reorganizes the timing and composition of the infection-induced phosphoproteome. |
| Connection to subsequent experiments | Coordinated trafficking/maturation signature predicts productive phagosomal acidification and maturation | PKA-linked and phagolysosomal abnormalities predict sustained cAMP–PKA activity and defective maturation | These phosphoproteomic predictions are subsequently tested directly by cAMP/PKA measurements, pharmacologic rescue, lysosomal acidification, pHrodo assays, and ultrastructural analysis. |

This table summarizes biologically informative temporal patterns emerging from the complete genotype-resolved phosphoproteomic datasets in **Tables S1–S4**. Protein-level values represent phosphorylation-associated abundance and should not be interpreted as measurements of individual phosphosites. Functional annotations indicate established or plausible relationships to cAMP/PKA signaling, small-GTPase regulation, cytoskeletal remodeling, or phagosome/lysosome biology. Because the 10- and 45-min genotype comparisons contain two biological samples per genotype, these analyses are considered exploratory and hypothesis-generating.

†Established direct PKA substrate.

**Table S6**: **Raw EDM-TEER values (Ω·cm²) following *AIEC* LF82 infection,** **related to Figure 7F-H**

| | WT + DMSO | WT + IGGi-11-me | WT + IGGi-11-me + H89 | KO + DMSO |
|---|---|---|---|---|
| **Pre-infection** | 349 | 258 | 314.33 | 268 |
| | 359 | 262.33 | 342.66 | 226 |
| | 234 | 235 | 219.33 | 233 |
| | 219.69 | 213.66 | 217.33 | 194.66 |
| | 229.66 | | 230.66 | |
| **6 hours** | 2428.33 | 1887.66 | 3242 | 2176.66 |
| | 2904.32 | 2006 | 3184 | 1562.33 |
| | 2238 | 1944.66 | 2610.33 | 1598.66 |
| | 2426 | 1399.33 | 2304.66 | 792 |
| | 1866 | | 2146.66 | |
| **24 hours** | 1028.66 | 2176.66 | 2332 | 932.66 |
| | 842.65 | 1562.37 | 1706.66 | 618.33 |
| | 1346 | 901 | 1093.33 | 1076.66 |
| | 698.61 | 1119.66 | 1040.66 | 590.66 |
| | 841.62 | | 1198.33 | |
| **36 hours** | 893.33 | 945 | 1823 | 708.33 |
| | 614.33 | 704 | 1426 | 644.33 |
| | 1140 | 692.33 | 907 | 387.33 |
| | 444.66 | 1011.66 | 826.66 | 362.33 |
| | 602.67 | | 839.33 | |
